# NFAT5 induction by the tumor microenvironment enforces CD8 T cell exhaustion

**DOI:** 10.1101/2022.03.15.484422

**Authors:** Laure Tillé, Daniela Cropp, Gabrielle Bodley, Massimo Andreatta, Mélanie Charmoy, Isaac Crespo, Sina Nassiri, Joao Lourenco, Marine Leblond, Cristina Lopez-Rodriguez, Daniel E. Speiser, Santiago J. Carmona, Werner Held, Grégory Verdeil

**Author notes:** Co-first authors (Laure Tillé, Daniela Cropp). Correspondence:* Grégory Verdeil, Department of Oncology, University of Lausanne, Biopole 3, Ch. des Boveresses 155, CH-1066 Epalinges, Switzerland. Author contributions:* L.T, D.C, WH and G.V conceived and designed the experiments. L.T, D.C, M.C, GB, M.L and G.V performed the experiments. J.L analyzed the scRNA-seq data and M.A, S.C and I.C performed bioinformatic analysis. C.L.R provided CD4-Cre NFAT5^fl/fl^ mice. L.T, D.C and G.V prepared the figures and wrote the manuscript with input from all authors. D.E.S, W.H, C.L.R, S.C significantly reviewed the manuscript.

## Abstract

Persistent exposure to antigen during chronic infection or cancer renders T cells dysfunctional. The molecular mechanisms regulating this state of exhaustion are thought to be common in infection and cancer, despite obvious differences in their microenvironments. We discovered that NFAT5, an NFAT family member lacking an AP-1 docking site, is highly expressed in exhausted T cells from murine and human tumors and is a central player in tumor-induced exhaustion. While NFAT5 overexpression reduced tumor control, NFAT5 deletion improved tumor control by promoting the accumulation of tumor-specific CD8+ T cells that expressed less TOX and PD-1 and produced more cytokines particularly among precursor exhausted cells. Conversely, NFAT5 had no effect on chronic infection-induced T cell exhaustion. Mechanistically we found that TCR triggering induced NFAT5 expression and that hyperosmolarity stimulated transcriptional activity of NFAT5. We propose that NFAT5 takes over NFAT1/2 to promote exhaustion specifically in tumor-infiltrating CD8+ T cells.

## Introduction

CD8 T cells can actively recognize and eliminate tumor cells. However, CD8 tumor-infiltrating lymphocytes (TILs) are mostly dysfunctional, commonly referred to as exhausted. Exhausted CD8 T cells responding to chronic infection and cancer show high expression of multiple inhibitory receptors, reduced effector functions and are not able to efficiently control pathogens or tumors^1^. The exhaustion state is strongly related to the constant presence of antigen, resulting in continuous triggering of the TCR^2^, but the composition of the local microenvironment further influences the gene expression of exhausted CD8 T cells ^3^.

Despite the tremendous progress in cancer immunotherapy during the past years^4^, a large fraction of patients’ cancers remains, or becomes, therapy resistant. Understanding the molecular mechanisms that regulate T cell exhaustion is a first step towards more efficient treatments. Recent studies highlighted the existence of a precursor exhausted T cell (Tpex) population with stem cell-like properties, which can further differentiate into terminally exhausted T cells (Tex) that have cytolytic potential but are short lived. Several transcription factors (TF) such as TOX and NFAT play central roles in the establishment of T cell exhaustion, while others, such as TCF-1, maintain the stemness properties of Tpex ^1, 5, 6^. TOX drives the expression of inhibitory receptors and negatively regulates the production of inflammatory cytokines allowing T cell maintenance in the context of chronic antigen stimulation^7, 8^. Importantly, it has been shown that TOX is directly regulated by NFAT1 and NFAT2^9, 10^, which are activated by calcineurin downstream of TCR signaling^11^. NFAT1 and NFAT2 are required for effective CD8 T cell differentiation into cytotoxic T cells by forming dimers with transcriptional partners such as AP-1^12^. However, the overexpression of a constitutively active version of NFAT1 unable to interact with AP-1 induces an exhausted phenotype in CD8 TILs^13^. As AP-1 expression in chronically-stimulated T cells is reduced, exhaustion is at least partly induced by NFAT activation in the relative absence of AP-1^13^.

A previous transcriptomic analysis of CD8 T cells from tumor-infiltrated lymph nodes (TILN) showed that the NFAT family member NFAT5 is highly expressed in TILN cells^14^. In contrast to the classical NFAT proteins, NFAT5 lacks an AP-1 docking site and is not regulated by calcineurin^15^. Instead, NFAT5 is triggered by metabolic stress, such as hypertonicity, and regulates the transcription of proteins involved in the maintenance of an adequate osmotic balance in a cell type-unspecific manner^16^. However, the activity of NFAT5 varies according to the cell type or the stimulus^17, 18^. Recent studies found that NFAT5 regulates inflammatory responses in macrophages^19^ and CD4 T cells^20^, but so far there is no report on functional alterations of peripheral CD8 T cells through NFAT5^17^.

Our study showed that NFAT5 is highly expressed in CD8 TILs from murine and human tumors. Overexpression of NFAT5 dampened CD8 T cell responses against tumor cells, while deletion of NFAT5 further improved CD8 T cell anti-tumor functions without impacting their capacity to accumulate and differentiate. Surprisingly, NFAT5 deletion in CD8 T cells during chronic LCMV infection had no effect on T cell exhaustion and virus control, emphasizing a tumor-specific T cell regulatory role of NFAT5. By deciphering the different stimuli present in the tumor microenvironment (TME), we found that TCR triggering is the main inducer of NFAT5 expression and that hyperosmolarity increases NFAT5 activity in the TME. Therefore, our data established that NFAT5 is a tumor-specific regulator of CD8 T cell exhaustion.

## Results

### NFAT5 is upregulated in tumor-infiltrating CD8 T cells

NFAT5 was previously found highly expressed in Melan-A-specific CD8 T cells obtained from metastasized lymph nodes^14^. To confirm the expression of *NFAT5* in CD8 TILs, we took advantage of publicly available single cell RNA-seq (scRNA-seq) data from mouse B16 melanoma^21^, mouse MC38 adenocarcinoma^22^, human melanoma^23, 24^ and human breast cancer^25^. The CD8 TIL were classified into naïve like, early activated, effector memory, precursor exhausted (Tpex) and terminal exhausted (Tex) CD8 T cell subsets, using ProjecTILs^26^. *NFAT5* was highly expressed in Tpex and Tex compared to naïve like, early effector and effector memory CD8 TILs in all studies (Fig. 1a, Extended Data Fig. 3). We used the same datasets to identify the most relevant TFs regulating T cell exhaustion by comparing their regulon activity (AUC score) in the different subsets. A regulon represents a gene set regulated by the same transcription factor. The regulon activity of NFAT5 was upregulated in Tpex and Tex. NFAT5 is in the top 8 TFs showing statistically significant differences between these two subpopulations compared to the other subtypes, together with Tbet, Runx2 and Bhlhe40 for upregulated regulons (Fig. 1b). We further confirmed the upregulation of NFAT5 by quantitative PCR (qPCR) in CD8 T cells sorted from the spleen or tumors of B16 melanoma^27^ tumor-bearing mice (Fig. 1c) and in human Melan-A-specific CD8 TILN, using circulating EBV-specific or naïve circulating CD8 T cells as controls (Fig. 1d).

**Fig. 1:**
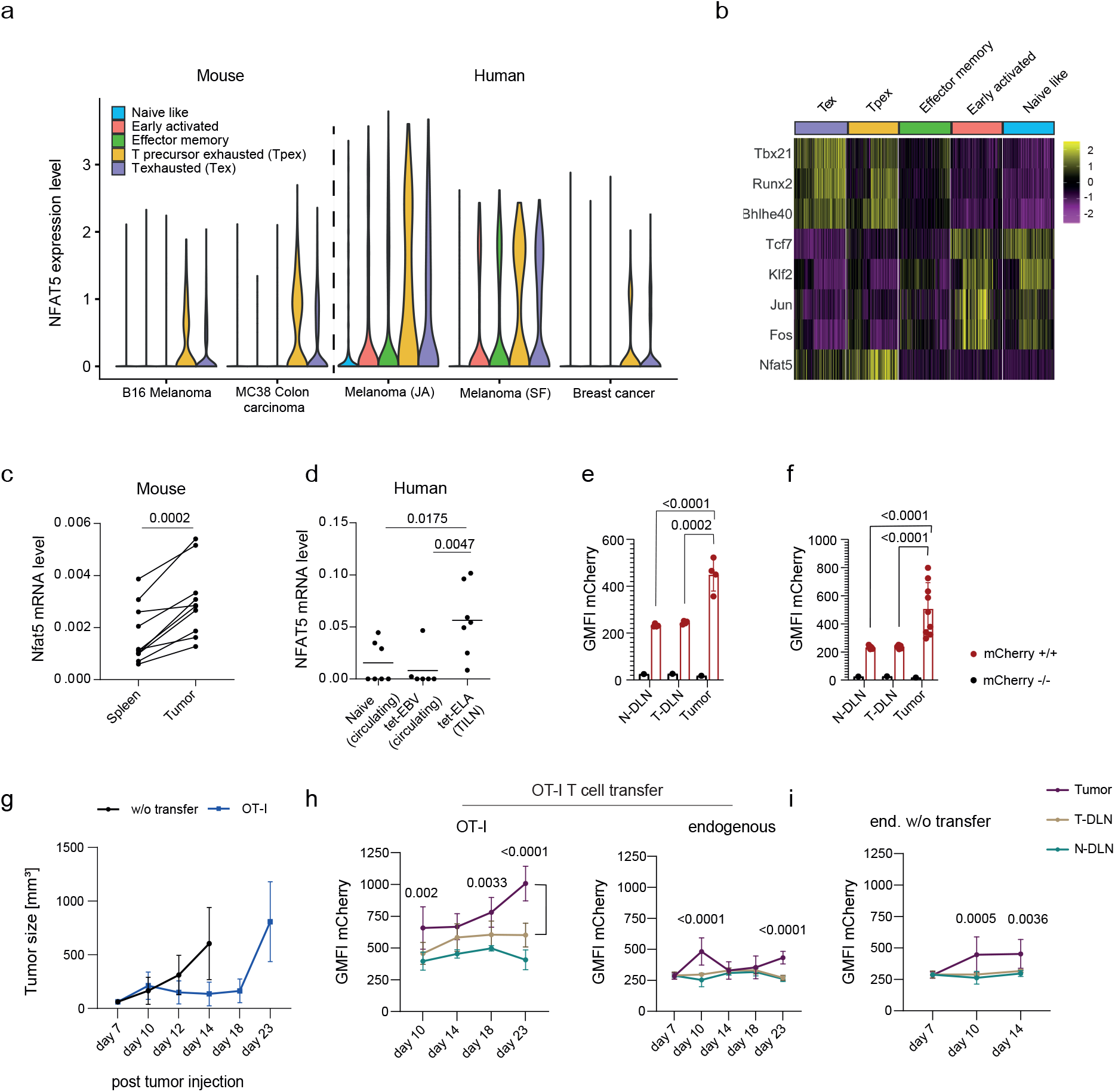
NFAT5 is upregulated in tumor-infiltrating CD8 T cells. **a)** *NFAT5* mRNA expression levels in each indicated CD8 TIL subtype, as classified by ProjecTILs, across five different mouse and patient cohorts/studies (CF Material and methods). **b)** Heatmap showing the activity (AUC score) of the top 8 TFs with the greatest difference in regulon activity (either up or down regulation) when comparing terminal exhausted CD8 T cells (Tex) and naïve-like CD8 T cells (Naive like) from tumor-infiltrating T lymphocytes (TILs dataset). **c)** *Nfat5* mRNA levels from CD8 T cells homing B16-gp33 tumors or spleens. Ten mice pooled from two independent experiments. **d)** *NFAT5* mRNA level in human naïve circulating CD8 T cells, EBV-specific CD8 T cells and ELA (Melan-A)-specific CD8 T cells from tumor-infiltrated lymph nodes (TILN). **e)** mCherry levels in non-draining lymph node (N-DLN), tumor-draining lymph node (T-DLN) and B16-gp33 tumors from NFAT5_mCherry+/+_ or NFAT5_mCherry-/-_ CD8 T cells on day 16 post tumor injection. Four mice from one representative experiment out of two. **f)** mCherry levels in N-DLN, T-DLN and MC38 tumors from NFAT5_mCherry+/+_ or NFAT5_mCherry-/-_ CD8 T cells on day 16 post tumor injection. Nine mice pooled from two independent experiments. **g)** Tumor growth measured in NFAT5_mCherry+/+_ mice transferred with OT-I-NFAT5_mCherry+/+_ CD8 T cells (day 7 post tumor injection) or no T cell transfer (w/o transfer). **h)** mCherry expression of OT-I-NFAT5_mCherry+/+_ (left) or endogenous-NFAT5_mCherry+/+_ (right) CD8 T cells from the N-DLN, T-DLN and tumor. Statistical comparison between CD8 T cells from the T-DLN and tumor. **i)** mCherry expression of NFAT5_mCherry+/+_ mice not receiving T cell transfer, TILs, T-DLN and N-DLN CD8 T cells. **g-i)** Two pooled independent experiments with 9 mice per condition. **c)** Paired student t-test. **d)** Mann-Whitney test. Mean. **e-i)** Two-way ANOVA. Mean with SD

To follow NFAT5 expression at the single cell level, we generated an NFAT5 reporter mouse strain, in which the stop codon in exon 14 of NFAT5 was replaced by a P2A-mCherry cassette (Extended Data Fig. 1a). In these mice, mCherry expression in CD8 T cells correlated with the level of NFAT5 mRNA (Extended Data Fig. 1b). We further confirmed that the introduction of the P2A-mCherry cassette did not alter NFAT5 expression. We compared the NFAT5 protein level in CD8 T cells from WT, NFAT5 KO and NFAT5_mCherry_ mice (Extended Data Fig. 1c, d). We did not observe any effect on the viability or breeding capacity, or thymic development, of the NFAT5-mCherry reporter mouse strain (Extended Data Fig. 1e-g). Using this model, we showed that polyclonal CD8 TILs from B16 or MC38 tumors expressed significantly higher levels of NFAT5 compared to CD8 T cells from the tumor-draining lymph node (T-DLN) or the non-draining lymph node (N-DLN) on day 16 post tumor injection (Fig. 1e-f). To define the kinetic of NFAT5 induction in CD8 TILs, we engrafted NFAT5_mCherry_ mice with B16-OVA (Fig. 1g). Seven days post engraftment, endogenous CD8 T cells from the tumor and the LNs expressed similar levels of mCherry. At this time point we transferred activated OT-I-NFAT5_mCherry_ CD8 T cells into some of the tumor bearing mice. Three days after transfer, both endogenous and OT-I CD8 TILs showed an increased level of mCherry, which remained stable for OT-I cells but dropped for endogenous CD8 TILs, while tumor growth was transiently controlled, until day 18 (Fig. 1h). Once tumor growth resumed, we observed a strong increase of mCherry levels in OT-I cells and to a lesser extent in endogenous CD8 TILs. In the absence of OT-I CD8 T cell transfer, B16-OVA tumor growth was not controlled. In this situation, the level of mCherry also increased at day 10 and remained stable at day 14 (Fig. 1i). The rapid growth of the tumors did not allow us to measure further time points. Altogether, we found enhanced NFAT5 expression in Tpex and Tex CD8 TILs both in human and murine tumors and increasing NFAT5 levels during tumor progression.

### NFAT5 overexpression dampens CD8 T cell tumor control

To test whether high NFAT5 levels in CD8 T cells impact their response against established tumors, we cloned different NFAT5 isoforms that differ in the alternative splicing of the first and last exons, into GFP-expressing retroviral vectors (Fig. S2a) and transduced TCRP1A-luc+ CD8 T cells expressing luciferase and a TCR recognizing the P1A epitope expressed by P511 mastocytoma cells^28, 29^. Adoptive transfer of as little as 10^4^ transduced TCRP1A-luc+ CD8 T cells was sufficient to induce tumor regression^30, 31^. Following the transfer of sorted NFAT5 isoform A, that lacks the Nucleor Export Signal (NES) sequence, overexpressing CD8 T cells into P511 mastocytoma-bearing Rag1^-/-^B10D2 mice (Fig. 2a-b), we observed reduced tumor control compared to control eGFP-transduced CD8 T cells (Fig. 2c). A similar reduction in tumor control was obtained with NFAT5 isoform D, that contains the NES, while the deletion of the DNA binding domain of NFAT5 restored tumor control (Extended Data Fig. 2). Furthermore, overexpression of NFAT1 CA-RIT, a constitutively active form of NFAT1 unable to bind AP-1, reduced tumor control in a similar extent as NFAT5 (Fig. 2c). In this model we have previously shown a disadvantage of CD8 T cells overexpressing exhaustion-associated genes to sustain in the host, leading to a rapid enrichment of GFP-TCRP1A T cells that do not express our gene of interest^30, 31^. Therefore, even the slight, but significant, delay in tumor control was a good indicator of NFAT5-mediated impairment of CD8 TIL anti-tumor functions. We did not detect any difference in T cell accumulation by measuring bioluminescence throughout the experiment, suggesting that NFAT5 did not impair the infiltration of CD8 T cells into the tumor (Extended Data Fig. 2). To further characterize their phenotype, we sorted eGFP+ CD8 TILs seven days after T cell transfer and performed RNA-seq analysis. Principal component (PC) analysis revealed that NFAT5 and NFAT1 CA-RIT-overexpressing CD8 TILs clustered together, distant from control eGFP-transduced CD8 T cells (Fig. 2d). Most of the differentially expressed genes (n=35), compared to control eGFP CD8 TILs, were upregulated in both NFAT5 and NFAT1 CA-RIT-overexpressing CD8 TILs (Fig. 2e). Within the shared genes we found Dusp family phosphatases (2, 5 and 10) as well as *Nr4a1* and *Nr4a3*, which have been associated with the regulation of T cell exhaustion^32^. Gene Ontology (GO) term analysis revealed similar transcriptomes of both NFAT5 and NFAT1 CA-RIT-overexpressing CD8 TILs, namely signatures associated with cell cycling (chromosome condensation, mitotic regulation), inhibition of MAPK signaling and cell differentiation. Altogether, NFAT5 overexpression in tumor-specific CD8 T cells reduced tumor control through the induction of a transcriptional program similar to that induced by a NFAT1 construct that cannot associate with AP-1.

**Fig. 2:**
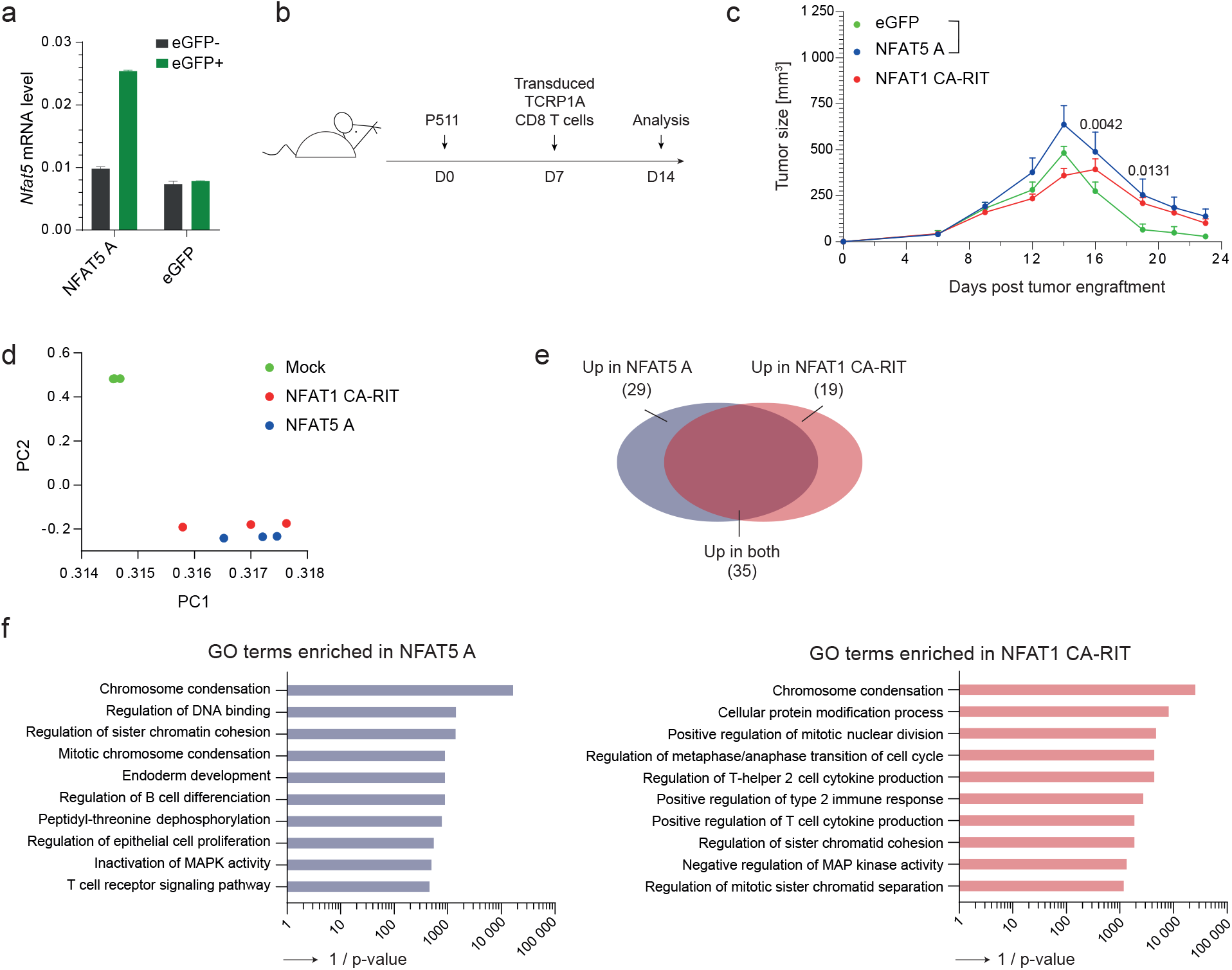
NFAT5 overexpression dampens CD8 T cell tumor control. **a)** *Nfat5* mRNA levels in CD8 T cells transduced with a vector encoding for NFAT5 isoform A (NFAT5 A) (left) or control eGFP (right). **b)** Timeline of the experiment. Activated TCRP1A CD8 T cells were transduced and transferred into Rag1^-/-^B10D2 mice seven days post P511 tumor engraftment. CD8 T cells were sorted for RNA-sequencing on day 14. **c)** Tumor growth in mice transferred with TCRP1A CD8 T cells transduced with control eGFP, NFAT5 A, or NFAT1 CA-RIT. **d)** PC analysis of CD8 TILs transduced with control eGFP, NFAT5 A, or NFAT1 CA-RIT. **e)** Venn diagram showing the number of genes upregulated in NFAT5 A transduced CD8 TILs, NFAT1 CA-RIT transduced TILs, or both. **f)** GO terms enriched in CD8 TILs with NFAT5 A (left), or NFAT1 CA-RIT (right). **c)** Two way ANOVA. Error bars represent SEM. One representative experiment out of two with 5 mice per group.

### NFAT5 deletion in tumor-specific T cells improves tumor control

To test whether NFAT5 deletion influenced the tumor response, we crossed CD4-Cre NFAT5^flox/flox^ mice with a P14 TCR transgenic strain, which is specific for the LCMV-derived gp33 epitope. In this model, T cell-specific deletion of NFAT5 does not impair T cell development or alter the peripheral T cell compartment^33^. We transferred activated NFAT5^flox/flox^ CD4-Cre-/- (WT) or NFAT5^flox/flox^ CD4-Cre+/- (KO) P14 CD8 T cells into mice bearing subcutaneous gp33-expressing B16 melanoma (B16-gp33) (Fig. 3a). Mice transferred with KO P14 CD8 T cells developed much smaller (and later) tumors compared to the transfer of WT P14 CD8 T cells (Fig. 3b). Seven days after T cell transfer, higher proportions of KO than WT P14 CD8 T cells infiltrated the tumor. Strikingly, KO CD8 TILs produced more IFN-γ, TNF-α and IL-2 upon *ex-vivo* stimulation and expressed less PD-1 than WT P14 CD8 TILs, while CD44 expression was comparable (Fig. 3c), indicating reduced exhaustion of NFAT5 KO CD8 T cells.

**Fig. 3:**
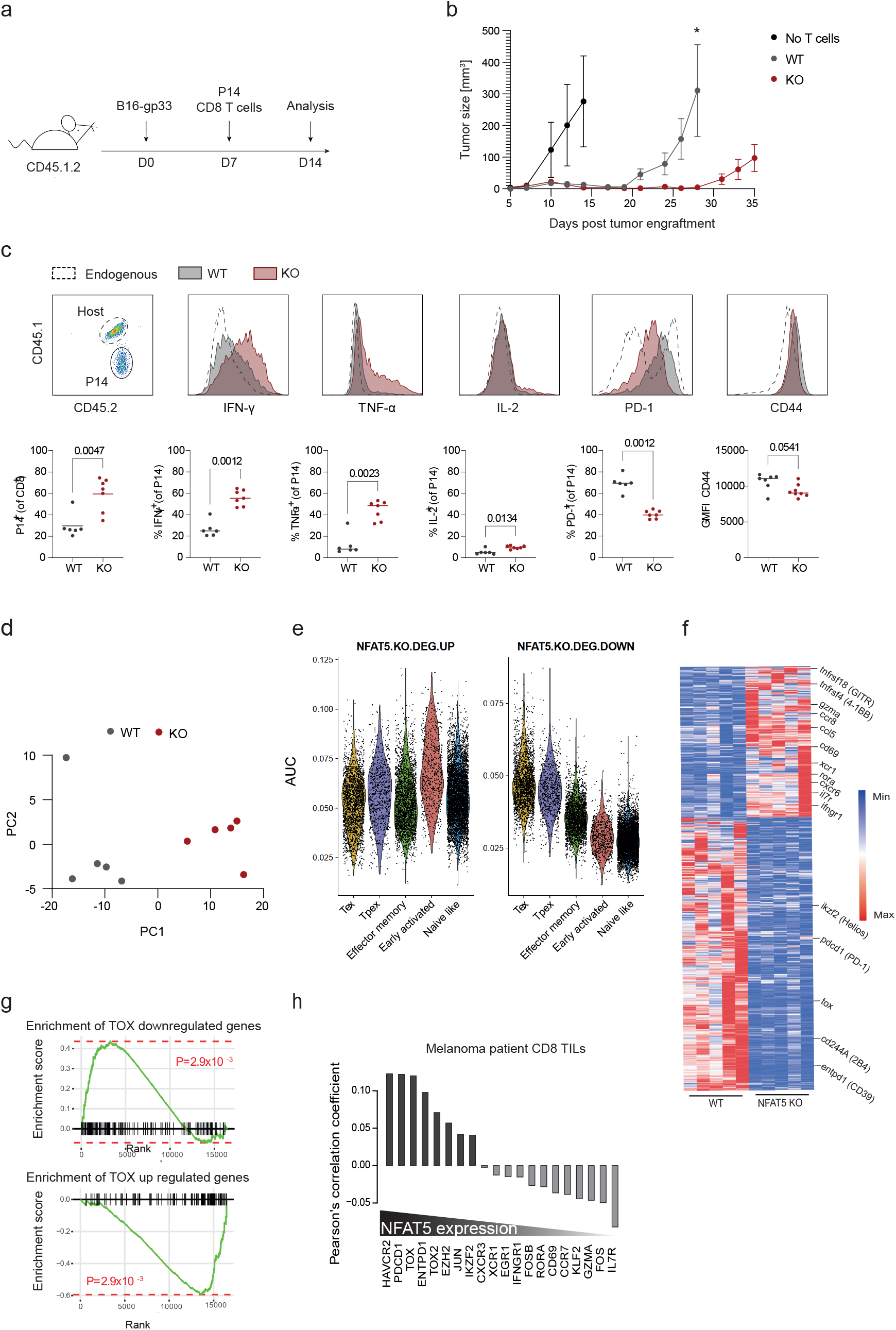
NFAT5 deletion in tumor-specific T cells improves tumor control. **a)** Timeline of the experiment: activated WT or NFAT5 KO P14 CD8 T cells were transferred into B16-gp33-bearing CD45.1.2 mice. CD8 TILs were analyzed seven days after transfer. **b)** Tumor growth of mice transferred with WT (grey) or NFAT5 KO (red) P14 CD8 T cells. In black is the control group without T cell transfer. **c)** WT or NFAT5 KO P14 CD8 TILs were analyzed by flow cytometry seven days after transfer. Bars represent the geometric mean. **d)** PC analysis of WT and NFAT5 KO P14 CD8 TILs. **e)** Violin plots showing the distribution of upregulated (left) or downregulated (right) genes among indicated tumor-infiltrating CD8 T cell subpopulations, including precursor exhausted (Tpex) and terminal exhausted (Tex). **f)** Heatmap displaying 1294 genes differentially expressed in WT versus NFAT5 KO P14 CD8 TILs, fold change of 1.5; adjusted p value <0.05). **g)** Genes differentially expressed in TOX KO CD8 T cells were compared to genes deferentially expressed in NFAT5 KO CD8 T cells using the GSEA analysis. **h)** Analysis of scRNA-seq data of human melanoma tumors: Pearson correlation coefficients were calculated between *NFAT5* mRNA levels and mRNA levels of indicated genes. **a-c)** Seven mice per condition from one representative experiment out of three. Mann-Whitney test.

To decipher the molecular mechanism for the enhanced tumor control by KO CD8 T cells, we performed RNA sequencing on sorted P14 CD8 TILs seven days after transfer. PC analysis revealed that WT and KO CD8 TILs clustered separately (Fig. 3d). We found 458 genes significantly upregulated and 833 genes significantly downregulated in KO CD8 TILs (Extended Data Table 1). By comparing the gene signatures to available scRNA-seq data from CD8 TILs^21, 22, 23, 24, 25^, we found that KO CD8 TILs overexpressed genes found in early activated CD8 TILs, while genes highly expressed in Tex and Tpex were downregulated in KO CD8 TILs (Fig 3e). Indeed, several genes previously associated with CD8 T cell exhaustion were downregulated in KO cells, including *Cd244a* (2B4), *Entpd1* (CD39), *Pdcd1* (PD-1), *Ikzf2* (Helios) and *Tox* (Extended Data Table 1). Conversely, KO cells overexpressed the cytotoxic molecule Granzyme A, activation-associated genes such as *Tnfrsf4* (4-1BB), *Tnfsfr18* (GITR) and *Ccl5*, genes expressed by T resident memory cells (*Cd69, Cxcr6, Ccr8*), memory T cells (*Il7r*) or associated with T cell differentiation (*Rora*) (Fig. 3f, Extended Data Table 1). Given the role of TOX in T cell exhaustion^7, 8, 9, 34^, we assessed whether a reduced TOX expression accounted for the effect of NFAT5 KO. We compared our signature with the one obtained after *Tox* inactivation using GSEA^34^. We found that genes downregulated upon NFAT5 KO were significantly enriched among TOX-dependent genes and *vice versa* (Fig. 3g). To extend our findings to human T cells, we studied the correlation of NFAT5 expression with exhaustion-related genes in human melanoma TILs, using single cell RNA-seq^35^. We calculated the Pearson correlation coefficients between *NFAT5* and all detected genes in activated CD8 TILs as judged by *CD44, PDCD1* or *TNFRSF4* expression. Consistently, we found that *NFAT5* expression in human TILs positively correlated with *HAVCR2, PDCD1* and *TOX* expression. Conversely, *NFAT5* expression negatively correlated with *IL7R, GZMA* and *CD69* expression (Fig. 3h). Altogether, NFAT5 KO CD8 TILs expressed less PD-1 and TOX and produced more inflammatory cytokines, resulting in a more efficient tumor control. Furthermore, we found that in both murine and human datasets, NFAT5 plays a major role in the regulation of exhaustion-associated genes.

### NFAT5 does not alter the CD8 T cell response to chronic LCMV infection

T cell exhaustion also develops during chronic infection^36^. We therefore tested whether NFAT5 regulated the CD8 T cell response to chronic viral infection. We adoptively transferred mice with naïve WT or KO P14 cells one day prior LCMV clone 13 infection, which causes chronic infection (Fig. 4a). At day 28 post-infection, we found slightly higher proportions of KO P14 cells. Compared to WT P14 cells, these cells produced comparable levels of effector cytokines (IFN-γ, TNF-α and IL-2) (Fig. 4c) and only slightly overexpressed PD-1 and CD44. Finally, weight loss during infection, which is indicative of immunopathology, was comparable between mice receiving WT and KO P14 cells (Fig. 4b). Altogether, NFAT5 deficiency did not have a significant effect on the CD8 T cell response to chronic LCMV infection, suggesting a tumor-specific role of NFAT5 in T cell exhaustion.

**Fig. 4:**
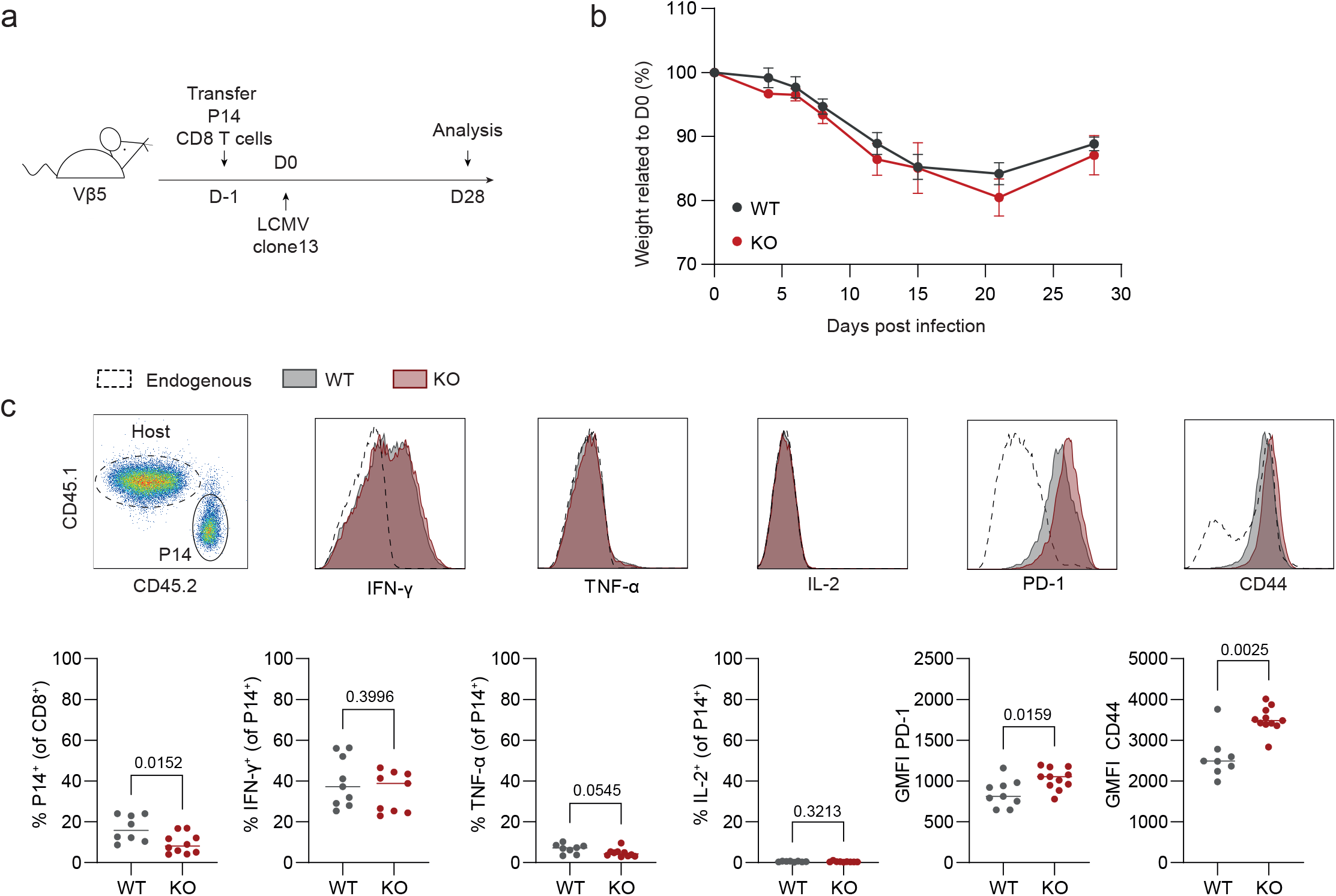
NFAT5 inactivation has no effect on CD8 T cell activity during chronic LCMV infection. **a)** Timeline of the experiment. WT or NFAT5 KO P14 CD8 T cells were transferred into Vβ5 mice one day prior LCMV clone 13 infection. **b)** The body weight of infected mice was monitored over time and normalized to day 0. **c)** Flow cytometry analysis was performed 28 days after infection. Two pooled experiments. Mann-Whitney test.

### NFAT5 is preferentially expressed in Tpex within CD8 TILs

To understand why NFAT5 inactivation did not restore CD8 T cell function during chronic infection while CD8 TILs function was strongly improved, we performed a more detailed analysis of NFAT5 expression in exhausted CD8 T cells. We used available scRNA-seq data from murine CD8+ T cells responding LCMV clone 13 infection. Focusing on the various CD8 T cell subsets, we found that NFAT5 levels were high in Tpex compared to Tex and to the other subtypes. Overall, NFAT5 levels were lower in P14 cells responding to chronic infection than P14 TIL subsets (Extended Data Figure 3b, Fig. 1a). To confirm these data, we took advantage of our NFAT5_mCherry_ reporter mouse strain crossed to the P14 TCR transgenic mice (P14-NFAT5_mCherry_). P14-NFAT5_mCherry_ cells were either transferred into B16-gp33 melanoma-bearing mice or into WT mice one day prior infection with LCMV clone 13 (chronic) or Armstrong (acute) strains. After seven days in B16-gp33 tumors and 8 or 28 days in LCMV infected mice, we measured the mCherry levels in Tpex and Tex P14 CD8 T cells (Fig. 5a-b). To compare the two models, mCherry expression was normalized to the fluorescence of endogenous CD8 T cells. NFAT5 expression was higher in Tpex than in Tex after chronic infection at day 28, while the level was low in both populations after acute infection and at day 8 after chronic infection. Strikingly, at day 14 (seven days post-transfer) the fold change of NFAT5 in Tpex from CD8 TILs was higher than in the chronic infection on day 28 (Fig. 5a, b). We further confirmed these data by using OT-I-NFAT5_mCherry_ CD8 T cells transferred into B16-OVA-bearing mice (Fig. 5c). We followed mCherry expression in Tpex and Tex from CD8 TILs on day 10, 14, 16 and 23 post-tumor injection (corresponding to day 3, 7, 11 and 16 post-OT-I transfer). Initially, the levels in Tpex and Tex were similar at day 10 but preferentially increased in Tpex on day 14 and 18, reaching a maximum on day 23. In conclusion, we established that NFAT5 expression levels are lower in CD8 T cells in the context of acute and chronic infection as compared to CD8 TILs and that NFAT5 levels are higher in Tpex compared to Tex, with an increasing level as tumor progresses.

**Fig. 5:**
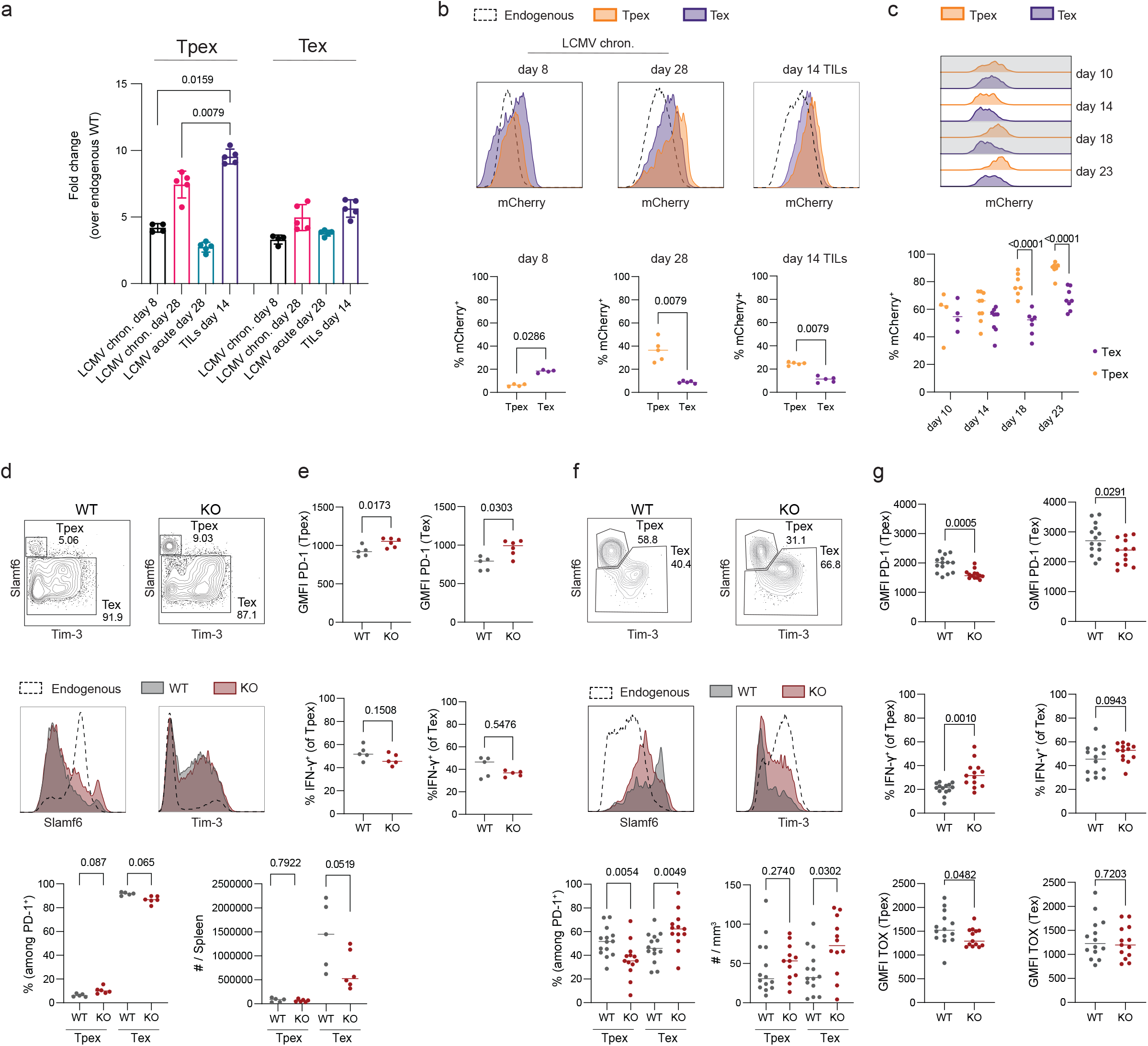
NFAT5 inactivation impacts more strongly Tpex CD8 TILs. **a)** Fold change of P14-NFAT5_mCherry_ cells over endogenous WT CD8 T cells at indicated conditions and time points for Tpex (left) and Tex (right). **b)** Representative histograms of mCherry expression by P14-NFAT5_mCherry_ Tpex (orange) and Tex (purple) compared to endogenous WT (dotted line) CD8 T cells at given time points post LCMV clone 13 infection (left) or tumor injection (right) with quantification of mCherry+ Tpex and Tex at respective time points (below). One representative experiment out of two. **c)** Representative histograms of mCherry expression by OT1-NFAT5_mCherry_ Tpex and Tex at different time points post-tumor injection (up) with quantification of mCherry+ Tpex and Tex at respective time points (below). Two independent experiments pooled. **d)** Contour plots of Slamf6 and Tim-3 expression in WT (left) and NFAT5 KO (right) with the respective histograms comparing to endogenous CD8 T cells on day 28 post LCMV clone 13 infection. Frequencies and numbers per spleen of Tpex and Tex in NFAT5 KO and WT recipients are shown below. **e)** Immunophenotyping of Tpex and Tex by their PD-1 expression and IFN-γ production. **d-e)** One representative experiment out of three. **f)** Contour plots of Slamf6 and Tim-3 expression in WT (left) and NFAT5 KO (right) with the respective histograms comparing to endogenous CD8 T cells on day 14 post tumor injection (day 7 post T cell transfer). Frequencies and numbers per mm^3^ (tumor mass) of Tpex and Tex in NFAT5 KO and WT recipients (below). **g)** Immunophenotyping of Tpex and Tex by their PD-1 expression and IFN-γ production. Two independent experiment pooled out of three. **a**,**b**,**d-g)** Mann-Whitney test. **c)** Two ways ANOVA. Bars representing geometric mean.

### NFAT5 inactivation impacts strongly Tpex CD8 TILs

Considering the differential NFAT5 expression in Tpex and Tex, we looked at the effects of NFAT5 KO on these populations. After chronic infection, we did not observe significant changes neither in the proportions of the two populations (Fig. 5d) nor in absolute numbers. IFN-γ production, as well as the level of PD-1, were also similar (Fig. 5e). In contrast, NFAT5 KO resulted in a decreased proportion, but similar number of Tpex CD8 TILs compared to WT CD8 TILs, while both the proportion and number of Tex increased seven days after T cell transfer in-tumor bearing mice (Fig. 5f). Furthermore, NFAT5 KO Tpex expressed lower levels of PD-1 and TOX and included higher frequencies of IFN-γ-producing cells as compared to WT CD8 TILs, while Tex, although in greater number, were less impacted by the NFAT5 deletion (Fig. 5g). We established that NFAT5 KO led to an increased number of tumor-specific CD8 T cells. This effect was mostly mediated through Tpex, which displayed a less exhausted phenotype and differentiated more efficiently into cytotoxic Tex, resulting in improved tumor control.

### NFAT5 expression is driven by TCR signaling in CD8 TILs

To explore how NFAT5 is triggered in CD8 TILs, we tested various stimuli known to be associated with the TME or described to regulate NFAT5 expression and/or activity. To test the effect on NFAT5 expression in CD8 T cells, we cultured P14-NFAT5_mCherry_ splenocytes for 72h in hyperosmotic and/or hypoxic conditions with or without TCR stimulation. TCR stimulation with gp33 peptide or anti-CD3 and anti-CD28 antibodies drastically increased mCherry levels. Separately, the addition of NaCl or KCl to the cell culture medium lead to a dose-dependent (380mOsm vs 420mOsm) increase of mCherry levels. Strikingly, combining TCR triggering and hypertonic medium (with NaCl or KCl) led to maximal mCherry expression. On the other hand, hypoxic conditions (0,5% O_2_) did not induce mCherry expression alone or in combination with TCR triggering or high osmolarity (Fig. 6a).

**Fig. 6:**
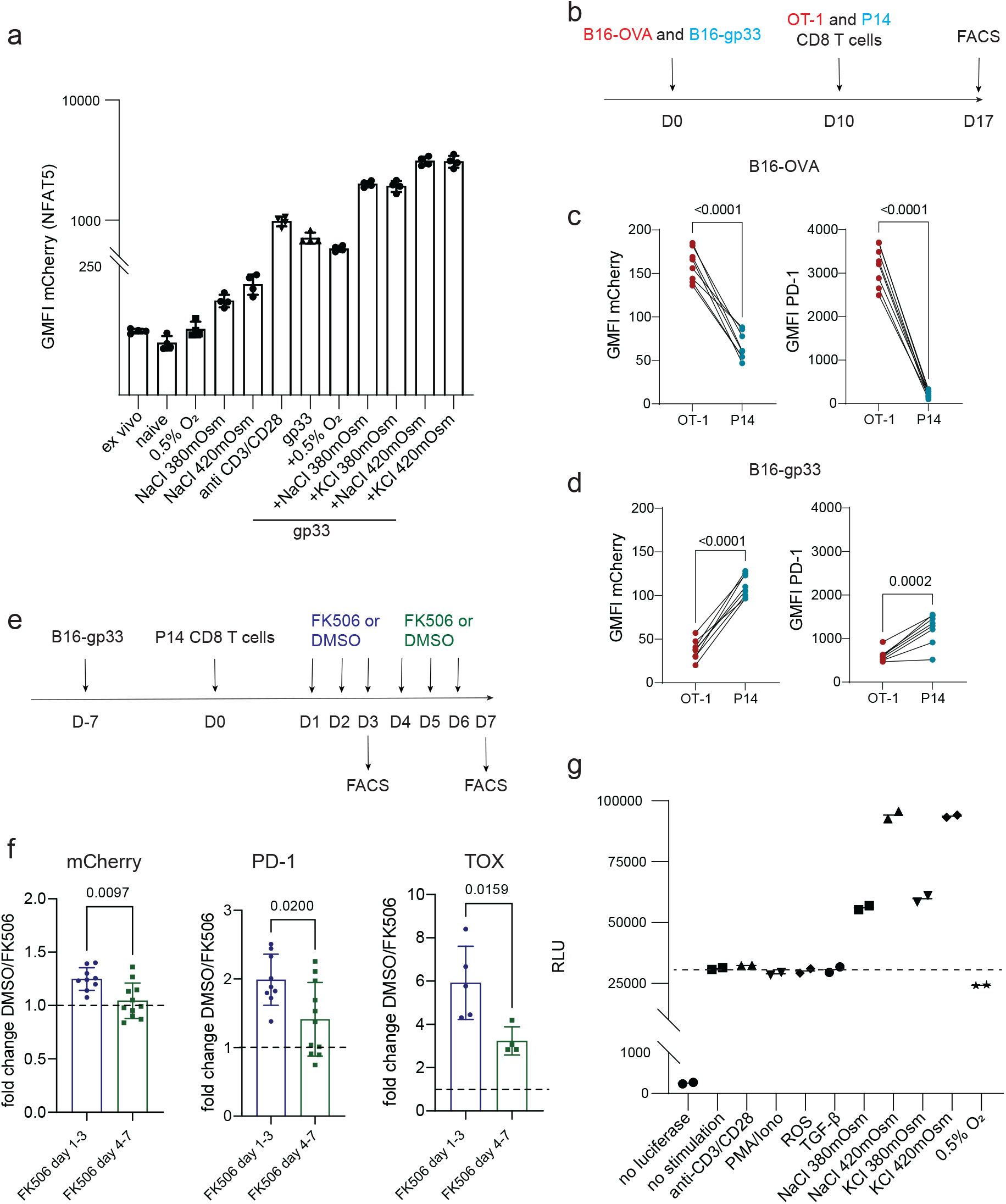
NFAT5 expression is driven by TCR signaling in CD8 TILs. **a)** mCherry levels in NFAT5_mCherry_ splenocytes *ex-vivo*, after three days in culture in the presence of IL-2 (naïve) or in combination with the indicated stimuli. **b)** Timeline of the experiment. Activated OT-I and P14 CD8 T cells were transferred into B16-OVA and B16-gp33-bearing mice. Flow cytometry analysis seven days after T cell transfer (day 16). **c)** Paired comparison of mCherry and PD-1 expression plotted as GMFI of indicated TILs within B16-OVA tumors. **d)** Paired comparison of mCherry expression plotted as GMFI of indicated TILs within B16-gp33 tumors. **e)** Timeline of the experiment. B16-gp33-bearing mice were treated with FK506 either the first three days after T cell transfer (7 days post tumor injection) or 4-6 days after T cell transfer. TILs were analyzed on day 3 and day 7 post T cell transfer. **f)** Fold change of mCherry (left), PD-1 (middle) and TOX (right) between DMSO and FK506-receiving mice at respective time points. Dotted line represents fold change equal to one. Two independent experiments pooled. Geometric mean with error bars representing SD. **g)** Luciferase activity measured from Jurkat-TonE reporter cells cultured for 24h with the indicated stimuli. **a-d, g)** One representative experiment out of two. **c-d)** Paired student t-test. **f)** Mann-Whitney test.

To assess the importance of TCR triggering on NFAT5 induction in the TME, we co-transferred P14- and OT-I-NFAT5_mCherry_ CD8 T cells into mice bearing two B16 tumors expressing the respective epitopes recognized by these TCRs; gp33 or OVA (Fig. 6b). We observed a TCR/peptide-MHC dependence of mCherry expression for P14 and OT-I CD8 TILs in both TME (Fig. 6c-d). Interestingly, OT-I CD8 T cells, which have a higher affinity to their cognate peptide than P14 CD8 T cells (K_d_ OT-I/SIINFEKL = 5.9 µM^37^ versus P14/gp33 = 3.5 µM^38^), showed an even stronger induction of mCherry in both the T-DLN and the tumor site of B16-OVA (Fig. 6c, d, Extended Data Fig. 4). Therefore, TCR stimulation in the TME is necessary to induce NFAT5 expression. Since TCR stimulation leads to Ca^2+^/calcineurin-induced NFAT1 or NFAT2 activation^13^, we wondered to which extent NFAT5 expression is dependent on this TCR-Ca^2+^/calcineurin-NFAT axis. The calcineurin inhibitor FK506 is widely used to block calcineurin targets such as NFAT1 and NFAT2. As in previous experiments, we transferred pre-activated P14-NFAT5_mCherry_ into B16-gp33-bearing mice. Mice received FK506 either the first three days (first phase) or the fourth to sixth day (second phase) post T cell transfer (Fig. 6e). The inhibition of calcineurin targets by FK506 partially decreased mCherry expression when FK506 was administrated during the first phase, while it had no effect during the second phase on mCherry expression (Fig. 6f). This effect was not observed in the T-DLN and N-DLN, showing a TME-specific effect of calcineurin inhibition on NFAT5 expression in the first phase (Extended Data Fig. 4). Strikingly, PD-1 and TOX levels showed similar trends, with a drastic decrease when FK506 was given during the first phase, but a mild decrease in the second phase (Fig. 6f), suggesting NFAT-independent regulation of PD-1 and TOX at later stages.

It was previously established that the concentration of K^+^, but not Na^+^ is increased in the TME of melanoma compared to the serum or healthy tissue from both mouse and human^39^. We questioned whether the ionic imbalance induced by K^+^ in the TME could participate in the transcriptional regulation of NFAT5 as we observed *in vitro*. We overexpressed KCNA3, a potassium channel enabling T cells to equalize intracellular potassium concentration in its wild type form (KCNA3), or in a non-conducting form (KCNA3 mutant) in activated P14-NFAT5_mCherry_ CD8 T cells and transferred them into B16-gp33 tumor-bearing mice. After 17 days, KCNA3-overexpressing P14 CD8 TILs showed a trend for decreased mCherry expression compared to the KCNA3 mutant control (Extended Data Fig. 4).

Beside the level of expression, the capacity of NFAT5 to act as a transcription factor is subject to further regulation^16, 40^. To test how different stimuli can impact the DNA binding capacity of NFAT5, we cloned a NFAT5 binding motif, the tonicity-responsive enhancer (TonE), into a luciferase-expressing lentiviral vector and transduced Jurkat cells, which express endogenous NFAT5 (Extended Data Fig. 4a). Culture of Jurkat cells (stably expressing the NFAT5 reporter construct) in the presence of NaCl or KCl induced luciferase activity in a dose-dependent manner. In contrast, TCR triggering, hypoxia, ROS inducers (Butyl-Hydroperoxid) or cytokine stimulation (TGF-β) did not upregulate NFAT5 activity, suggesting a dominant role for hypertonic stress to induce NFAT5 activity (Fig. 6g).

Altogether, TCR stimulation and hypertonicity regulated the NFAT5 transcriptional level, while only the osmolar changes increased NFAT5 activity *in vitro. In vivo*, TCR triggering is the main driver of NFAT5 transcription.

## Discussion

We showed that NFAT5, an unconventional member of the NFAT family, plays a crucial role in the regulation of tumor-induced T cell exhaustion. NFAT5 was expressed in CD8 TILs in various cancers (melanoma, adenocarcinoma, breast cancer) and in different species (mouse and human). In our study, overexpression of NFAT5 in tumor-specific CD8 T cells limited their anti-tumor response, while its inactivation strongly increased tumor control. NFAT5 deletion had a different effect depending on the subtype of CD8 TILs. NFAT5 KO Tpex showed increased production of cytokines and decreased levels of PD-1 and TOX, while their absolute number remained constant. In contrast, we found higher frequencies and absolute numbers of Tex, but the effect of NFAT5 deletion on the function of these cells was limited. These effects correlated with the NFAT5 expression levels in Tpex and Tex CD8 TILs, with significantly higher NFAT5 expression in Tpex compared to Tex. RNA-seq analysis confirmed that NFAT5 KO CD8 TILs expressed genes associated with early activation, with a signature that correlated with the one measured in TOX-KO CD8 TILs. We observed that the overexpression of NFAT5, or of a mutated form of NFAT1 unable to cooperate with AP-1 and involved in the regulation of exhaustion, has similar effects on the behavior and transcriptional program of tumor-specific T cells^13^. Since TOX is a direct target of NFAT1 and 2^9, 34^, our data suggest that a part of the effect of NFAT5 inactivation is mediated via the reduced level of TOX in Tpex. The TME inhibits NFAT activation through glucose deprivation or accumulation of lactic acid^41, 42^. This goes in parallel with an increased osmotic stress related to dead cell accumulation^39, 43^. We hypothesize that, at this stage, NFAT5 takes over classical NFAT to enforce an NFAT-induced transcriptional program and thus stabilizes the expression of exhaustion-associated genes.

Interestingly, the inactivation of NFAT5 in CD8 T cells during chronic infection with LCMV clone 13 did not improve viral control nor restore CD8 T cell functions. This argues in favor of a tumor-specific role of NFAT5, explained by a higher expression level of NFAT5 in CD8 TILs compared to CD8 T cells from chronic LCMV infection. Furthermore, our comparison was done at day 7 post-transfer for CD8 TILs, which does not correspond to the highest NFAT5 levels in Tpex CD8 TILs. NFAT5 is primarily described to regulate osmolarity-regulated genes, which were not differentially expressed in our experiments. However, NFAT5 activity is not limited to the regulation of this panel of genes. In macrophages, expression of NFAT5 drives a pro-inflammatory phenotype, which further supports T cell-mediated tumor control^19^. Interestingly, while NFAT5 drives the expression of tonicity-responsive genes in macrophages cultured in a hypertonic environment, it drives IL-6 production when stimulated with LPS^44^. Therefore, NFAT5 regulates gene expression in a context- and cell type-dependent manner. Similarly, while NFAT5 drives the expression of tonicity-responsive genes in T cells cultured in a hypertonic environment^45^, it was shown to negatively regulate IFN-γ production in CD4 T cells *in vitro*^20^.

We demonstrated that the main inducer of NFAT5 in T cells *in vivo* is TCR stimulation, at least in part through calcineurin activation, suggesting a regulation by classical members of the NFAT family. This regulation was more important during the first phase post T cell transfer (until day 3), while in a later phase (until day 6), calcineurin inhibition had a reduced effect on NFAT5 expression, but also on PD-1 and TOX levels. In addition to TCR stimulation, hypertonicity is a major inducer of NFAT5 *in vitro*, as well as an inducer of NFAT5 activity^16^. The massive death of tumor cells results in hypertonicity in the TME^39^. We showed that this factor slightly affected the regulation of NFAT5 expression within the tumor, but its main effect could be related to the activity of NFAT5 rather than its transcriptional regulation. Our study is in line with previous observations that increased osmolarity dampens T cell effector functions, unraveling another mechanism in place within the TME^39, 46^.

Altogether, we uncovered a new central player in the regulation of T cell exhaustion, acting only within tumors, but not during chronic infection. This discovery is particularly important in the frame of adoptive cell therapy (ACT) where a patients’ T cells are expanded before being transferred back into the patient. Acting on NFAT5, either genetically or using specific inhibitors, would favor a stronger T cell response against cancer, without decreasing the stemness capacity of transferred T cells.

## Figure legends

**Extended data Fig. 1:**
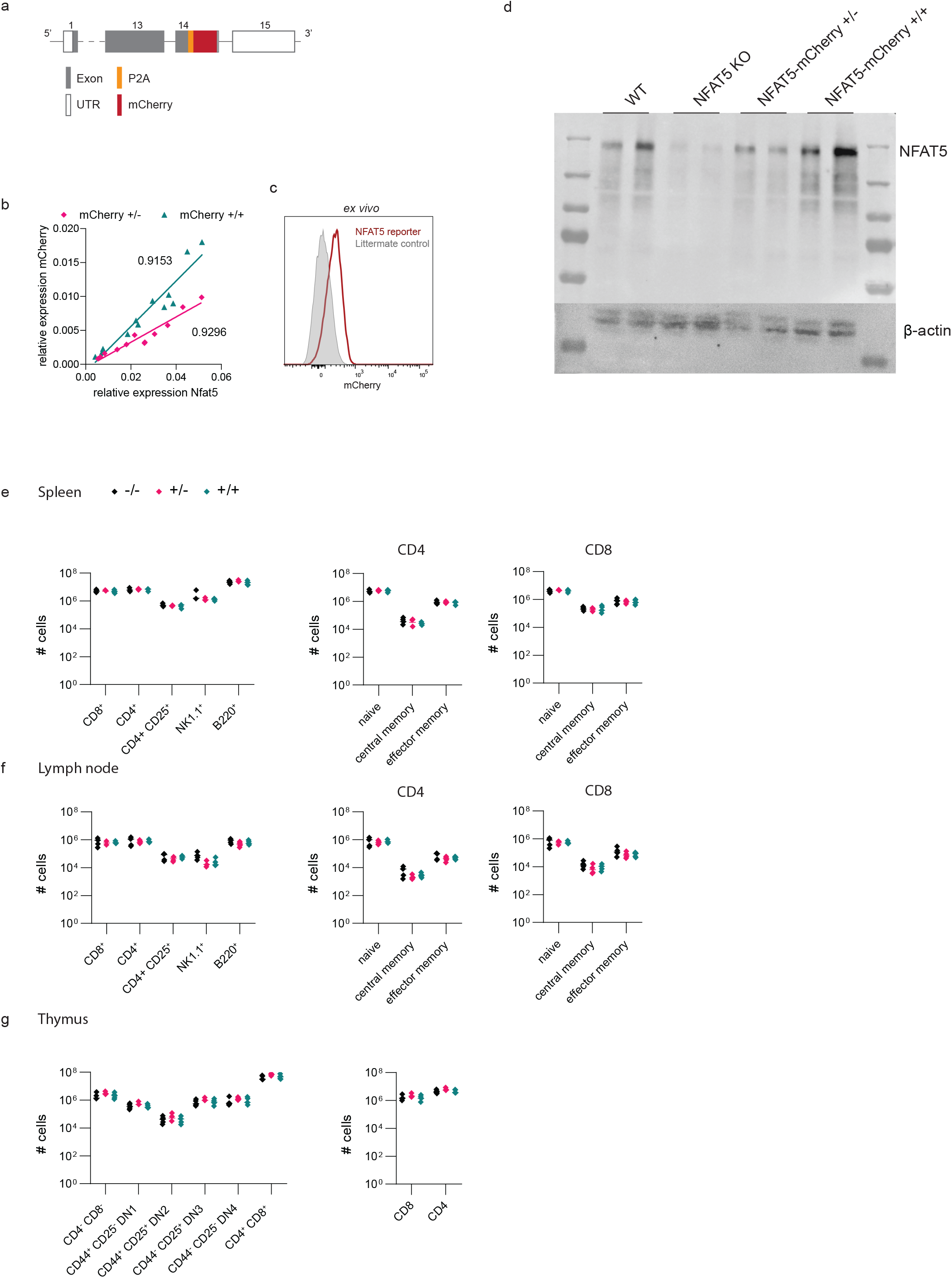
**a)** NFAT5 reporter mouse strain: The TAG stop codon in exon 14 of the mouse *Nfat5* gene was replaced by CRISPR/Cas-mediated genome engineering with the P2A-mCherry cassette to create a knock-in *NFAT5-P2A-mCherry* reporter model in C57BL/6 mice. **b)** Relative expression (2^-deltaCq^) of *mCherry* and *Nfat5* assessed by RT-PCR and normalized to *β-2-microglobulin* of NFAT5_mCherry+/-_ or NFAT5_mCherry+/1_ CD8 T cells cultured for 72h under different hyperosmolar conditions ranging from 300mOsm/kg to 500mOsm/kg. **c)** Histogram showing mCherry expression of NFAT5_mCherry+/+_ CD8 T cells from the lymph node in comparison to a littermate control mouse (mCherry-/-). **d)** Western blot showing the protein level of NFAT5^flox/flox^ CD4-Cre-/- (WT), CD4-Cre+/- (KO) NFAT5^flox/flox^, NFAT5_mCherry +/-_ and NFAT5_mCherry +/+_ P14 CD8 T cells cultured in complete RPMI with 1uM gp33 peptide and 20U/ml rhIL-2 for 72 hours. β-actin serves as a housekeeping gene. **e-g)** Spleen (**e**), lymph node (**f**) and thymus (**g**) from NFAT5_mCherry-/-,_ NFAT5_mCherry+/-_ and NFAT5_mCherry+/+_ mice were collected and analyzed for their immune compartment and thymic development **(g)** by flow cytometry.

**Extended data Fig. 2:**
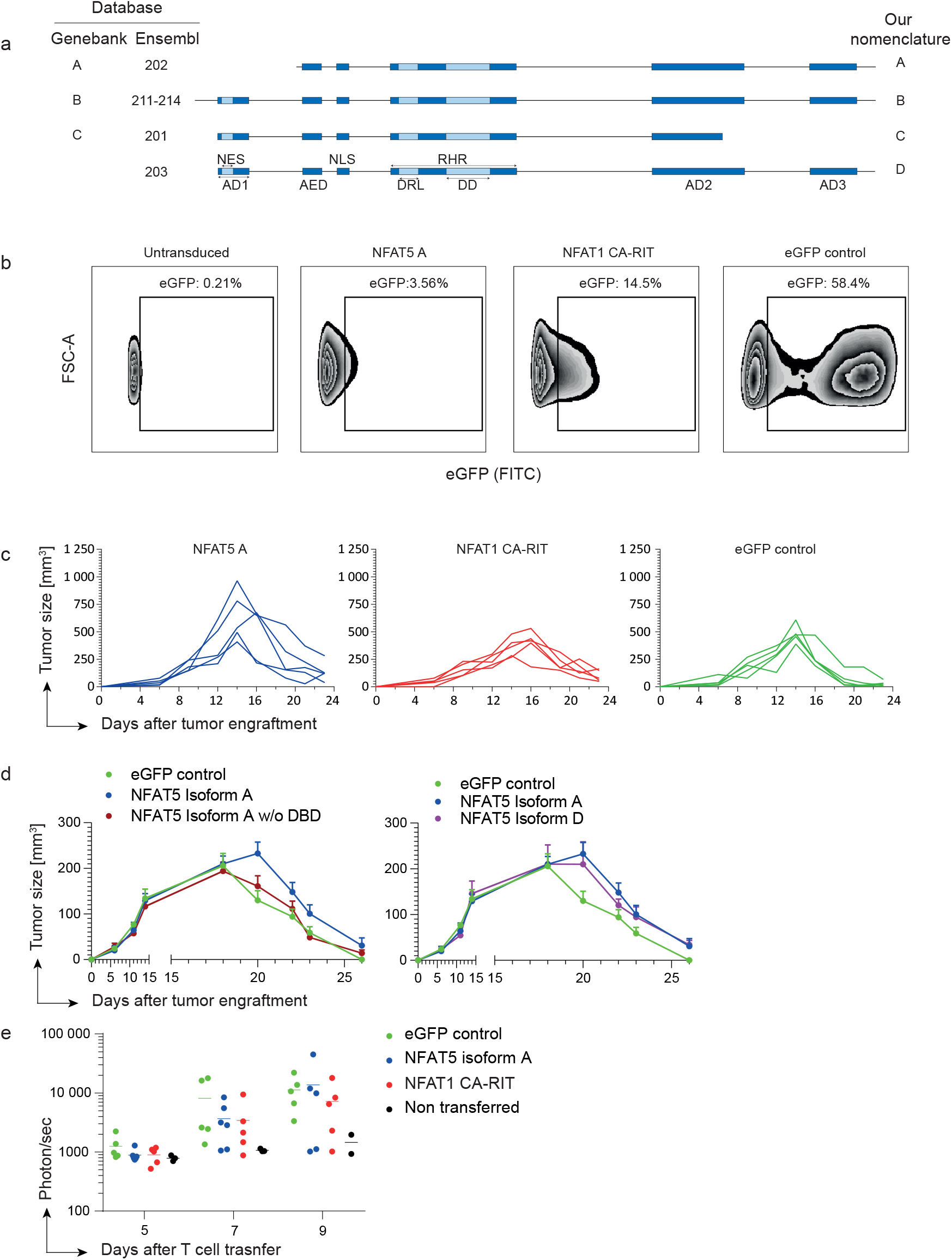
**a)** The four murine isoforms of NFAT5 were aligned on the software Geneious using the entries from genebank and Ensmbl. Domains described by Cheung *et al*. were aligned against mouse isoforms 203, whose length corresponds to human isoform C. NES: nuclear export signal, AD1: activation domain 1, AES: auxiliary export signal, DRL: DNA recognition loop, DD: dimerization domain, RHD: rel homology region, AD2: activation domain 2, AD3: activation domain 3. **b)** Gating for the sorting of NFAT5 A, NFAT1 CA-RIT or control eGFP transduced CD8 T cells. **c)** Individual tumor growth per mouse per group as described in Figure 2. **d)** Tumor growth comparing NFAT5 isoform A lacking the DNA binding domain (DBD) (left panel) and NFAT5 isoform D (right panel) to the one of NFAT5 isoform A. **e)** Bioluminescence of the luciferase expressing TCRP1A CD8 T cells after injection of the mice with luciferin.

**Extended data Fig. 3:**
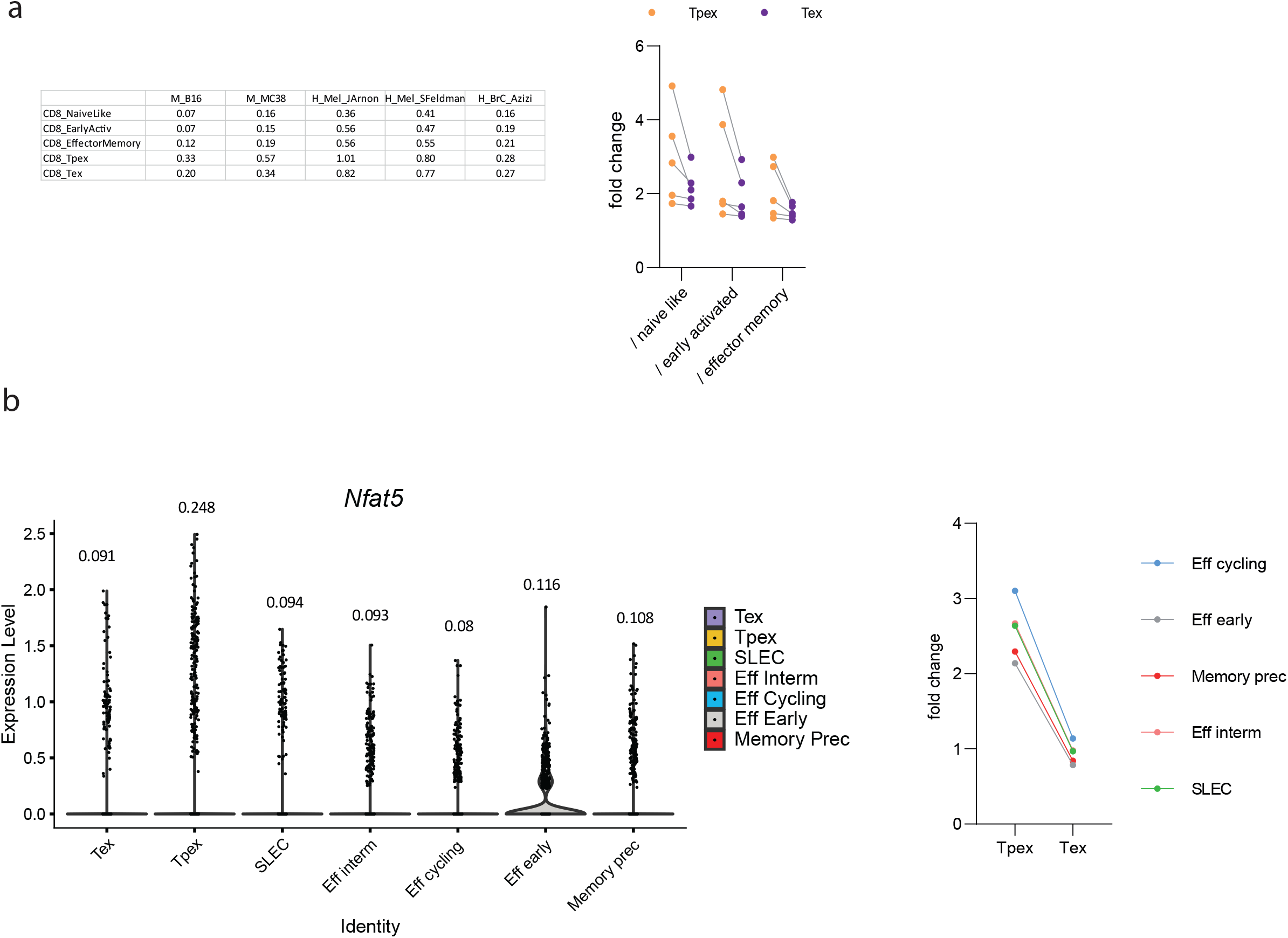
**a)** Mean *Nfat5* expression level of various CD8 T cell populations in indicated studies assessed by TILatlas (left) and the fold change of *Nfat5* expression level of Tpex or Tex over indicated populations (right). **b)** Violin plot showing the *Nfat5* expression level (number above) in LCMV infection (left) and fold change of *Nfat5* expression level of Tpex or Tex over indicated populations (right).

**Extended data Fig. 4:**
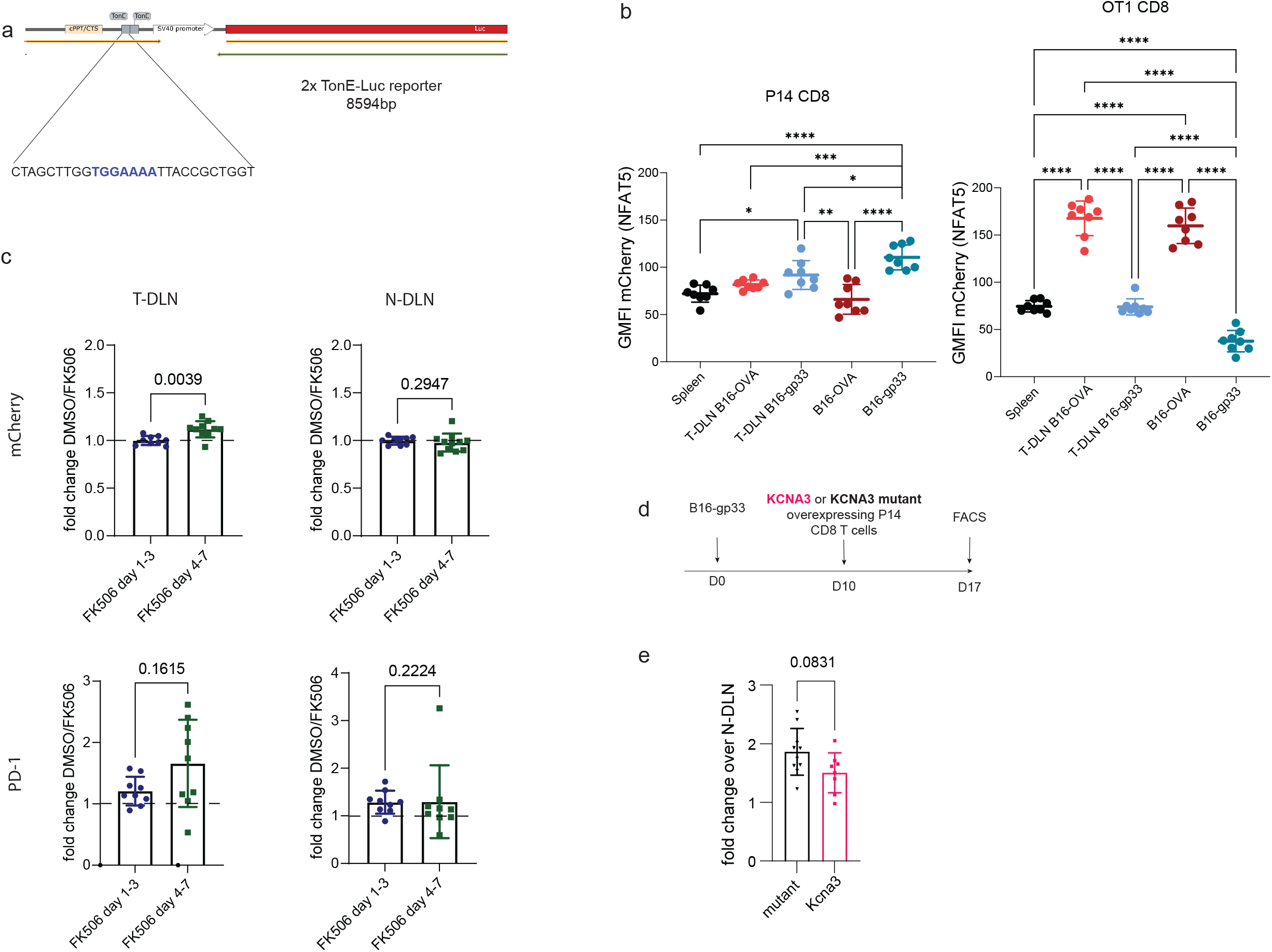
**a)** Schematic representation of the luciferase NFAT5 activity reporter plasmid. Blue sequence represents the NFAT5 (TonEBP) binding site (TonE). **b)** mCherry expression level in the spleen, T-DLN, and tumor of B16-gp33 / B16-OVA-bearing mice transferred with P14-(left) and OT-I-(right) NFAT5_mCherry+/+_ CD8 T cells. Two ways ANOVA. **c)** Fold change of mCherry (upper panel) or PD-1 (lower panel) calculated by DMSO receivers over FK506 receivers during indicated time periods in the T-DLN and N-DLN. **d)** Timeline of the experiment. B16-gp33 bearing mice were transferred with either KCNA3 or KCNA3 mutant overexpressing P14-NFAT5_mCherry+/+_ CD8 T cells ten days after tumor injection. **e)** mCherry expression on day 17 plotted as fold change of P14 TILs over P14 CD8 T cells from N-DLN.

## Materials and methods

### Patient material

To assess NFAT5 expression in human TILs by qPCR, we used amplified cDNA from Melan-A-specific CD8 TILs, EBV-specific and naïve (CD8^+^CD45^+^CCR7^+^CD27^+^CD28^+^) CD8 T cells isolated from healthy donor PBMCs, patient PBMCs and metastatic lymph nodes from stage III/IV metastatic melanoma patients (clinical study NCT00112229)^14^.

### Animals

CD45.1, CD45.1.2 and Rag1^-/-^B10D2 TCRP1A mice were bred in house. CD45.2 CD4-Cre NFAT5^flox/flox^ mice were kindly provided by Prof. Cristina López-Rodríguez^45^. NFAT5-mCherry reporter mice were generated by Cyagen. The TAG stop codon in exon 14 of the mouse *Nfat5* gene was replaced by CRISPR/Cas-mediated genome engineering with the P2A-mCherry cassette on a C57BL/6 background. CD45.2 CD4-Cre NFAT5^flox/flox^ mice and NFAT5-mCherry reporter mice were crossed with P14 or OT-I TCR transgenic mice and kept on a C57BL/6 background. Mice were kept in an SPF animal facility. Experiments were approved by the veterinarian authorities and performed in compliance with the University of Lausanne internal regulations (authorization VD2943, VD359). Tumor volume was calculated with the following formula: volume [mm^3^] = length [mm] x width [mm] x height [mm].

### Cell lines

Complete medium was composed of 10% heat-inactivated FBS (Gibco), penicillin/streptomycin 100 U/ml (Gibco), Hepes 10mM (Gibco), 1mM sodium pyruvate (Gibco) and 50μM 2-mercaptoethanol (Gibco). B16-gp33^27^ and B16-OVA were cultured in complete DMEM GlutaMAX™-I with 100μg/ml G418 (Calbiochem). P511^29^, Jurkat and primary T cells were cultured in complete RPMI 1640 GlutaMAX™-I. PlatinumE (PlatE) cells were cultured in complete DMEM with 10ug/ml Blasticidin (Invivogen) and 1ug/ml Puromycin (Invivogen).

### Flow cytometry

The following protocol was used for all tumor experiments: after the Fc receptor of the cells was blocked with anti-mouse CD16/32 (Biolegend, 101320), extracellular staining was performed in FACS buffer for 30 minutes. Dead cells were stained with Aqua Vivid (Invitrogen, L34966) in PBS for 15-20 minutes or by adding DAPI (Thermo Fischer Scientific, D3571) directly before flow cytometry analysis. After 20 minutes of fixation, intracellular staining was performed for 30 minutes. The Biolegend intracellular staining kit (421002) was used for cytokines and the FoxP3 staining kit (00-5523-00) was used for transcription factor staining. When cytokine levels were assessed, the cells were stimulated with their cognate peptide gp33 (10^−6^ M) and Golgistop (BD, 554724) for 5 hours. Antibodies are listed in Extended Data Table 2.

### RNA extraction for sequencing / RT-qPCR

Cells were centrifuged for 5 minutes at maximum speed and RNA was extracted with the RNeasy Plus Micro Kit (Qiagen) following the manufacturer’s recommendations. For RNA sequencing, RNA quality was measured with a fragment analyzer. Reverse transcription was achieved using the High-capacity cDNA Reverse Transcription kit (Applied biosystems). For qPCR, KAPA SYBR® Fast qPCR Master Mix (2x) Kit (Sigma), was used. PCR amplification was performed in a 48 well plate (Illumina) on an Eco machine (Illumina). Primer pairs: β2M-F: AGACTGATACATACGCCTGCAG, β2M-R GCAGGTTCAAATGAATCTTCAG, Murine NFAT5-F: GGTACAGCCTGAAACCCAAC, Murine NFAT5-R TGCAACACCACTGGTTCATT, Human NFAT5-F: ATT GCA AAA CCA AGG GAA CA, Human NFAT5-R: TTG GAA TCA GGA TTT TCT TCG, mCherry-F: CCC ACA ACG AGG ACT ACA CC, mCherry-R: TTG TAC AGC TCG TCC ATG CC.

### Vectors

NFAT1 CA-RIT (IRES-GFP) retroviral vector was obtained from Addgene (plasmid # 85181). MSCV-Kcna3-Thy1.1 (pMSCV-Thy1.1:F2A:mKcna3[NM_008418.2]) and non-conducting ‘pore dead’ construct MSGV-Kcna3-Thy1.1 (pMSCV-Thy1.1:F2A:mKcna3 W389F) (referred to as Kcna3 mutant) were generated by Vector builder.

Cloning of the four isoforms of NFAT5: The NFAT5 coding sequences were added after an enhanced GFP (eGFP) separated by the self-cleaving peptide P2A. The stop codon was removed and a FlagTag sequence was added at the end of the NFAT5 sequence. The four isoforms differ in the first and last exons. We first cloned the N- and C-termini of NFAT5 isoform A synthetized by Addgene into a pMSGV retroviral vector. We then inserted the core murine cDNA of NFAT5 obtained on the transOMIC platform to create the isoform A. We modified the first and last exon to achieve the three other complete isoforms by replacing the N- and C-termini sequences by newly synthetized sequences generated on GeneArt.

### Overexpression experiments

TCRP1A CD8 T cells or P14-NFAT5_mCherry_ cells isolated from the lymph nodes. Lymphocytes were activated with 1ug/ml of their respective peptides (P1A (LPYLGWLVF) or gp33 (KAVYNFATC)) and 20U/ml rhIL-2 (Proleukin Aldesleukin) one day before transduction. Viruses were produced in PlatE cells as previously described ^47^. Briefly, transfection of the respective plasmids was done with lipofectamine 2000 (Life technologies) in DMEM. Viral supernatants were collected and filtered with 0.45μm filters (Sarstedt Ag & Co) and either used as crude supernatant or concentrated. Transduced CD8 T cells were injected i.v. (5×10^6^) into tumor-bearing hosts one day post transduction. For NFAT5 overexpression, eGFP positive sorted cells were collected and injected i.v. into tumor-bearing mice (minimum 1×10^4^ transduced cells per mouse). To follow CD8 T cell infiltration, mice were injected intraperitoneal with 3mg of luciferin (Biosynth, L-8220), anesthetized with isoflurane (about 4% in air) and bioluminescence was captured with an IVIS LUMINA II machine. P511 tumors were cut into pieces and digested with 1mg/ml collagenase I and 100μg/ml DNAse (Sigma) for 30 minutes at 37°C. Tumors were passed through a 70μm cell strainer (Falcon). T cells from P511 were isolated with a Ficoll gradient (Fresenius Kabi Norge AS).

### T cell transfer experiments in tumor-bearing mice (B16-gp33 and B16-OVA)

LN cell suspensions were cultured in complete RPMI + 1µg/ml gp33 peptide (KAVYNFATC) or 1µg/ml Ovalbumin peptide (SIINFEKL) + 20U/ml rhIL-2 (Proleukin Aldesleukin) for two days. CD8 T cells were counted and resuspended into PBS before i.v. injection (5×10^6^ cells/mouse). On the day of the analysis, tumors were collected in complete RPMI and passed through 70μm cell strainers (Falcon). T cells from tumors were isolated with a 40/70 percoll gradient (VWR 17-0891). When sorted for RNA sequencing, the cells were collected in RNA later (Invitrogen).

### T cell transfer in LCMV-infected mice

Vβ5 mice were infected with 2×10^6^ PFU one day after transfer of 1×10^3^ P14 WT or NFAT5 KO P14 CD8 T cells. To stimulate cells for cytokine staining, cells were incubated 30 minutes with 1μM gp33 before addition of 5μg/ml of brefeldin A (Biolegend). The cells were then incubated 4 hours 30 minutes before staining.

### Jurkat TonE-NFAT5 reporter

Jurkat-TonE was generated by lentiviral transduction of Jurkat cells with a plasmid encoding for luciferase under the control of 2x TonE promoter. 5×10^5^ reporter-expressing Jurkat cells were plated in a 96-well plate and cultured for 24 hours with complete RPMI supplemented with the indicated (10ng PMA (Sigma-Aldrich) / 200ng ionomycin (Thermo Fisher), for mouse 10ug/mL CD3 (OKT3 - eBioscience)/CD28 (CD28.2 – eBioscience), ROS: 10uM Butyl-Hydroperoxid, 10ng/mL TGF-β (Roche), 0,5% O_2_ or NaCl / KCl as indicated)..

### Statistical and bioinformatic analyses

Statistical analyses were performed with Graphpad Prism version 9. Statistical tests are indicated in the legends. Comparisons of more than two groups and subsequent p-values were calculated by ANOVAs with corrections as needed and specified below the figures. A p-value of <0.05 was used as the threshold to define statistical significance.

### RNA-seq-Transcript quantification

Transcript abundance quantification was performed with Salmon 0.14.1 ^48^ in quasi-mapping-based mode using the mouse reference transcriptome (assembly GRCm38.p2) obtained from ENSEMBLE ^49^. Default parameters were used plus the --seqBias, --gcBias, --validateMappings, --fldMean 200 and – fldSD 30 parameters.

### Differential expression analyses

Differential gene expression analyses were performed using DESeq2 1.24.0 ^50^. Transcript-level abundances were summarized at the gene level using tximport 1.12.3 ^51^. Genes with low read counts were filtered out (requiring genes to have a count of at least 10 in at least a number of samples equal to the smallest group size). Overall similarity between samples was assessed by first applying a regularized stabilizing transformation (rlog) to the gene-level count matrices using the *rlog* function, and then performing a principal components analysis (PCA) on the regularized matrix using the *plotPCA* function. Significant genes were identified using a false discovery rate (FDR) threshold of 5% and an absolute log2 fold change threshold of 1.5. The GO term analysis were performed with the EnrichR platform ^52, 53^ using the “GO Biological Process 2018” algorithm. Enrichment of the NFAT5 KO differentially expressed gene set in TOX up or downregulated genes was evaluated using the TOX expression data published in ^34^, available from the Gene Expression Omnibus (GEO) database under accession number GSE126973. NFAT5 KO differentially expressed genes were ranked based on the log2 fold change (ranked list), and TOX signatures of up or down gene regulation were created based on TOX knockout differentially expressed genes with positive or negative log2 fold change. Enrichment scores and adjusted p-values were computed using the *fgsea* function and the ranked list and regulation signatures mentioned above, with the number of permutations set to 1,000. Enrichment plots were obtained using the *plotEnrichment* function.

### TIL atlas

The TIL atlas dataset used in Fig.1a, b and Extended data Fig.3, which includes 16,803 high-quality single-cell transcriptomes from 25 samples (B16 melanoma and MC38 colon adenocarcinoma tumors) from six different studies, has been collected, thoroughly analyzed and annotated by Andreatta and co-workers^26^, and it is publicly available (https://doi.org/10.6084/m9.figshare.12478571).

### Regulon analysis of tumor-infiltrating T lymphocyte

Regulons (gene sets regulated by the same transcription factor) and their activity were inferred and evaluated using the SCENIC pipeline (https://scenic.aertslab.org)^54^, which can be described in three steps. Step 1) Infer gene regulatory network (GRN) using grnBoost2, which is a faster implementation of the original algorithm Genie3^55^; scRNA-seq transcriptomics data is used as the input to infer causality from the expression levels of the transcription factors to the targets based on co-expression patterns. The importance of each transcription factor in the prediction of the target gene expression pattern is taken as an indication of a putative regulatory event. The aggregation of the top 50 targets per TF and the top 5 TF per target was used to define raw putative regulons. Step 2) Co-expression modules (raw putative regulons, i.e. sets of genes regulated by the same transcription factor) derived from the GRN generated in Step 1 are refined by pruning indirect targets by motif discovery analysis using cisTarget algorithm and a cis-regulatory motif database^56, 57^. We used mm9-500bp-upstream-7species.mc9nr.feather and mm9-tss-centered-10kb-7species.mc9nr.feather databases. The motif database includes a score for each pair motif-gene, so that a motif-gene ranking can be derived. A motif enrichment score is then calculated for the list of transcription factor selected targets by calculating the Area Under the recovery Curve (AUC) on the motif-gene ranking 1 using the RcisTarget R package (https://github.com/aertslab/RcisTarget). If a motif is enriched among the list of transcription factor targets, a regulon is derived including the target genes with a high motif-gene score. Step 3) evaluation of regulon activity of each individual cell using AUCell (https://github.com/aertslab/AUCell), which provides an AUC score for each regulon; we discarded regulons with less than 5 constituent elements, as the estimation of the activity of small regulons is less reliable. In order to compare the regulon activity profile of CD8 exhausted T cells and CD8 naïve cells from the TILs dataset, we used the regulon activity (AUC score) matrix and performed, for each regulon, a Wilcoxon Rank Sum test implemented within the FindMarkers function of the SEURAT R package (version 4.0.3). Only regulons with an adjusted p-value for this test of 0.05 or less were considered as differentially active (Bonferroni correction).

### Evaluation of NFAT5 KO signature in tumor-infiltrating T lymphocytes

Differentially expressed genes up (n=458) and down (n=832) regulated from the comparison NFAT5.KO Vs. WT were considered separately as two different signatures to be analyzed in the TILs dataset (NFAT5.KO.DEG.UP and NFAT5.KO.DEG.DOWN, respectively). The AUCell R package^54^ was used to evaluate these signatures across tumor-infiltrating T lymphocyte subpopulations using the normalized gene expression matrix of this dataset.

### Western Blot

NFAT5^flox/flox^ CD4-Cre-/- (WT), CD4-Cre+/- (KO) NFAT5^flox/flox^, NFAT5_mCherry +/-_ and NFAT5_mCherry +/+_ P14 CD8 T cells were cultured in complete RPMI with 1uM gp33 peptide and 20U/ml rhIL-2 (Proleukin Aldesleukin) for 72 hours. Whole cell extracts were isolated using RIPA lysis buffer (25mM Tris HCl pH 7.6, 150mM NaCl, 1% NP 40, 1% sodium deoxychlorate, 0,1% SDS; Cell Signaling Technology) supplemented with DNase, 1x phosphatase inhibitor (PhosSTOP Roche) and 1x protease inhibitor (complete protease inhibitor cocktail Roche). Protein concentration was assessed by the Bradford Assay (Biorad Protein Assay Kit II) and 25ug of cell lysates from each condition were resolved by 8% SDS-PAGE with NuPAGE electrophoresis system (Invitrogen/Thermo Fisher Scientific). Proteins were then transferred to PVDF membrane (Invitrogen/Thermo Fisher Scientific) by Trans-Blot SD semidry transfer cell (Bio-Rad) at 18 V for 30 minutes. Membranes were blocked with 5% w/v nonfat dry milk (Sigma-Aldrich) and then incubated with the indicated primary antibodies at 4 °C overnight. The bands were visualized using Thermo Scientific™ Pierce™ ECL Western Blotting Substrate after incubation with horseradish peroxidase (HRP)-conjugated antibodies for 1 hour at room temperature. Primary antibodies: anti-NFAT5 antibody (Santa Cruz) and anti-beta-actin antibody (Invitrogen).

## Acknowledgement

We thank Hana Zdimerova for editing the manuscript, Dr Ping-Chi Ho and Dr Thierry Walzer for suggestions about the project and Patrick Reichenbach for technical counseling. This work was funded by the University of Lausanne, and grants from the Max Cloëtta Foundation (GV), the Swiss National Science Foundation (GV: 310030_182680, WH: 310030B_179570), ISREC institute (DC), the Swiss cancer league (GV).

## Notes

### Competing Interest Statement

The authors have declared no competing interest.

