## Supplemental Table 1 for "NFAT5 induction by the tumor microenvironment enforces CD8 T cell exhaustion": extended data Table 1.pdf

| | baseMean | log2FoldCha | lfcSE | stat | pvalue | padj |
|---|---|---|---|---|---|---|
| Hc (ENSMUSG00000026874) | 22,76 | 7,49 | 1,89 | 3,97 | 0,00 | 0,00 |
| Gm43024 (ENSMUSG00000105472) | 16,01 | 7,49 | 1,71 | 4,38 | 0,00 | 0,00 |
| Hrh4 (ENSMUSG00000037346) | 12,17 | 7,10 | 2,10 | 3,38 | 0,00 | 0,01 |
| F3 (ENSMUSG00000028128) | 12,09 | 7,09 | 1,80 | 3,94 | 0,00 | 0,00 |
| Nhsl2 (ENSMUSG00000079481) | 22,51 | 6,76 | 1,97 | 3,43 | 0,00 | 0,01 |
| Gm42515 (ENSMUSG00000105378) | 9,01 | 6,66 | 1,79 | 3,72 | 0,00 | 0,00 |
| D830024N08Rik (ENSMUSG00000110580) | 9,00 | 6,66 | 2,18 | 3,05 | 0,00 | 0,02 |
| Fscn1 (ENSMUSG00000029581) | 8,79 | 6,62 | 2,39 | 2,77 | 0,01 | 0,04 |
| Sap25 (ENSMUSG00000079165) | 8,64 | 6,60 | 1,67 | 3,95 | 0,00 | 0,00 |
| Gm44901 (ENSMUSG00000108954) | 8,39 | 6,56 | 2,21 | 2,97 | 0,00 | 0,03 |
| Zbed6 (ENSMUSG00000102049) | 8,12 | 6,51 | 2,27 | 2,86 | 0,00 | 0,04 |
| Mrvi1 (ENSMUSG00000005611) | 11,40 | 6,43 | 1,16 | 5,52 | 0,00 | 0,00 |
| 4931415C17Rik (ENSMUSG00000054910) | 7,35 | 6,37 | 2,23 | 2,86 | 0,00 | 0,04 |
| Capn11 (ENSMUSG00000058626) | 7,12 | 6,32 | 1,77 | 3,58 | 0,00 | 0,01 |
| Cbs (ENSMUSG00000024039) | 13,92 | 6,26 | 1,88 | 3,33 | 0,00 | 0,01 |
| Ripor3 (ENSMUSG00000074577) | 10,15 | 6,24 | 1,72 | 3,62 | 0,00 | 0,00 |
| Yes1 (ENSMUSG00000014932) | 6,19 | 6,12 | 1,76 | 3,47 | 0,00 | 0,01 |
| Olfr1382 (ENSMUSG00000063827) | 6,06 | 6,09 | 1,91 | 3,19 | 0,00 | 0,02 |
| Trim17 (ENSMUSG00000036964) | 7,76 | 5,85 | 1,55 | 3,77 | 0,00 | 0,00 |
| Gm26890 (ENSMUSG00000097246) | 11,27 | 5,80 | 1,94 | 3,00 | 0,00 | 0,03 |
| Gm13369 (ENSMUSG00000083796) | 6,24 | 5,53 | 2,01 | 2,75 | 0,01 | 0,05 |
| Dusp8 (ENSMUSG00000037887) | 16,78 | 5,09 | 1,18 | 4,30 | 0,00 | 0,00 |
| Gm43753 (ENSMUSG00000105707) | 21,00 | 5,02 | 1,24 | 4,04 | 0,00 | 0,00 |
| Gm47404 (ENSMUSG00000113261) | 5,72 | 4,81 | 1,63 | 2,95 | 0,00 | 0,03 |
| Gm49883 (ENSMUSG00000117113) | 5,28 | 4,69 | 1,62 | 2,89 | 0,00 | 0,03 |
| Gm37423 (ENSMUSG00000104436) | 6,44 | 4,48 | 1,50 | 2,99 | 0,00 | 0,03 |
| Gm35572 (ENSMUSG00000110462) | 6,19 | 4,38 | 1,24 | 3,55 | 0,00 | 0,01 |
| Gm47963 (ENSMUSG00000110993) | 10,95 | 4,21 | 0,98 | 4,29 | 0,00 | 0,00 |
| C230038L03Rik (ENSMUSG00000085560) | 6,46 | 4,02 | 1,27 | 3,16 | 0,00 | 0,02 |
| Ppp2r3d (ENSMUSG00000099077) | 5,89 | 3,84 | 1,36 | 2,81 | 0,00 | 0,04 |
| Gm2449 (ENSMUSG00000097223) | 12,15 | 3,65 | 1,05 | 3,47 | 0,00 | 0,01 |
| Chil5 (ENSMUSG00000043873) | 62,53 | 3,52 | 0,48 | 7,42 | 0,00 | 0,00 |
| Gm4787 (ENSMUSG00000072974) | 10,62 | 3,38 | 1,22 | 2,76 | 0,01 | 0,04 |
| Plekha7 (ENSMUSG00000045659) | 106,38 | 3,24 | 0,95 | 3,41 | 0,00 | 0,01 |
| Gm36931 (ENSMUSG00000103779) | 7,62 | 3,17 | 1,12 | 2,83 | 0,00 | 0,04 |
| Adamtsl4 (ENSMUSG00000015850) | 31,29 | 3,14 | 1,04 | 3,03 | 0,00 | 0,02 |
| 9530027J09Rik (ENSMUSG00000097197) | 11,28 | 3,12 | 1,14 | 2,73 | 0,01 | 0,05 |
| Cpne5 (ENSMUSG00000024008) | 18,96 | 3,09 | 1,08 | 2,85 | 0,00 | 0,04 |
| Pmepa1 (ENSMUSG00000038400) | 126,41 | 3,06 | 0,72 | 4,25 | 0,00 | 0,00 |
| C230085N15Rik (ENSMUSG00000102212) | 305,65 | 2,90 | 0,82 | 3,55 | 0,00 | 0,01 |
| Gm11400 (ENSMUSG00000081208) | 28,19 | 2,76 | 0,74 | 3,74 | 0,00 | 0,00 |
| Fam84b (ENSMUSG00000072568) | 127,20 | 2,65 | 0,43 | 6,18 | 0,00 | 0,00 |
| Atp8a2 (ENSMUSG00000021983) | 32,06 | 2,52 | 0,78 | 3,24 | 0,00 | 0,01 |
| Phxr4 (ENSMUSG00000031802) | 29,02 | 2,52 | 0,88 | 2,86 | 0,00 | 0,04 |
| Mast1 (ENSMUSG00000053693) | 7,99 | 2,51 | 0,84 | 2,97 | 0,00 | 0,03 |
| Pisd-ps1 (ENSMUSG00000082286) | 32,53 | 2,44 | 0,71 | 3,44 | 0,00 | 0,01 |
| Gm44694 (ENSMUSG00000108633) | 25,77 | 2,40 | 0,69 | 3,47 | 0,00 | 0,01 |
| Gm49759 (ENSMUSG00000111394) | 48,24 | 2,28 | 0,81 | 2,81 | 0,00 | 0,04 |
| Cxadr (ENSMUSG00000022865) | 26,26 | 2,19 | 0,78 | 2,81 | 0,00 | 0,04 |
| Gm26816 (ENSMUSG00000097380) | 37,99 | 2,18 | 0,78 | 2,79 | 0,01 | 0,04 |
| Wfikkn2 (ENSMUSG00000044177) | 28,25 | 2,17 | 0,75 | 2,90 | 0,00 | 0,03 |

| | | | | | | |
|---|---|---|---|---|---|---|
| Gm26648 (ENSMUSG00000097275) | 38,39 | 2,14 | 0,69 | 3,08 | 0,00 | 0,02 |
| Prpf40b (ENSMUSG00000023007) | 74,71 | 2,09 | 0,57 | 3,65 | 0,00 | 0,00 |
| Gm4208 (ENSMUSG00000100426) | 97,91 | 2,08 | 0,47 | 4,38 | 0,00 | 0,00 |
| Inka2 (ENSMUSG00000048458) | 95,32 | 2,05 | 0,62 | 3,32 | 0,00 | 0,01 |
| E430014B02Rik (ENSMUSG00000102973) | 57,52 | 1,98 | 0,67 | 2,95 | 0,00 | 0,03 |
| Cd40lg (ENSMUSG00000031132) | 324,38 | 1,97 | 0,30 | 6,49 | 0,00 | 0,00 |
| Ramp3 (ENSMUSG00000041046) | 299,10 | 1,94 | 0,47 | 4,15 | 0,00 | 0,00 |
| Tle1 (ENSMUSG00000008305) | 191,46 | 1,93 | 0,35 | 5,55 | 0,00 | 0,00 |
| Esm1 (ENSMUSG00000042379) | 38,94 | 1,93 | 0,69 | 2,79 | 0,01 | 0,04 |
| AA386476 (ENSMUSG00000074357) | 16,07 | 1,87 | 0,62 | 3,01 | 0,00 | 0,02 |
| Vps13d (ENSMUSG00000020220) | 170,08 | 1,87 | 0,51 | 3,68 | 0,00 | 0,00 |
| Pde2a (ENSMUSG00000110195) | 1675,76 | 1,85 | 0,39 | 4,74 | 0,00 | 0,00 |
| Gm21897 (ENSMUSG00000094472) | 26,55 | 1,85 | 0,64 | 2,88 | 0,00 | 0,03 |
| Gm26885 (ENSMUSG00000097125) | 143,01 | 1,85 | 0,39 | 4,73 | 0,00 | 0,00 |
| Enc1 (ENSMUSG00000041773) | 72,41 | 1,85 | 0,58 | 3,19 | 0,00 | 0,02 |
| Gdap10 (ENSMUSG00000096954) | 156,72 | 1,81 | 0,45 | 4,00 | 0,00 | 0,00 |
| Tdrp (ENSMUSG00000050052) | 304,64 | 1,77 | 0,33 | 5,33 | 0,00 | 0,00 |
| BC043934 (ENSMUSG00000056418) | 141,49 | 1,76 | 0,60 | 2,91 | 0,00 | 0,03 |
| 5830444F18Rik (ENSMUSG00000102744) | 97,98 | 1,76 | 0,47 | 3,77 | 0,00 | 0,00 |
| Usp31 (ENSMUSG00000063317) | 648,68 | 1,75 | 0,47 | 3,75 | 0,00 | 0,00 |
| Smox (ENSMUSG00000027333) | 1137,14 | 1,73 | 0,36 | 4,86 | 0,00 | 0,00 |
| Wsb2-ps (ENSMUSG00000084020) | 63,31 | 1,71 | 0,38 | 4,45 | 0,00 | 0,00 |
| S1pr1 (ENSMUSG00000045092) | 1522,02 | 1,69 | 0,36 | 4,64 | 0,00 | 0,00 |
| Znrf3 (ENSMUSG00000041961) | 157,91 | 1,68 | 0,46 | 3,62 | 0,00 | 0,00 |
| Klf3 (ENSMUSG00000029178) | 969,64 | 1,64 | 0,31 | 5,24 | 0,00 | 0,00 |
| Kcnj8 (ENSMUSG00000030247) | 58,28 | 1,62 | 0,53 | 3,03 | 0,00 | 0,02 |
| Ttc39c (ENSMUSG00000024424) | 1058,72 | 1,62 | 0,40 | 4,04 | 0,00 | 0,00 |
| Gm20696 (ENSMUSG00000030068) | 278,91 | 1,61 | 0,48 | 3,38 | 0,00 | 0,01 |
| Ccr8 (ENSMUSG00000042262) | 656,90 | 1,60 | 0,34 | 4,69 | 0,00 | 0,00 |
| Lamc2 (ENSMUSG00000026479) | 99,56 | 1,59 | 0,58 | 2,75 | 0,01 | 0,05 |
| Ccdc184 (ENSMUSG00000029875) | 94,85 | 1,56 | 0,48 | 3,26 | 0,00 | 0,01 |
| Mir155hg (ENSMUSG00000097418) | 54,76 | 1,55 | 0,54 | 2,85 | 0,00 | 0,04 |
| Gzma (ENSMUSG00000023132) | 3174,89 | 1,52 | 0,48 | 3,17 | 0,00 | 0,02 |
| Hs3st3b1 (ENSMUSG00000070407) | 160,94 | 1,52 | 0,42 | 3,63 | 0,00 | 0,00 |
| Zbtb37 (ENSMUSG00000043467) | 87,51 | 1,50 | 0,52 | 2,85 | 0,00 | 0,04 |
| Nr1d2 (ENSMUSG00000021775) | 417,09 | 1,49 | 0,34 | 4,37 | 0,00 | 0,00 |
| Galnt10 (ENSMUSG00000020520) | 2218,68 | 1,48 | 0,24 | 6,06 | 0,00 | 0,00 |
| Rarg (ENSMUSG00000001288) | 72,72 | 1,48 | 0,49 | 3,00 | 0,00 | 0,03 |
| Rbpms2 (ENSMUSG00000032387) | 66,12 | 1,46 | 0,51 | 2,87 | 0,00 | 0,04 |
| Gm27194 (ENSMUSG00000098292) | 106,78 | 1,45 | 0,30 | 4,82 | 0,00 | 0,00 |
| Gm37758 (ENSMUSG00000102712) | 429,98 | 1,44 | 0,18 | 8,02 | 0,00 | 0,00 |
| Zbtb42 (ENSMUSG00000037638) | 199,88 | 1,43 | 0,41 | 3,54 | 0,00 | 0,01 |
| Camsap2 (ENSMUSG00000041570) | 268,12 | 1,43 | 0,37 | 3,87 | 0,00 | 0,00 |
| Pou6f1 (ENSMUSG00000009739) | 397,42 | 1,43 | 0,42 | 3,39 | 0,00 | 0,01 |
| Farp2 (ENSMUSG00000034066) | 63,19 | 1,42 | 0,51 | 2,80 | 0,01 | 0,04 |
| Hcn3 (ENSMUSG00000028051) | 36,52 | 1,41 | 0,51 | 2,76 | 0,01 | 0,05 |
| Cnnm3 (ENSMUSG00000001138) | 292,52 | 1,40 | 0,45 | 3,15 | 0,00 | 0,02 |
| Fcgrt (ENSMUSG00000003420) | 82,89 | 1,37 | 0,33 | 4,19 | 0,00 | 0,00 |
| Gm17491 (ENSMUSG00000097042) | 31,15 | 1,37 | 0,50 | 2,72 | 0,01 | 0,05 |
| Gm26853 (ENSMUSG00000097230) | 211,85 | 1,37 | 0,37 | 3,74 | 0,00 | 0,00 |
| Per2 (ENSMUSG00000055866) | 303,05 | 1,37 | 0,45 | 3,06 | 0,00 | 0,02 |
| Tg (ENSMUSG00000053469) | 2062,87 | 1,35 | 0,29 | 4,59 | 0,00 | 0,00 |

| | | | | | | |
|---|---|---|---|---|---|---|
| 9230114K14Rik (ENSMUSG00000097145) | 61,03 | 1,34 | 0,48 | 2,78 | 0,01 | 0,04 |
| Gpc1 (ENSMUSG00000034220) | 1631,98 | 1,34 | 0,27 | 4,99 | 0,00 | 0,00 |
| Gm17655 (ENSMUSG00000090963) | 115,12 | 1,34 | 0,47 | 2,85 | 0,00 | 0,04 |
| Pgm2l1 (ENSMUSG00000030729) | 127,12 | 1,34 | 0,44 | 3,02 | 0,00 | 0,02 |
| Cdkn1a (ENSMUSG00000023067) | 2845,02 | 1,33 | 0,38 | 3,50 | 0,00 | 0,01 |
| Zmym2 (ENSMUSG00000021945) | 1028,25 | 1,33 | 0,28 | 4,75 | 0,00 | 0,00 |
| Pex26 (ENSMUSG00000067825) | 158,33 | 1,33 | 0,46 | 2,90 | 0,00 | 0,03 |
| Zfyve21 (ENSMUSG00000021286) | 67,71 | 1,32 | 0,28 | 4,66 | 0,00 | 0,00 |
| Gm17334 (ENSMUSG00000091191) | 791,31 | 1,32 | 0,33 | 3,96 | 0,00 | 0,00 |
| Spire1 (ENSMUSG00000024533) | 151,51 | 1,31 | 0,46 | 2,85 | 0,00 | 0,04 |
| Il7r (ENSMUSG00000003882) | 1208,36 | 1,30 | 0,30 | 4,36 | 0,00 | 0,00 |
| Ccl5 (ENSMUSG00000035042) | 27817,21 | 1,29 | 0,23 | 5,62 | 0,00 | 0,00 |
| Trav14d-2 (ENSMUSG00000095560) | 16,61 | 1,28 | 0,45 | 2,87 | 0,00 | 0,04 |
| Gm45153 (ENSMUSG00000108693) | 46,43 | 1,28 | 0,35 | 3,63 | 0,00 | 0,00 |
| Magef1 (ENSMUSG00000116632) | 97,67 | 1,28 | 0,44 | 2,92 | 0,00 | 0,03 |
| Cd69 (ENSMUSG00000030156) | 7971,77 | 1,27 | 0,21 | 5,94 | 0,00 | 0,00 |
| Phlda3 (ENSMUSG00000041801) | 865,88 | 1,27 | 0,39 | 3,28 | 0,00 | 0,01 |
| Arl4c (ENSMUSG00000049866) | 410,47 | 1,27 | 0,25 | 5,15 | 0,00 | 0,00 |
| Zfp58 (ENSMUSG00000071291) | 99,46 | 1,26 | 0,38 | 3,32 | 0,00 | 0,01 |
| Zfpm1 (ENSMUSG00000049577) | 1109,30 | 1,26 | 0,27 | 4,68 | 0,00 | 0,00 |
| Furin (ENSMUSG00000030530) | 3164,30 | 1,26 | 0,33 | 3,85 | 0,00 | 0,00 |
| Gadd45b (ENSMUSG00000015312) | 6963,98 | 1,25 | 0,22 | 5,77 | 0,00 | 0,00 |
| Map3k14 (ENSMUSG00000020941) | 1331,45 | 1,25 | 0,31 | 4,06 | 0,00 | 0,00 |
| Gm49701 (ENSMUSG00000116579) | 93,28 | 1,25 | 0,32 | 3,94 | 0,00 | 0,00 |
| Brwd3 (ENSMUSG00000063663) | 92,49 | 1,24 | 0,43 | 2,89 | 0,00 | 0,03 |
| Lncpint (ENSMUSG00000044471) | 89,29 | 1,24 | 0,33 | 3,79 | 0,00 | 0,00 |
| Gm49339 (ENSMUSG00000062593) | 1016,65 | 1,23 | 0,23 | 5,42 | 0,00 | 0,00 |
| Egr3 (ENSMUSG00000033730) | 1179,39 | 1,23 | 0,36 | 3,43 | 0,00 | 0,01 |
| Tada2b (ENSMUSG00000029196) | 110,83 | 1,22 | 0,39 | 3,12 | 0,00 | 0,02 |
| Plcb3 (ENSMUSG00000024960) | 341,97 | 1,22 | 0,38 | 3,22 | 0,00 | 0,01 |
| Uhrf1bp1 (ENSMUSG00000039512) | 155,60 | 1,21 | 0,37 | 3,27 | 0,00 | 0,01 |
| Pim1 (ENSMUSG00000024014) | 3408,90 | 1,21 | 0,36 | 3,31 | 0,00 | 0,01 |
| Psrc1 (ENSMUSG00000068744) | 413,59 | 1,20 | 0,35 | 3,46 | 0,00 | 0,01 |
| Smad3 (ENSMUSG00000032402) | 1197,63 | 1,20 | 0,36 | 3,30 | 0,00 | 0,01 |
| Nfkbiz (ENSMUSG00000035356) | 6154,07 | 1,18 | 0,20 | 6,01 | 0,00 | 0,00 |
| Tom1l2 (ENSMUSG00000000538) | 212,03 | 1,18 | 0,40 | 2,96 | 0,00 | 0,03 |
| Gm29719 (ENSMUSG00000116719) | 36,74 | 1,17 | 0,41 | 2,88 | 0,00 | 0,03 |
| Tnfsf14 (ENSMUSG00000005824) | 956,32 | 1,17 | 0,33 | 3,58 | 0,00 | 0,00 |
| Slc19a2 (ENSMUSG00000040918) | 836,55 | 1,17 | 0,27 | 4,34 | 0,00 | 0,00 |
| Tnfrsf26 (ENSMUSG00000045362) | 606,82 | 1,17 | 0,28 | 4,10 | 0,00 | 0,00 |
| Slc41a1 (ENSMUSG00000013275) | 1571,49 | 1,16 | 0,33 | 3,50 | 0,00 | 0,01 |
| H2-Q10 (ENSMUSG00000067235) | 1075,38 | 1,16 | 0,14 | 8,47 | 0,00 | 0,00 |
| Satb1 (ENSMUSG00000023927) | 20576,10 | 1,15 | 0,18 | 6,22 | 0,00 | 0,00 |
| Arap2 (ENSMUSG00000037999) | 6570,92 | 1,14 | 0,12 | 9,30 | 0,00 | 0,00 |
| Gm8818 (ENSMUSG00000079138) | 143,18 | 1,14 | 0,37 | 3,10 | 0,00 | 0,02 |
| Gm16062 (ENSMUSG00000087249) | 134,43 | 1,14 | 0,29 | 3,92 | 0,00 | 0,00 |
| Plcxd2 (ENSMUSG00000087141) | 937,55 | 1,13 | 0,30 | 3,79 | 0,00 | 0,00 |
| Amigo1 (ENSMUSG00000050947) | 146,79 | 1,13 | 0,40 | 2,83 | 0,00 | 0,04 |
| Icam5 (ENSMUSG00000032174) | 111,76 | 1,13 | 0,40 | 2,82 | 0,00 | 0,04 |
| Tagap (ENSMUSG00000033450) | 12012,73 | 1,13 | 0,19 | 6,08 | 0,00 | 0,00 |
| Gm44873 (ENSMUSG00000109105) | 36,13 | 1,12 | 0,35 | 3,20 | 0,00 | 0,01 |
| Usp32 (ENSMUSG00000000804) | 255,88 | 1,12 | 0,40 | 2,83 | 0,00 | 0,04 |

| | | | | | | |
|---|---|---|---|---|---|---|
| Mylip (ENSMUSG00000038175) | 278,90 | 1,11 | 0,33 | 3,39 | 0,00 | 0,01 |
| Crem (ENSMUSG00000063889) | 2665,41 | 1,11 | 0,24 | 4,54 | 0,00 | 0,00 |
| Gm17041 (ENSMUSG00000090570) | 180,89 | 1,11 | 0,25 | 4,45 | 0,00 | 0,00 |
| Prnp (ENSMUSG00000079037) | 165,62 | 1,11 | 0,39 | 2,86 | 0,00 | 0,04 |
| Gm26732 (ENSMUSG00000097344) | 85,71 | 1,10 | 0,36 | 3,07 | 0,00 | 0,02 |
| Cd7 (ENSMUSG00000025163) | 1339,84 | 1,09 | 0,22 | 4,86 | 0,00 | 0,00 |
| Fosb (ENSMUSG00000003545) | 1221,65 | 1,09 | 0,26 | 4,23 | 0,00 | 0,00 |
| Frat1 (ENSMUSG00000067199) | 104,87 | 1,08 | 0,28 | 3,88 | 0,00 | 0,00 |
| Xcr1 (ENSMUSG00000060509) | 216,27 | 1,08 | 0,34 | 3,14 | 0,00 | 0,02 |
| Rora (ENSMUSG00000032238) | 315,64 | 1,07 | 0,36 | 3,03 | 0,00 | 0,02 |
| Rab20 (ENSMUSG00000031504) | 118,27 | 1,07 | 0,39 | 2,73 | 0,01 | 0,05 |
| Rab6b (ENSMUSG00000032549) | 392,18 | 1,07 | 0,20 | 5,25 | 0,00 | 0,00 |
| Cxcr4 (ENSMUSG00000045382) | 415,51 | 1,07 | 0,26 | 4,17 | 0,00 | 0,00 |
| Mirt1 (ENSMUSG00000097636) | 323,77 | 1,06 | 0,28 | 3,82 | 0,00 | 0,00 |
| Lilr4b (ENSMUSG00000112023) | 2479,36 | 1,06 | 0,25 | 4,26 | 0,00 | 0,00 |
| Cox6b2 (ENSMUSG00000051811) | 74,69 | 1,06 | 0,37 | 2,85 | 0,00 | 0,04 |
| Clcf1 (ENSMUSG00000040663) | 204,25 | 1,05 | 0,27 | 3,84 | 0,00 | 0,00 |
| Mex3d (ENSMUSG00000048696) | 157,38 | 1,05 | 0,30 | 3,52 | 0,00 | 0,01 |
| Maff (ENSMUSG00000042622) | 284,65 | 1,04 | 0,28 | 3,72 | 0,00 | 0,00 |
| Ssh2 (ENSMUSG00000037926) | 3481,76 | 1,04 | 0,26 | 3,99 | 0,00 | 0,00 |
| Errfi1 (ENSMUSG00000028967) | 782,30 | 1,03 | 0,24 | 4,31 | 0,00 | 0,00 |
| Trp53inp1 (ENSMUSG00000028211) | 1682,89 | 1,03 | 0,25 | 4,14 | 0,00 | 0,00 |
| Trdv4 (ENSMUSG00000076867) | 73,87 | 1,02 | 0,36 | 2,82 | 0,00 | 0,04 |
| Brd1 (ENSMUSG00000022387) | 548,06 | 1,02 | 0,30 | 3,37 | 0,00 | 0,01 |
| Csrnp2 (ENSMUSG00000044636) | 219,34 | 1,02 | 0,35 | 2,93 | 0,00 | 0,03 |
| Gm19261 (ENSMUSG00000100529) | 101,29 | 1,02 | 0,33 | 3,05 | 0,00 | 0,02 |
| Lrp11 (ENSMUSG00000019796) | 108,79 | 1,01 | 0,26 | 3,84 | 0,00 | 0,00 |
| Ifitm2 (ENSMUSG00000060591) | 728,67 | 1,01 | 0,37 | 2,75 | 0,01 | 0,05 |
| Dgat1 (ENSMUSG00000022555) | 1171,60 | 1,01 | 0,30 | 3,36 | 0,00 | 0,01 |
| Zc3h12a (ENSMUSG00000042677) | 1034,62 | 1,01 | 0,31 | 3,27 | 0,00 | 0,01 |
| Nfkbid (ENSMUSG00000036931) | 8087,79 | 1,00 | 0,26 | 3,91 | 0,00 | 0,00 |
| Gm9776 (ENSMUSG00000042857) | 69,09 | 0,99 | 0,29 | 3,42 | 0,00 | 0,01 |
| Acss2 (ENSMUSG00000027605) | 993,16 | 0,99 | 0,17 | 5,72 | 0,00 | 0,00 |
| Sipa1l1 (ENSMUSG00000042700) | 2103,85 | 0,99 | 0,28 | 3,58 | 0,00 | 0,01 |
| Ifitm3 (ENSMUSG00000025492) | 775,89 | 0,99 | 0,31 | 3,16 | 0,00 | 0,02 |
| Zxda (ENSMUSG00000073060) | 301,41 | 0,99 | 0,31 | 3,16 | 0,00 | 0,02 |
| Gm43672 (ENSMUSG00000106019) | 108,61 | 0,98 | 0,32 | 3,04 | 0,00 | 0,02 |
| Bcl6 (ENSMUSG00000022508) | 734,16 | 0,98 | 0,29 | 3,40 | 0,00 | 0,01 |
| Dglucy (ENSMUSG00000021185) | 550,71 | 0,98 | 0,30 | 3,29 | 0,00 | 0,01 |
| Adarb1 (ENSMUSG00000020262) | 197,04 | 0,98 | 0,30 | 3,27 | 0,00 | 0,01 |
| Otud1 (ENSMUSG00000043415) | 427,47 | 0,97 | 0,24 | 4,03 | 0,00 | 0,00 |
| Fpgt (ENSMUSG00000053870) | 290,16 | 0,97 | 0,25 | 3,89 | 0,00 | 0,00 |
| Gm26804 (ENSMUSG00000097263) | 160,00 | 0,96 | 0,30 | 3,25 | 0,00 | 0,01 |
| Frmd4b (ENSMUSG00000030064) | 1023,38 | 0,96 | 0,19 | 5,14 | 0,00 | 0,00 |
| Dock5 (ENSMUSG00000044447) | 435,52 | 0,95 | 0,28 | 3,45 | 0,00 | 0,01 |
| Blcap (ENSMUSG00000067787) | 328,67 | 0,95 | 0,31 | 3,10 | 0,00 | 0,02 |
| Trib1 (ENSMUSG00000032501) | 885,16 | 0,95 | 0,21 | 4,60 | 0,00 | 0,00 |
| Ifngr1 (ENSMUSG00000020009) | 18723,51 | 0,95 | 0,17 | 5,45 | 0,00 | 0,00 |
| Chsy1 (ENSMUSG00000032640) | 1684,45 | 0,95 | 0,17 | 5,62 | 0,00 | 0,00 |
| Slmapos2 (ENSMUSG00000090104) | 119,77 | 0,95 | 0,21 | 4,45 | 0,00 | 0,00 |
| Dnajb2 (ENSMUSG00000026203) | 417,72 | 0,94 | 0,34 | 2,81 | 0,00 | 0,04 |
| Cxcl10 (ENSMUSG00000034855) | 533,44 | 0,94 | 0,34 | 2,78 | 0,01 | 0,04 |

| | | | | | | |
|---|---|---|---|---|---|---|
| Fam210b (ENSMUSG00000027495) | 578,75 | 0,94 | 0,27 | 3,49 | 0,00 | 0,01 |
| Irs2 (ENSMUSG00000038894) | 5121,21 | 0,94 | 0,19 | 5,07 | 0,00 | 0,00 |
| Snhg15 (ENSMUSG00000085156) | 358,55 | 0,94 | 0,26 | 3,60 | 0,00 | 0,00 |
| Dtx1 (ENSMUSG00000029603) | 849,52 | 0,94 | 0,32 | 2,92 | 0,00 | 0,03 |
| Il12rb2 (ENSMUSG00000018341) | 7510,41 | 0,94 | 0,27 | 3,48 | 0,00 | 0,01 |
| Tnfrsf12a (ENSMUSG00000023905) | 179,61 | 0,94 | 0,27 | 3,41 | 0,00 | 0,01 |
| Cul3 (ENSMUSG00000004364) | 1234,24 | 0,93 | 0,30 | 3,15 | 0,00 | 0,02 |
| Gm37939 (ENSMUSG00000102725) | 149,35 | 0,93 | 0,28 | 3,32 | 0,00 | 0,01 |
| Trav14-1 (ENSMUSG00000076840) | 9117,74 | 0,93 | 0,14 | 6,50 | 0,00 | 0,00 |
| E430018J23Rik (ENSMUSG00000078580) | 140,19 | 0,93 | 0,31 | 2,97 | 0,00 | 0,03 |
| Gm45552 (ENSMUSG00000110279) | 210,01 | 0,92 | 0,33 | 2,83 | 0,00 | 0,04 |
| 4833407H14Rik (ENSMUSG00000097779) | 213,53 | 0,92 | 0,33 | 2,81 | 0,00 | 0,04 |
| Gm12505 (ENSMUSG00000087070) | 67,23 | 0,92 | 0,27 | 3,36 | 0,00 | 0,01 |
| Relb (ENSMUSG00000002983) | 3740,44 | 0,92 | 0,19 | 4,85 | 0,00 | 0,00 |
| Vps37b (ENSMUSG00000066278) | 11182,45 | 0,92 | 0,17 | 5,44 | 0,00 | 0,00 |
| Dop1a (ENSMUSG00000034973) | 295,36 | 0,91 | 0,26 | 3,53 | 0,00 | 0,01 |
| Bcl3 (ENSMUSG00000053175) | 1868,47 | 0,91 | 0,22 | 4,18 | 0,00 | 0,00 |
| Kcnq1ot1 (ENSMUSG00000101609) | 293,25 | 0,91 | 0,30 | 3,04 | 0,00 | 0,02 |
| Sun2 (ENSMUSG00000042524) | 514,64 | 0,90 | 0,23 | 3,89 | 0,00 | 0,00 |
| Smad7 (ENSMUSG00000025880) | 856,93 | 0,90 | 0,26 | 3,52 | 0,00 | 0,01 |
| Egr2 (ENSMUSG00000037868) | 2648,82 | 0,90 | 0,22 | 4,07 | 0,00 | 0,00 |
| Arl5b (ENSMUSG00000017418) | 302,29 | 0,90 | 0,19 | 4,67 | 0,00 | 0,00 |
| Malat1 (ENSMUSG00000092341) | 5304,68 | 0,90 | 0,20 | 4,51 | 0,00 | 0,00 |
| Ccr2 (ENSMUSG00000049103) | 2699,92 | 0,89 | 0,26 | 3,39 | 0,00 | 0,01 |
| Gm2682 (ENSMUSG00000115681) | 450,31 | 0,89 | 0,20 | 4,55 | 0,00 | 0,00 |
| P2rx4 (ENSMUSG00000029470) | 402,09 | 0,89 | 0,28 | 3,23 | 0,00 | 0,01 |
| Ccdc6 (ENSMUSG00000048701) | 314,93 | 0,89 | 0,29 | 3,06 | 0,00 | 0,02 |
| Fam160a2 (ENSMUSG00000044465) | 893,09 | 0,89 | 0,19 | 4,68 | 0,00 | 0,00 |
| Zfp748 (ENSMUSG00000095432) | 373,30 | 0,88 | 0,27 | 3,25 | 0,00 | 0,01 |
| Pogk (ENSMUSG00000040596) | 1059,76 | 0,88 | 0,32 | 2,75 | 0,01 | 0,05 |
| Tsc22d2 (ENSMUSG00000027806) | 1256,62 | 0,87 | 0,20 | 4,35 | 0,00 | 0,00 |
| Phf1 (ENSMUSG00000024193) | 520,85 | 0,87 | 0,21 | 4,23 | 0,00 | 0,00 |
| Gpr132 (ENSMUSG00000021298) | 1406,09 | 0,87 | 0,24 | 3,59 | 0,00 | 0,00 |
| Cish (ENSMUSG00000032578) | 6516,97 | 0,87 | 0,19 | 4,60 | 0,00 | 0,00 |
| S1pr2 (ENSMUSG00000043895) | 659,86 | 0,86 | 0,18 | 4,67 | 0,00 | 0,00 |
| Myo9a (ENSMUSG00000039585) | 249,16 | 0,86 | 0,30 | 2,88 | 0,00 | 0,03 |
| Tgtp1 (ENSMUSG00000078922) | 5179,49 | 0,86 | 0,18 | 4,79 | 0,00 | 0,00 |
| Rcn3 (ENSMUSG00000019539) | 754,48 | 0,85 | 0,15 | 5,87 | 0,00 | 0,00 |
| Dhx37 (ENSMUSG00000029480) | 1535,37 | 0,85 | 0,24 | 3,55 | 0,00 | 0,01 |
| Cd274 (ENSMUSG00000016496) | 1137,58 | 0,85 | 0,16 | 5,34 | 0,00 | 0,00 |
| Rtn4rl1 (ENSMUSG00000045287) | 338,63 | 0,85 | 0,26 | 3,23 | 0,00 | 0,01 |
| Zfp513 (ENSMUSG00000043059) | 315,95 | 0,85 | 0,23 | 3,61 | 0,00 | 0,00 |
| Dennd5a (ENSMUSG00000035901) | 606,43 | 0,85 | 0,26 | 3,24 | 0,00 | 0,01 |
| Ric8b (ENSMUSG00000035620) | 159,23 | 0,84 | 0,29 | 2,94 | 0,00 | 0,03 |
| Slco3a1 (ENSMUSG00000025790) | 1183,67 | 0,84 | 0,26 | 3,29 | 0,00 | 0,01 |
| Dusp1 (ENSMUSG00000024190) | 11155,75 | 0,84 | 0,13 | 6,59 | 0,00 | 0,00 |
| Sh2b3 (ENSMUSG00000042594) | 1048,84 | 0,84 | 0,25 | 3,30 | 0,00 | 0,01 |
| Rundc3b (ENSMUSG00000040570) | 490,49 | 0,84 | 0,27 | 3,15 | 0,00 | 0,02 |
| Tob1 (ENSMUSG00000037573) | 1971,60 | 0,83 | 0,19 | 4,44 | 0,00 | 0,00 |
| Gm49272 (ENSMUSG00000115329) | 105,57 | 0,83 | 0,30 | 2,80 | 0,01 | 0,04 |
| Gm20186 (ENSMUSG00000106874) | 2375,77 | 0,83 | 0,20 | 4,18 | 0,00 | 0,00 |
| Tnfaip3 (ENSMUSG00000019850) | 34928,44 | 0,83 | 0,15 | 5,57 | 0,00 | 0,00 |

| | | | | | | |
|---|---|---|---|---|---|---|
| Plk3 (ENSMUSG00000028680) | 926,50 | 0,82 | 0,16 | 5,22 | 0,00 | 0,00 |
| Nr4a1 (ENSMUSG00000023034) | 26311,84 | 0,82 | 0,20 | 4,04 | 0,00 | 0,00 |
| Klhl24 (ENSMUSG00000062901) | 787,25 | 0,82 | 0,28 | 2,96 | 0,00 | 0,03 |
| Nmrk1 (ENSMUSG00000037847) | 470,73 | 0,82 | 0,24 | 3,45 | 0,00 | 0,01 |
| Btg1 (ENSMUSG00000036478) | 23342,42 | 0,81 | 0,14 | 5,96 | 0,00 | 0,00 |
| E430024P14Rik (ENSMUSG00000090069) | 247,61 | 0,81 | 0,21 | 3,92 | 0,00 | 0,00 |
| 5031425E22Rik (ENSMUSG00000073147) | 229,91 | 0,81 | 0,28 | 2,90 | 0,00 | 0,03 |
| Sgk1 (ENSMUSG00000019970) | 1314,71 | 0,81 | 0,16 | 5,05 | 0,00 | 0,00 |
| Dusp5 (ENSMUSG00000034765) | 27777,33 | 0,81 | 0,17 | 4,82 | 0,00 | 0,00 |
| Zfp36 (ENSMUSG00000044786) | 9119,25 | 0,81 | 0,16 | 5,15 | 0,00 | 0,00 |
| Tasor2 (ENSMUSG00000033799) | 642,13 | 0,81 | 0,22 | 3,76 | 0,00 | 0,00 |
| Ldlrap1 (ENSMUSG00000037295) | 955,36 | 0,81 | 0,22 | 3,68 | 0,00 | 0,00 |
| Arrdc3 (ENSMUSG00000074794) | 345,16 | 0,81 | 0,25 | 3,30 | 0,00 | 0,01 |
| Birc3 (ENSMUSG00000032000) | 1806,35 | 0,81 | 0,15 | 5,27 | 0,00 | 0,00 |
| Plekha1 (ENSMUSG00000040268) | 476,67 | 0,81 | 0,13 | 6,15 | 0,00 | 0,00 |
| Armc7 (ENSMUSG00000057219) | 1288,42 | 0,80 | 0,21 | 3,76 | 0,00 | 0,00 |
| Casc4 (ENSMUSG00000060227) | 280,51 | 0,80 | 0,25 | 3,20 | 0,00 | 0,01 |
| Dnajb9 (ENSMUSG00000014905) | 1176,50 | 0,80 | 0,12 | 6,46 | 0,00 | 0,00 |
| Gm10125 (ENSMUSG00000063087) | 54,50 | 0,80 | 0,25 | 3,21 | 0,00 | 0,01 |
| Kdm5b (ENSMUSG00000042207) | 2384,06 | 0,80 | 0,18 | 4,44 | 0,00 | 0,00 |
| Nfkb1 (ENSMUSG00000028163) | 2017,55 | 0,80 | 0,20 | 3,98 | 0,00 | 0,00 |
| Rapgef1 (ENSMUSG00000039844) | 2503,34 | 0,80 | 0,28 | 2,83 | 0,00 | 0,04 |
| Cxcr6 (ENSMUSG00000048521) | 21992,64 | 0,80 | 0,21 | 3,85 | 0,00 | 0,00 |
| Utf1 (ENSMUSG00000047751) | 1016,19 | 0,80 | 0,23 | 3,42 | 0,00 | 0,01 |
| Taz (ENSMUSG00000009995) | 997,43 | 0,80 | 0,26 | 3,02 | 0,00 | 0,02 |
| Rhoc (ENSMUSG00000002233) | 290,54 | 0,80 | 0,22 | 3,60 | 0,00 | 0,00 |
| Zmat3 (ENSMUSG00000027663) | 439,18 | 0,80 | 0,25 | 3,19 | 0,00 | 0,02 |
| Zfp281 (ENSMUSG00000041483) | 2696,43 | 0,80 | 0,21 | 3,80 | 0,00 | 0,00 |
| Kansl1l (ENSMUSG00000026004) | 112,02 | 0,79 | 0,29 | 2,73 | 0,01 | 0,05 |
| Trbv13-3 (ENSMUSG00000076470) | 4456,51 | 0,79 | 0,18 | 4,29 | 0,00 | 0,00 |
| Tmem37 (ENSMUSG00000050777) | 236,01 | 0,79 | 0,29 | 2,76 | 0,01 | 0,05 |
| H2-T-ps (ENSMUSG00000073405) | 2187,52 | 0,79 | 0,14 | 5,77 | 0,00 | 0,00 |
| Prkx (ENSMUSG00000035725) | 1878,14 | 0,79 | 0,12 | 6,67 | 0,00 | 0,00 |
| Pfkfb3 (ENSMUSG00000026773) | 1048,48 | 0,78 | 0,22 | 3,61 | 0,00 | 0,00 |
| Jun (ENSMUSG00000052684) | 5749,13 | 0,78 | 0,24 | 3,24 | 0,00 | 0,01 |
| Snapc4 (ENSMUSG00000036281) | 809,30 | 0,78 | 0,27 | 2,93 | 0,00 | 0,03 |
| Emb (ENSMUSG00000021728) | 7307,67 | 0,78 | 0,16 | 4,76 | 0,00 | 0,00 |
| Mapkapk3 (ENSMUSG00000032577) | 4429,38 | 0,77 | 0,18 | 4,19 | 0,00 | 0,00 |
| Thap6 (ENSMUSG00000102644) | 202,73 | 0,77 | 0,26 | 3,03 | 0,00 | 0,02 |
| Map3k8 (ENSMUSG00000024235) | 1034,84 | 0,77 | 0,12 | 6,31 | 0,00 | 0,00 |
| Arc (ENSMUSG00000022602) | 587,95 | 0,77 | 0,18 | 4,29 | 0,00 | 0,00 |
| Samhd1 (ENSMUSG00000027639) | 11460,65 | 0,77 | 0,14 | 5,35 | 0,00 | 0,00 |
| AC137853.2 (ENSMUSG00000118210) | 1056,17 | 0,77 | 0,15 | 4,97 | 0,00 | 0,00 |
| Rab21 (ENSMUSG00000020132) | 1096,12 | 0,77 | 0,27 | 2,82 | 0,00 | 0,04 |
| Tent4a (ENSMUSG00000034575) | 433,52 | 0,76 | 0,23 | 3,30 | 0,00 | 0,01 |
| Gm10687 (ENSMUSG00000097617) | 118,58 | 0,76 | 0,22 | 3,39 | 0,00 | 0,01 |
| Dlg1 (ENSMUSG00000022770) | 891,60 | 0,76 | 0,25 | 3,05 | 0,00 | 0,02 |
| Zdhhc18 (ENSMUSG00000037553) | 7292,17 | 0,76 | 0,19 | 4,05 | 0,00 | 0,00 |
| Ubald2 (ENSMUSG00000050628) | 2758,18 | 0,76 | 0,19 | 4,05 | 0,00 | 0,00 |
| Rnf19b (ENSMUSG00000028793) | 916,71 | 0,76 | 0,20 | 3,75 | 0,00 | 0,00 |
| Gm37376 (ENSMUSG00000102349) | 4583,98 | 0,75 | 0,19 | 3,94 | 0,00 | 0,00 |
| Gm26617 (ENSMUSG00000097340) | 184,50 | 0,75 | 0,18 | 4,21 | 0,00 | 0,00 |

| | | | | | | |
|---|---|---|---|---|---|---|
| Vps54 (ENSMUSG00000020128) | 1541,97 | 0,75 | 0,27 | 2,81 | 0,01 | 0,04 |
| Mia3 (ENSMUSG00000056050) | 2655,96 | 0,75 | 0,13 | 5,95 | 0,00 | 0,00 |
| Ehd1 (ENSMUSG00000024772) | 6941,54 | 0,75 | 0,21 | 3,60 | 0,00 | 0,00 |
| Irf2bp2 (ENSMUSG00000051495) | 766,22 | 0,75 | 0,23 | 3,26 | 0,00 | 0,01 |
| Sgms1 (ENSMUSG00000040451) | 664,54 | 0,75 | 0,22 | 3,40 | 0,00 | 0,01 |
| Rgs1 (ENSMUSG00000026358) | 3810,09 | 0,75 | 0,19 | 3,91 | 0,00 | 0,00 |
| Sema7a (ENSMUSG00000038264) | 279,21 | 0,75 | 0,28 | 2,71 | 0,01 | 0,05 |
| Nfkb2 (ENSMUSG00000025225) | 5779,81 | 0,74 | 0,21 | 3,53 | 0,00 | 0,01 |
| Galnt2 (ENSMUSG00000089704) | 2256,44 | 0,74 | 0,14 | 5,25 | 0,00 | 0,00 |
| Gm3052 (ENSMUSG00000101462) | 115,33 | 0,74 | 0,27 | 2,75 | 0,01 | 0,05 |
| Znrf2 (ENSMUSG00000058446) | 725,75 | 0,74 | 0,27 | 2,73 | 0,01 | 0,05 |
| Snx33 (ENSMUSG00000032733) | 342,74 | 0,74 | 0,26 | 2,86 | 0,00 | 0,04 |
| Gm18853 (ENSMUSG00000098934) | 667,23 | 0,74 | 0,20 | 3,72 | 0,00 | 0,00 |
| Msl2 (ENSMUSG00000066415) | 2302,76 | 0,74 | 0,20 | 3,72 | 0,00 | 0,00 |
| Eps15l1 (ENSMUSG00000006276) | 1406,56 | 0,74 | 0,19 | 3,88 | 0,00 | 0,00 |
| Rere (ENSMUSG00000039852) | 373,72 | 0,73 | 0,22 | 3,33 | 0,00 | 0,01 |
| Gm17132 (ENSMUSG00000061331) | 282,89 | 0,73 | 0,22 | 3,34 | 0,00 | 0,01 |
| Zfp869 (ENSMUSG00000054648) | 1567,56 | 0,73 | 0,21 | 3,50 | 0,00 | 0,01 |
| Smurf2 (ENSMUSG00000018363) | 271,40 | 0,73 | 0,26 | 2,82 | 0,00 | 0,04 |
| Mif4gd (ENSMUSG00000020743) | 1618,99 | 0,73 | 0,17 | 4,33 | 0,00 | 0,00 |
| Jak2 (ENSMUSG00000024789) | 3573,71 | 0,73 | 0,15 | 4,81 | 0,00 | 0,00 |
| Ddit3 (ENSMUSG00000025408) | 402,80 | 0,73 | 0,21 | 3,48 | 0,00 | 0,01 |
| E2f5 (ENSMUSG00000027552) | 362,65 | 0,73 | 0,23 | 3,13 | 0,00 | 0,02 |
| Susd6 (ENSMUSG00000021133) | 1625,71 | 0,72 | 0,21 | 3,48 | 0,00 | 0,01 |
| Gm9973 (ENSMUSG00000113175) | 2367,05 | 0,72 | 0,16 | 4,62 | 0,00 | 0,00 |
| H2-Q5 (ENSMUSG00000055413) | 1817,90 | 0,72 | 0,16 | 4,52 | 0,00 | 0,00 |
| Smg5 (ENSMUSG00000001415) | 1322,37 | 0,72 | 0,25 | 2,86 | 0,00 | 0,04 |
| Gm26905 (ENSMUSG00000097695) | 251,41 | 0,72 | 0,21 | 3,49 | 0,00 | 0,01 |
| Gm37324 (ENSMUSG00000104329) | 162,56 | 0,72 | 0,20 | 3,56 | 0,00 | 0,01 |
| Hs6st1 (ENSMUSG00000045216) | 425,13 | 0,71 | 0,20 | 3,66 | 0,00 | 0,00 |
| Calcoco1 (ENSMUSG00000023055) | 884,38 | 0,71 | 0,24 | 2,93 | 0,00 | 0,03 |
| Gpr171 (ENSMUSG00000050075) | 9810,92 | 0,71 | 0,13 | 5,47 | 0,00 | 0,00 |
| 1700025G04Rik (ENSMUSG00000032666) | 367,52 | 0,71 | 0,21 | 3,44 | 0,00 | 0,01 |
| Uspl1 (ENSMUSG00000041264) | 1487,98 | 0,71 | 0,16 | 4,38 | 0,00 | 0,00 |
| Tmem185b (ENSMUSG00000098923) | 970,02 | 0,71 | 0,15 | 4,61 | 0,00 | 0,00 |
| Siah1a (ENSMUSG00000036840) | 256,48 | 0,71 | 0,16 | 4,30 | 0,00 | 0,00 |
| Thada (ENSMUSG00000024251) | 1665,88 | 0,71 | 0,25 | 2,81 | 0,00 | 0,04 |
| Lilrb4a (ENSMUSG00000112148) | 3319,07 | 0,71 | 0,18 | 3,84 | 0,00 | 0,00 |
| Dennd1b (ENSMUSG00000056268) | 1166,81 | 0,71 | 0,17 | 4,12 | 0,00 | 0,00 |
| Peli1 (ENSMUSG00000020134) | 3418,16 | 0,71 | 0,16 | 4,31 | 0,00 | 0,00 |
| AC109138.2 (ENSMUSG00000111640) | 4362,08 | 0,71 | 0,14 | 5,19 | 0,00 | 0,00 |
| Dyrk2 (ENSMUSG00000028630) | 897,37 | 0,71 | 0,20 | 3,55 | 0,00 | 0,01 |
| 1810026B05Rik (ENSMUSG00000101970) | 376,05 | 0,70 | 0,19 | 3,64 | 0,00 | 0,00 |
| Bbs9 (ENSMUSG00000035919) | 573,94 | 0,70 | 0,22 | 3,26 | 0,00 | 0,01 |
| Diaph1 (ENSMUSG00000024456) | 1041,07 | 0,70 | 0,17 | 4,14 | 0,00 | 0,00 |
| Ppp1r13b (ENSMUSG00000021285) | 1282,68 | 0,70 | 0,21 | 3,41 | 0,00 | 0,01 |
| Irf1 (ENSMUSG00000018899) | 19252,25 | 0,70 | 0,17 | 4,10 | 0,00 | 0,00 |
| Cyhr1 (ENSMUSG00000053929) | 1496,32 | 0,70 | 0,23 | 3,02 | 0,00 | 0,02 |
| Gm42566 (ENSMUSG00000104806) | 881,38 | 0,70 | 0,11 | 6,21 | 0,00 | 0,00 |
| Foxk1 (ENSMUSG00000056493) | 310,22 | 0,70 | 0,24 | 2,85 | 0,00 | 0,04 |
| Rbpj (ENSMUSG00000039191) | 5528,49 | 0,69 | 0,14 | 4,79 | 0,00 | 0,00 |
| Ifrd1 (ENSMUSG00000001627) | 1180,14 | 0,69 | 0,24 | 2,87 | 0,00 | 0,04 |

| | | | | | | |
|---|---|---|---|---|---|---|
| N4bp1 (ENSMUSG00000031652) | 845,73 | 0,69 | 0,20 | 3,52 | 0,00 | 0,01 |
| Themis (ENSMUSG00000049109) | 4026,14 | 0,69 | 0,12 | 5,78 | 0,00 | 0,00 |
| Pogz (ENSMUSG00000038902) | 349,55 | 0,69 | 0,23 | 3,04 | 0,00 | 0,02 |
| Zfp1 (ENSMUSG00000055835) | 286,52 | 0,69 | 0,23 | 3,04 | 0,00 | 0,02 |
| Traf1 (ENSMUSG00000026875) | 11096,51 | 0,69 | 0,14 | 5,05 | 0,00 | 0,00 |
| Aff4 (ENSMUSG00000049470) | 819,86 | 0,69 | 0,18 | 3,91 | 0,00 | 0,00 |
| Vgll4 (ENSMUSG00000030315) | 3487,27 | 0,69 | 0,16 | 4,41 | 0,00 | 0,00 |
| Plekhm1 (ENSMUSG00000034247) | 1083,37 | 0,69 | 0,25 | 2,78 | 0,01 | 0,04 |
| Sfn (ENSMUSG00000047281) | 293,02 | 0,68 | 0,22 | 3,09 | 0,00 | 0,02 |
| Pqlc1 (ENSMUSG00000034006) | 411,79 | 0,68 | 0,19 | 3,67 | 0,00 | 0,00 |
| Zfp62 (ENSMUSG00000046311) | 840,69 | 0,68 | 0,20 | 3,38 | 0,00 | 0,01 |
| Gm33104 (ENSMUSG00000110902) | 295,40 | 0,68 | 0,20 | 3,44 | 0,00 | 0,01 |
| Pwwp2a (ENSMUSG00000044950) | 589,70 | 0,68 | 0,20 | 3,30 | 0,00 | 0,01 |
| Gramd3 (ENSMUSG00000001700) | 6144,04 | 0,67 | 0,14 | 4,77 | 0,00 | 0,00 |
| Mepce (ENSMUSG00000029726) | 850,93 | 0,67 | 0,20 | 3,39 | 0,00 | 0,01 |
| Rbm26 (ENSMUSG00000022119) | 1381,68 | 0,67 | 0,20 | 3,42 | 0,00 | 0,01 |
| Egr1 (ENSMUSG00000038418) | 7513,87 | 0,67 | 0,16 | 4,15 | 0,00 | 0,00 |
| Sidt2 (ENSMUSG00000034908) | 13276,12 | 0,67 | 0,16 | 4,24 | 0,00 | 0,00 |
| Setd1b (ENSMUSG00000038384) | 192,91 | 0,67 | 0,20 | 3,36 | 0,00 | 0,01 |
| Btrc (ENSMUSG00000025217) | 576,35 | 0,67 | 0,21 | 3,14 | 0,00 | 0,02 |
| Pkp4 (ENSMUSG00000026991) | 310,05 | 0,66 | 0,16 | 4,14 | 0,00 | 0,00 |
| Sult2b1 (ENSMUSG00000003271) | 694,09 | 0,66 | 0,21 | 3,11 | 0,00 | 0,02 |
| Mex3c (ENSMUSG00000037253) | 805,85 | 0,66 | 0,24 | 2,78 | 0,01 | 0,04 |
| Sidt1 (ENSMUSG00000022696) | 3095,50 | 0,66 | 0,18 | 3,59 | 0,00 | 0,00 |
| Crtc1 (ENSMUSG00000003575) | 553,58 | 0,66 | 0,20 | 3,26 | 0,00 | 0,01 |
| Arl4d (ENSMUSG00000034936) | 825,00 | 0,66 | 0,20 | 3,22 | 0,00 | 0,01 |
| Zfp507 (ENSMUSG00000044452) | 448,16 | 0,66 | 0,21 | 3,17 | 0,00 | 0,02 |
| Man1a (ENSMUSG00000003746) | 3075,35 | 0,65 | 0,17 | 3,88 | 0,00 | 0,00 |
| AC160336.1 (ENSMUSG00000115801) | 13146,75 | 0,65 | 0,10 | 6,52 | 0,00 | 0,00 |
| Lfng (ENSMUSG00000029570) | 5161,55 | 0,65 | 0,20 | 3,20 | 0,00 | 0,01 |
| Myc (ENSMUSG00000022346) | 8101,82 | 0,65 | 0,18 | 3,54 | 0,00 | 0,01 |
| Il4ra (ENSMUSG00000030748) | 4960,97 | 0,65 | 0,23 | 2,82 | 0,00 | 0,04 |
| Tnfrsf9 (ENSMUSG00000028965) | 42318,46 | 0,65 | 0,20 | 3,24 | 0,00 | 0,01 |
| Arhgap12 (ENSMUSG00000041225) | 411,44 | 0,65 | 0,18 | 3,69 | 0,00 | 0,00 |
| Adrb2 (ENSMUSG00000045730) | 734,05 | 0,65 | 0,22 | 2,93 | 0,00 | 0,03 |
| Btg2 (ENSMUSG00000020423) | 6974,25 | 0,65 | 0,11 | 5,83 | 0,00 | 0,00 |
| Slamf1 (ENSMUSG00000015316) | 4705,78 | 0,64 | 0,14 | 4,59 | 0,00 | 0,00 |
| Clk1 (ENSMUSG00000026034) | 1179,76 | 0,64 | 0,20 | 3,20 | 0,00 | 0,01 |
| Zfp51 (ENSMUSG00000023892) | 490,52 | 0,64 | 0,21 | 2,99 | 0,00 | 0,03 |
| Itk (ENSMUSG00000020395) | 13345,47 | 0,64 | 0,17 | 3,76 | 0,00 | 0,00 |
| Lpar5 (ENSMUSG00000067714) | 388,76 | 0,64 | 0,17 | 3,66 | 0,00 | 0,00 |
| Crim1 (ENSMUSG00000024074) | 1330,89 | 0,64 | 0,23 | 2,80 | 0,01 | 0,04 |
| Eif4a2 (ENSMUSG00000022884) | 2020,24 | 0,64 | 0,21 | 2,99 | 0,00 | 0,03 |
| Dyrk1a (ENSMUSG00000022897) | 1200,94 | 0,64 | 0,18 | 3,56 | 0,00 | 0,01 |
| Nr3c1 (ENSMUSG00000024431) | 1973,83 | 0,64 | 0,17 | 3,66 | 0,00 | 0,00 |
| Traf4 (ENSMUSG00000017386) | 2390,74 | 0,64 | 0,17 | 3,75 | 0,00 | 0,00 |
| Vps33b (ENSMUSG00000030534) | 1000,56 | 0,63 | 0,14 | 4,40 | 0,00 | 0,00 |
| Klrc2 (ENSMUSG00000052736) | 282,46 | 0,63 | 0,14 | 4,50 | 0,00 | 0,00 |
| Gpr174 (ENSMUSG00000073008) | 2427,61 | 0,63 | 0,14 | 4,66 | 0,00 | 0,00 |
| Il21r (ENSMUSG00000030745) | 8662,85 | 0,63 | 0,15 | 4,16 | 0,00 | 0,00 |
| Slamf7 (ENSMUSG00000038179) | 6009,99 | 0,63 | 0,15 | 4,18 | 0,00 | 0,00 |
| Ctu1 (ENSMUSG00000038888) | 463,99 | 0,63 | 0,21 | 3,01 | 0,00 | 0,02 |

| | | | | | | |
|---|---|---|---|---|---|---|
| Gm26809 (ENSMUSG00000097815) | 62167,83 | 0,63 | 0,13 | 4,95 | 0,00 | 0,00 |
| Tug1 (ENSMUSG00000056579) | 2322,33 | 0,63 | 0,18 | 3,55 | 0,00 | 0,01 |
| Zbtb11 (ENSMUSG00000022601) | 975,24 | 0,63 | 0,22 | 2,89 | 0,00 | 0,03 |
| Zfp36l2 (ENSMUSG00000045817) | 11262,72 | 0,63 | 0,16 | 4,02 | 0,00 | 0,00 |
| Tgif1 (ENSMUSG00000047407) | 4073,58 | 0,62 | 0,21 | 2,96 | 0,00 | 0,03 |
| Rnf219 (ENSMUSG00000022120) | 424,93 | 0,62 | 0,16 | 3,91 | 0,00 | 0,00 |
| Fas (ENSMUSG00000024778) | 1377,55 | 0,62 | 0,16 | 3,86 | 0,00 | 0,00 |
| Sytl3 (ENSMUSG00000041831) | 2179,02 | 0,62 | 0,14 | 4,48 | 0,00 | 0,00 |
| Mief1 (ENSMUSG00000022412) | 812,83 | 0,61 | 0,21 | 2,92 | 0,00 | 0,03 |
| Tgtp2 (ENSMUSG00000078921) | 2616,84 | 0,61 | 0,14 | 4,33 | 0,00 | 0,00 |
| Gm26616 (ENSMUSG00000102425) | 82,75 | 0,61 | 0,22 | 2,82 | 0,00 | 0,04 |
| Vopp1 (ENSMUSG00000037788) | 1603,91 | 0,61 | 0,14 | 4,50 | 0,00 | 0,00 |
| Man2a2 (ENSMUSG00000038886) | 1725,85 | 0,61 | 0,20 | 3,11 | 0,00 | 0,02 |
| Slc12a4 (ENSMUSG00000017765) | 2255,71 | 0,61 | 0,21 | 2,97 | 0,00 | 0,03 |
| Neurl3 (ENSMUSG00000047180) | 3026,26 | 0,61 | 0,15 | 4,15 | 0,00 | 0,00 |
| Smg7 (ENSMUSG00000042772) | 3497,65 | 0,61 | 0,21 | 2,95 | 0,00 | 0,03 |
| Gbp8 (ENSMUSG00000034438) | 1217,21 | 0,61 | 0,18 | 3,34 | 0,00 | 0,01 |
| Ythdc1 (ENSMUSG00000035851) | 1486,49 | 0,61 | 0,20 | 3,08 | 0,00 | 0,02 |
| Gpr65 (ENSMUSG00000021886) | 2891,99 | 0,61 | 0,12 | 5,05 | 0,00 | 0,00 |
| Klhdc4 (ENSMUSG00000040263) | 2046,35 | 0,61 | 0,14 | 4,33 | 0,00 | 0,00 |
| Fbrs (ENSMUSG00000042423) | 449,81 | 0,61 | 0,13 | 4,82 | 0,00 | 0,00 |
| Flcn (ENSMUSG00000032633) | 1389,14 | 0,61 | 0,13 | 4,65 | 0,00 | 0,00 |
| Gigyf1 (ENSMUSG00000029714) | 394,99 | 0,61 | 0,22 | 2,79 | 0,01 | 0,04 |
| Avl9 (ENSMUSG00000029787) | 891,87 | 0,61 | 0,21 | 2,90 | 0,00 | 0,03 |
| Clk4 (ENSMUSG00000020385) | 914,90 | 0,60 | 0,20 | 3,08 | 0,00 | 0,02 |
| Cpne3 (ENSMUSG00000028228) | 1759,28 | 0,60 | 0,14 | 4,43 | 0,00 | 0,00 |
| Larp1 (ENSMUSG00000037331) | 7218,35 | 0,60 | 0,21 | 2,93 | 0,00 | 0,03 |
| Zfp598 (ENSMUSG00000041130) | 661,42 | 0,60 | 0,17 | 3,51 | 0,00 | 0,01 |
| Irgm2 (ENSMUSG00000069874) | 632,24 | 0,60 | 0,17 | 3,55 | 0,00 | 0,01 |
| Map2k3 (ENSMUSG00000018932) | 8461,66 | 0,60 | 0,16 | 3,69 | 0,00 | 0,00 |
| Prmt2 (ENSMUSG00000020230) | 542,52 | 0,60 | 0,18 | 3,36 | 0,00 | 0,01 |
| Map3k2 (ENSMUSG00000024383) | 369,61 | 0,60 | 0,17 | 3,47 | 0,00 | 0,01 |
| Rbm12b1 (ENSMUSG00000046667) | 196,75 | 0,60 | 0,20 | 2,96 | 0,00 | 0,03 |
| AC149090.1 (ENSMUSG00000095041) | 642,85 | 0,60 | 0,17 | 3,42 | 0,00 | 0,01 |
| Uhrf1bp1l (ENSMUSG00000019951) | 2699,42 | 0,60 | 0,19 | 3,11 | 0,00 | 0,02 |
| Stat3 (ENSMUSG00000004040) | 25287,00 | 0,60 | 0,19 | 3,21 | 0,00 | 0,01 |
| Tnfrsf18 (ENSMUSG00000041954) | 11603,53 | 0,60 | 0,19 | 3,19 | 0,00 | 0,01 |
| Ankrd44 (ENSMUSG00000052331) | 4588,12 | 0,60 | 0,20 | 2,94 | 0,00 | 0,03 |
| Pprc1 (ENSMUSG00000055491) | 2052,66 | 0,59 | 0,21 | 2,78 | 0,01 | 0,04 |
| Ptprc (ENSMUSG00000026395) | 57719,48 | 0,59 | 0,11 | 5,25 | 0,00 | 0,00 |
| Zbtb9 (ENSMUSG00000079605) | 484,96 | 0,59 | 0,16 | 3,68 | 0,00 | 0,00 |
| Pik3c3 (ENSMUSG00000033628) | 1395,80 | 0,59 | 0,11 | 5,54 | 0,00 | 0,00 |
| Abcg1 (ENSMUSG00000024030) | 1008,69 | 0,59 | 0,15 | 3,95 | 0,00 | 0,00 |

| | baseMean | log2FoldChange | lfcSE | stat | pvalue | padj |
|---|---|---|---|---|---|---|
| Epdr1 (ENSMUSG00000002808) | 72,70 | -9,58 | 1,01 | -9,50 | 0,00 | 0,00 |
| Chl1 (ENSMUSG00000030077) | 31,36 | -8,37 | 1,49 | -5,63 | 0,00 | 0,00 |
| Tpbgl (ENSMUSG00000096606) | 26,21 | -8,11 | 1,11 | -7,31 | 0,00 | 0,00 |
| Ckm (ENSMUSG00000030399) | 16,50 | -7,44 | 1,21 | -6,15 | 0,00 | 0,00 |
| Adgrd1 (ENSMUSG00000044017) | 16,31 | -7,42 | 1,57 | -4,73 | 0,00 | 0,00 |
| Oit3 (ENSMUSG00000009654) | 14,04 | -7,20 | 1,71 | -4,20 | 0,00 | 0,00 |
| Csf2rb (ENSMUSG00000071713) | 13,59 | -7,16 | 2,18 | -3,28 | 0,00 | 0,01 |
| Ampd3 (ENSMUSG00000005686) | 12,86 | -7,08 | 1,62 | -4,37 | 0,00 | 0,00 |
| Arfgef3 (ENSMUSG00000019852) | 12,52 | -7,04 | 2,18 | -3,23 | 0,00 | 0,01 |
| Gm47571 (ENSMUSG00000112702) | 66,04 | -6,99 | 0,78 | -8,96 | 0,00 | 0,00 |
| Cdh17 (ENSMUSG00000028217) | 11,39 | -6,91 | 2,13 | -3,25 | 0,00 | 0,01 |
| Fbln1 (ENSMUSG00000006369) | 16,53 | -6,81 | 1,52 | -4,47 | 0,00 | 0,00 |
| Cd300c2 (ENSMUSG00000044811) | 10,44 | -6,78 | 1,21 | -5,60 | 0,00 | 0,00 |
| Plppr2 (ENSMUSG00000040563) | 10,35 | -6,76 | 2,22 | -3,04 | 0,00 | 0,02 |
| Csf2rb2 (ENSMUSG00000071714) | 9,93 | -6,71 | 1,79 | -3,76 | 0,00 | 0,00 |
| Gm46519 (ENSMUSG00000115969) | 9,75 | -6,68 | 1,20 | -5,58 | 0,00 | 0,00 |
| Lpcat2 (ENSMUSG00000033192) | 9,50 | -6,64 | 1,67 | -3,99 | 0,00 | 0,00 |
| Klra17 (ENSMUSG00000014543) | 9,04 | -6,57 | 1,34 | -4,91 | 0,00 | 0,00 |
| Rtkn (ENSMUSG00000034930) | 8,76 | -6,52 | 1,68 | -3,88 | 0,00 | 0,00 |
| Gm14377 (ENSMUSG00000109501) | 18,17 | -6,42 | 1,18 | -5,45 | 0,00 | 0,00 |
| Gas6 (ENSMUSG00000031451) | 7,93 | -6,38 | 2,23 | -2,87 | 0,00 | 0,04 |
| Slc40a1 (ENSMUSG00000025993) | 7,87 | -6,37 | 1,90 | -3,35 | 0,00 | 0,01 |
| Gm20667 (ENSMUSG00000093726) | 7,80 | -6,36 | 1,38 | -4,60 | 0,00 | 0,00 |
| Gpr179 (ENSMUSG00000070337) | 7,74 | -6,35 | 1,71 | -3,71 | 0,00 | 0,00 |
| Bco1 (ENSMUSG00000031845) | 26,59 | -6,32 | 0,95 | -6,68 | 0,00 | 0,00 |
| Syna (ENSMUSG00000085957) | 7,37 | -6,28 | 2,27 | -2,76 | 0,01 | 0,04 |
| Gm33424 (ENSMUSG00000113101) | 7,17 | -6,24 | 1,50 | -4,16 | 0,00 | 0,00 |
| Aif1l (ENSMUSG00000001864) | 7,17 | -6,21 | 1,62 | -3,83 | 0,00 | 0,00 |
| Csprs (ENSMUSG00000062783) | 6,59 | -6,12 | 2,25 | -2,72 | 0,01 | 0,05 |
| Pigz (ENSMUSG00000045625) | 6,69 | -6,11 | 1,85 | -3,30 | 0,00 | 0,01 |
| Itgb5 (ENSMUSG00000022817) | 9,82 | -6,10 | 1,58 | -3,86 | 0,00 | 0,00 |
| Igfbp7 (ENSMUSG00000036256) | 95,24 | -6,08 | 1,45 | -4,18 | 0,00 | 0,00 |
| A430028G04Rik (ENSMUSG00000112 | 6,17 | -6,02 | 1,82 | -3,31 | 0,00 | 0,01 |
| Myl10 (ENSMUSG00000005474) | 6,05 | -5,99 | 1,59 | -3,78 | 0,00 | 0,00 |
| Tmcc3 (ENSMUSG00000020023) | 40,91 | -5,97 | 1,02 | -5,84 | 0,00 | 0,00 |
| Gm15287 (ENSMUSG00000086480) | 195,17 | -5,97 | 1,11 | -5,39 | 0,00 | 0,00 |
| Ccdc162 (ENSMUSG00000075225) | 21,73 | -5,93 | 1,91 | -3,11 | 0,00 | 0,02 |
| Rab33a (ENSMUSG00000031104) | 5,75 | -5,92 | 1,29 | -4,58 | 0,00 | 0,00 |
| Cysltr1 (ENSMUSG00000052821) | 5,82 | -5,91 | 2,05 | -2,88 | 0,00 | 0,03 |
| Vcan (ENSMUSG00000021614) | 5,51 | -5,86 | 2,00 | -2,93 | 0,00 | 0,03 |
| Gm156 (ENSMUSG00000071158) | 1410,43 | -5,85 | 0,72 | -8,11 | 0,00 | 0,00 |
| Thsd7b (ENSMUSG00000042581) | 15,23 | -5,84 | 1,13 | -5,16 | 0,00 | 0,00 |
| Nxf2 (ENSMUSG00000009941) | 7,71 | -5,68 | 2,01 | -2,82 | 0,00 | 0,04 |
| Slc17a7 (ENSMUSG00000070570) | 8,03 | -5,33 | 1,66 | -3,22 | 0,00 | 0,01 |
| Trf (ENSMUSG00000032554) | 11,42 | -5,30 | 1,76 | -3,02 | 0,00 | 0,02 |
| Pif1 (ENSMUSG00000041064) | 35,70 | -5,29 | 1,17 | -4,53 | 0,00 | 0,00 |
| Spp1 (ENSMUSG00000029304) | 1558,90 | -5,16 | 0,97 | -5,32 | 0,00 | 0,00 |
| Tns3 (ENSMUSG00000020422) | 7,52 | -5,12 | 1,62 | -3,17 | 0,00 | 0,02 |
| Msr1 (ENSMUSG00000025044) | 23,85 | -5,04 | 1,20 | -4,22 | 0,00 | 0,00 |
| BC035947 (ENSMUSG00000090486) | 33,98 | -4,95 | 0,83 | -5,93 | 0,00 | 0,00 |
| Eldr (ENSMUSG00000087060) | 38,49 | -4,93 | 0,88 | -5,61 | 0,00 | 0,00 |
| Cd22 (ENSMUSG00000030577) | 34,51 | -4,90 | 1,24 | -3,95 | 0,00 | 0,00 |
| Ldlr (ENSMUSG00000032193) | 77,10 | -4,83 | 0,79 | -6,11 | 0,00 | 0,00 |
| Lncbate6 (ENSMUSG00000114665) | 8,69 | -4,75 | 1,58 | -3,02 | 0,00 | 0,02 |
| 2900026A02Rik (ENSMUSG0000005[illegible] | 155,44 | -4,71 | 1,07 | -4,40 | 0,00 | 0,00 |
| Exph5 (ENSMUSG00000034584) | 15,71 | -4,64 | 1,38 | -3,37 | 0,00 | 0,01 |

| | | | | | | |
|---|---|---|---|---|---|---|
| Krt222 (ENSMUSG00000035849) | 8,32 | -4,64 | 1,67 | -2,78 | 0,01 | 0,04 |
| Fam220a (ENSMUSG00000083012) | 12,11 | -4,62 | 1,15 | -4,02 | 0,00 | 0,00 |
| Nek2 (ENSMUSG00000026622) | 664,25 | -4,58 | 1,18 | -3,90 | 0,00 | 0,00 |
| Gucy1b1 (ENSMUSG00000028005) | 24,37 | -4,42 | 1,20 | -3,68 | 0,00 | 0,00 |
| Galnt9 (ENSMUSG00000033316) | 22,59 | -4,41 | 1,18 | -3,74 | 0,00 | 0,00 |
| Kif19a (ENSMUSG00000010021) | 9,47 | -4,40 | 1,41 | -3,12 | 0,00 | 0,02 |
| Adgre1 (ENSMUSG00000004730) | 40,25 | -4,39 | 1,04 | -4,25 | 0,00 | 0,00 |
| Troap (ENSMUSG00000032783) | 117,71 | -4,39 | 0,48 | -9,10 | 0,00 | 0,00 |
| Slc16a11 (ENSMUSG00000040938) | 20,27 | -4,36 | 1,07 | -4,06 | 0,00 | 0,00 |
| Zc3h12b (ENSMUSG00000035045) | 15,72 | -4,32 | 1,35 | -3,21 | 0,00 | 0,01 |
| 4632418H02Rik (ENSMUSG0000011 | 6,01 | -4,28 | 1,40 | -3,06 | 0,00 | 0,02 |
| Angptl2 (ENSMUSG00000004105) | 162,73 | -4,27 | 1,57 | -2,72 | 0,01 | 0,05 |
| Hist1h2af (ENSMUSG00000061991) | 6,31 | -4,27 | 1,20 | -3,55 | 0,00 | 0,01 |
| Hist1h2bm (ENSMUSG00000114279) | 18,24 | -4,23 | 0,74 | -5,73 | 0,00 | 0,00 |
| Hmgn3 (ENSMUSG00000066456) | 15,92 | -4,18 | 0,94 | -4,46 | 0,00 | 0,00 |
| Nrgn (ENSMUSG00000053310) | 1228,10 | -4,12 | 0,39 | -10,61 | 0,00 | 0,00 |
| 1700019D03Rik (ENSMUSG0000004 | 67,72 | -4,12 | 0,66 | -6,20 | 0,00 | 0,00 |
| Mettl5os (ENSMUSG00000085735) | 5,59 | -4,10 | 1,24 | -3,31 | 0,00 | 0,01 |
| Rab7b (ENSMUSG00000052688) | 14,45 | -4,03 | 1,39 | -2,90 | 0,00 | 0,03 |
| Ube2c (ENSMUSG00000001403) | 1223,96 | -4,00 | 0,39 | -10,15 | 0,00 | 0,00 |
| 2810025M15Rik (ENSMUSG00000049 | 51,98 | -3,98 | 0,84 | -4,75 | 0,00 | 0,00 |
| Gm42429 (ENSMUSG00000105373) | 11,34 | -3,92 | 1,33 | -2,95 | 0,00 | 0,03 |
| Sapcd2 (ENSMUSG00000026955) | 283,06 | -3,92 | 1,25 | -3,14 | 0,00 | 0,02 |
| Gm15428 (ENSMUSG00000046057) | 38,65 | -3,89 | 0,87 | -4,49 | 0,00 | 0,00 |
| Ccnb2 (ENSMUSG00000032218) | 1699,27 | -3,79 | 0,35 | -10,87 | 0,00 | 0,00 |
| Cdkn3 (ENSMUSG00000037628) | 203,95 | -3,79 | 0,69 | -5,46 | 0,00 | 0,00 |
| Ska1 (ENSMUSG00000036223) | 218,28 | -3,75 | 0,49 | -7,65 | 0,00 | 0,00 |
| Ccl8 (ENSMUSG00000009185) | 67,38 | -3,73 | 1,06 | -3,52 | 0,00 | 0,01 |
| E2f7 (ENSMUSG00000020185) | 261,95 | -3,73 | 0,95 | -3,93 | 0,00 | 0,00 |
| Cavin3 (ENSMUSG00000037060) | 5,49 | -3,72 | 1,23 | -3,03 | 0,00 | 0,02 |
| Cdc25c (ENSMUSG00000044201) | 156,87 | -3,69 | 0,60 | -6,15 | 0,00 | 0,00 |
| Gm47140 (ENSMUSG00000111325) | 8,62 | -3,69 | 0,91 | -4,06 | 0,00 | 0,00 |
| Gas2l3 (ENSMUSG00000074802) | 123,80 | -3,67 | 0,52 | -7,00 | 0,00 | 0,00 |
| Dbn1 (ENSMUSG00000034675) | 21,80 | -3,67 | 1,02 | -3,59 | 0,00 | 0,00 |
| Anln (ENSMUSG00000036777) | 117,27 | -3,66 | 0,93 | -3,92 | 0,00 | 0,00 |
| Cenpf (ENSMUSG00000026605) | 478,34 | -3,62 | 0,53 | -6,78 | 0,00 | 0,00 |
| Exoc3l (ENSMUSG00000043251) | 35,32 | -3,61 | 0,67 | -5,39 | 0,00 | 0,00 |
| Ankle1 (ENSMUSG00000046295) | 49,05 | -3,61 | 1,12 | -3,21 | 0,00 | 0,01 |
| Gm49364 (ENSMUSG00000115344) | 9,90 | -3,60 | 0,89 | -4,05 | 0,00 | 0,00 |
| Knl1 (ENSMUSG00000027326) | 500,07 | -3,57 | 0,85 | -4,21 | 0,00 | 0,00 |
| Speer6-ps1 (ENSMUSG00000091304) | 31,69 | -3,56 | 0,83 | -4,31 | 0,00 | 0,00 |
| Rab3il1 (ENSMUSG00000024663) | 12,84 | -3,55 | 1,28 | -2,77 | 0,01 | 0,04 |
| Gm49498 (ENSMUSG00000116109) | 10,88 | -3,54 | 1,24 | -2,85 | 0,00 | 0,04 |
| Hmmr (ENSMUSG00000020330) | 697,57 | -3,51 | 0,50 | -7,07 | 0,00 | 0,00 |
| Depdc1a (ENSMUSG00000028175) | 258,58 | -3,50 | 0,95 | -3,68 | 0,00 | 0,00 |
| Cep55 (ENSMUSG00000024989) | 582,99 | -3,45 | 0,32 | -10,91 | 0,00 | 0,00 |
| Ttk (ENSMUSG00000038379) | 260,82 | -3,44 | 0,46 | -7,48 | 0,00 | 0,00 |
| Cldn34c1 (ENSMUSG00000079450) | 5,99 | -3,41 | 1,04 | -3,28 | 0,00 | 0,01 |
| Zkscan8 (ENSMUSG00000063894) | 90,14 | -3,41 | 0,95 | -3,57 | 0,00 | 0,01 |
| Zranb3 (ENSMUSG00000036086) | 181,76 | -3,39 | 0,59 | -5,72 | 0,00 | 0,00 |
| Gm2788 (ENSMUSG00000085995) | 31,89 | -3,37 | 0,68 | -4,97 | 0,00 | 0,00 |
| Mxd3 (ENSMUSG00000021485) | 241,69 | -3,35 | 0,38 | -8,74 | 0,00 | 0,00 |
| Gm38271 (ENSMUSG00000103053) | 21,03 | -3,35 | 0,76 | -4,39 | 0,00 | 0,00 |
| Mki67 (ENSMUSG00000031004) | 1369,09 | -3,34 | 0,36 | -9,38 | 0,00 | 0,00 |
| Birc5 (ENSMUSG00000017716) | 1520,93 | -3,34 | 0,40 | -8,35 | 0,00 | 0,00 |
| Pla2g7 (ENSMUSG00000023913) | 102,11 | -3,33 | 0,87 | -3,81 | 0,00 | 0,00 |
| Depdc1b (ENSMUSG00000021697) | 177,46 | -3,32 | 0,59 | -5,62 | 0,00 | 0,00 |

| | | | | | | |
|---|---|---|---|---|---|---|
| Kif4 (ENSMUSG00000034311) | 755,29 | -3,32 | 0,46 | -7,18 | 0,00 | 0,00 |
| Gm21188 (ENSMUSG00000095609) | 21,04 | -3,31 | 1,19 | -2,78 | 0,01 | 0,04 |
| Rbpj-ps3 (ENSMUSG00000079575) | 483,85 | -3,30 | 0,23 | -14,60 | 0,00 | 0,00 |
| Ddah2 (ENSMUSG00000007039) | 39,70 | -3,26 | 0,98 | -3,32 | 0,00 | 0,01 |
| Cntn1 (ENSMUSG00000055022) | 9,94 | -3,25 | 1,09 | -2,97 | 0,00 | 0,03 |
| Tmie (ENSMUSG00000049555) | 35,54 | -3,23 | 1,15 | -2,82 | 0,00 | 0,04 |
| Tpx2 (ENSMUSG00000027469) | 1400,06 | -3,23 | 0,34 | -9,54 | 0,00 | 0,00 |
| Dnah11 (ENSMUSG00000018581) | 18,27 | -3,22 | 1,19 | -2,71 | 0,01 | 0,05 |
| Cdca3 (ENSMUSG00000023505) | 833,51 | -3,21 | 0,48 | -6,68 | 0,00 | 0,00 |
| Bub1 (ENSMUSG00000027379) | 843,42 | -3,20 | 0,65 | -4,93 | 0,00 | 0,00 |
| E2f8 (ENSMUSG00000046179) | 310,86 | -3,19 | 0,51 | -6,25 | 0,00 | 0,00 |
| Kif2c (ENSMUSG00000028678) | 557,11 | -3,18 | 0,52 | -6,18 | 0,00 | 0,00 |
| Hist1h1b (ENSMUSG00000058773) | 112,47 | -3,16 | 0,42 | -7,58 | 0,00 | 0,00 |
| A930002I21Rik (ENSMUSG000000501 | 94,50 | -3,14 | 0,88 | -3,58 | 0,00 | 0,00 |
| Mcoln3 (ENSMUSG00000036853) | 80,23 | -3,14 | 0,88 | -3,59 | 0,00 | 0,00 |
| Aurkb (ENSMUSG00000020897) | 1257,39 | -3,13 | 0,27 | -11,51 | 0,00 | 0,00 |
| Rad51c (ENSMUSG00000007646) | 83,73 | -3,12 | 0,93 | -3,33 | 0,00 | 0,01 |
| Ccna2 (ENSMUSG00000027715) | 2544,06 | -3,11 | 0,38 | -8,14 | 0,00 | 0,00 |
| Hist1h2ae (ENSMUSG00000069272) | 217,14 | -3,09 | 0,43 | -7,10 | 0,00 | 0,00 |
| Parpbp (ENSMUSG00000035365) | 135,67 | -3,09 | 0,44 | -6,96 | 0,00 | 0,00 |
| Nuf2 (ENSMUSG00000026683) | 615,13 | -3,08 | 0,36 | -8,55 | 0,00 | 0,00 |
| Rtkn2 (ENSMUSG00000037846) | 16,40 | -3,08 | 0,95 | -3,23 | 0,00 | 0,01 |
| Sgo1 (ENSMUSG00000023940) | 252,36 | -3,04 | 0,34 | -8,96 | 0,00 | 0,00 |
| Pimreg (ENSMUSG00000020808) | 262,63 | -3,03 | 0,89 | -3,41 | 0,00 | 0,01 |
| Pbk (ENSMUSG00000022033) | 277,65 | -3,02 | 0,58 | -5,21 | 0,00 | 0,00 |
| Rasgef1c (ENSMUSG00000020374) | 9,28 | -3,00 | 1,01 | -2,97 | 0,00 | 0,03 |
| Prr11 (ENSMUSG00000020493) | 107,68 | -2,99 | 0,67 | -4,47 | 0,00 | 0,00 |
| Mis18bp1 (ENSMUSG00000047534) | 260,16 | -2,98 | 0,54 | -5,49 | 0,00 | 0,00 |
| Nusap1 (ENSMUSG00000027306) | 1345,64 | -2,98 | 0,28 | -10,55 | 0,00 | 0,00 |
| Copz2 (ENSMUSG00000018672) | 14,89 | -2,97 | 1,02 | -2,91 | 0,00 | 0,03 |
| Pclaf (ENSMUSG00000040204) | 3484,06 | -2,96 | 0,47 | -6,31 | 0,00 | 0,00 |
| Cenpi (ENSMUSG00000031262) | 246,71 | -2,95 | 0,61 | -4,85 | 0,00 | 0,00 |
| Esco2 (ENSMUSG00000022034) | 292,03 | -2,94 | 0,42 | -6,97 | 0,00 | 0,00 |
| Clspn (ENSMUSG00000042489) | 294,72 | -2,94 | 0,50 | -5,84 | 0,00 | 0,00 |
| Serpinf1 (ENSMUSG00000000753) | 28,02 | -2,92 | 0,91 | -3,21 | 0,00 | 0,01 |
| Mybl2 (ENSMUSG00000017861) | 1138,84 | -2,91 | 0,67 | -4,34 | 0,00 | 0,00 |
| Shcbp1 (ENSMUSG00000022322) | 706,94 | -2,90 | 0,30 | -9,79 | 0,00 | 0,00 |
| Hist1h2ab (ENSMUSG00000061615) | 78,19 | -2,87 | 0,99 | -2,89 | 0,00 | 0,03 |
| Mlf1 (ENSMUSG00000048416) | 43,06 | -2,85 | 0,59 | -4,86 | 0,00 | 0,00 |
| Hist1h2ap (ENSMUSG00000094777) | 6704,30 | -2,84 | 0,34 | -8,38 | 0,00 | 0,00 |
| Ugt1a7c (ENSMUSG00000090124) | 183,28 | -2,82 | 0,44 | -6,47 | 0,00 | 0,00 |
| Hist1h3c (ENSMUSG00000069310) | 65,44 | -2,82 | 0,59 | -4,77 | 0,00 | 0,00 |
| Iqgap3 (ENSMUSG00000028068) | 139,70 | -2,81 | 0,42 | -6,74 | 0,00 | 0,00 |
| Cenpe (ENSMUSG00000045328) | 662,17 | -2,81 | 0,39 | -7,27 | 0,00 | 0,00 |
| Hist1h2bp (ENSMUSG00000069308) | 8,57 | -2,81 | 0,98 | -2,85 | 0,00 | 0,04 |
| Ccnb1 (ENSMUSG00000041431) | 1175,43 | -2,80 | 0,32 | -8,68 | 0,00 | 0,00 |
| Cdc20 (ENSMUSG00000006398) | 1235,76 | -2,80 | 0,38 | -7,36 | 0,00 | 0,00 |
| Gucy1a1 (ENSMUSG00000033910) | 278,90 | -2,80 | 0,50 | -5,54 | 0,00 | 0,00 |
| Ckap2l (ENSMUSG00000048327) | 674,71 | -2,79 | 0,39 | -7,12 | 0,00 | 0,00 |
| Melk (ENSMUSG00000035683) | 424,42 | -2,78 | 0,39 | -7,10 | 0,00 | 0,00 |
| Cd244a (ENSMUSG00000004709) | 247,99 | -2,77 | 0,36 | -7,64 | 0,00 | 0,00 |
| Hist1h2bj (ENSMUSG00000069300) | 80,89 | -2,77 | 0,37 | -7,53 | 0,00 | 0,00 |
| Cdk1 (ENSMUSG00000019942) | 1039,01 | -2,77 | 0,29 | -9,52 | 0,00 | 0,00 |
| Kif11 (ENSMUSG00000012443) | 1498,94 | -2,75 | 0,53 | -5,21 | 0,00 | 0,00 |
| Stil (ENSMUSG00000028718) | 527,38 | -2,74 | 0,36 | -7,60 | 0,00 | 0,00 |
| Plk1 (ENSMUSG00000030867) | 900,16 | -2,74 | 0,36 | -7,56 | 0,00 | 0,00 |
| Spc24 (ENSMUSG00000074476) | 851,35 | -2,73 | 0,45 | -6,14 | 0,00 | 0,00 |

| | | | | | | |
|---|---|---|---|---|---|---|
| Foxm1 (ENSMUSG00000001517) | 771,29 | -2,73 | 0,26 | -10,55 | 0,00 | 0,00 |
| Tmod1 (ENSMUSG00000028328) | 100,85 | -2,71 | 0,51 | -5,34 | 0,00 | 0,00 |
| Arhgef9 (ENSMUSG00000025656) | 104,35 | -2,71 | 0,50 | -5,46 | 0,00 | 0,00 |
| Pcx (ENSMUSG00000024892) | 80,20 | -2,70 | 0,74 | -3,64 | 0,00 | 0,00 |
| Klhl3 (ENSMUSG00000014164) | 13,92 | -2,69 | 0,88 | -3,05 | 0,00 | 0,02 |
| Mastl (ENSMUSG00000026779) | 185,06 | -2,69 | 0,48 | -5,66 | 0,00 | 0,00 |
| Fbxo5 (ENSMUSG00000019773) | 424,65 | -2,68 | 0,53 | -5,03 | 0,00 | 0,00 |
| Cit (ENSMUSG00000029516) | 493,58 | -2,67 | 0,46 | -5,85 | 0,00 | 0,00 |
| Napsa (ENSMUSG00000002204) | 605,65 | -2,66 | 0,17 | -15,58 | 0,00 | 0,00 |
| Kif20a (ENSMUSG00000003779) | 1286,02 | -2,65 | 0,31 | -8,45 | 0,00 | 0,00 |
| Hist1h2bk (ENSMUSG00000062727) | 40,82 | -2,65 | 0,65 | -4,05 | 0,00 | 0,00 |
| Stmn1 (ENSMUSG00000028832) | 6196,88 | -2,64 | 0,32 | -8,16 | 0,00 | 0,00 |
| Neil3 (ENSMUSG00000039396) | 264,55 | -2,63 | 0,45 | -5,80 | 0,00 | 0,00 |
| Dlgap5 (ENSMUSG00000037544) | 760,01 | -2,63 | 0,33 | -7,89 | 0,00 | 0,00 |
| Gm44346 (ENSMUSG00000106281) | 56,87 | -2,61 | 0,51 | -5,11 | 0,00 | 0,00 |
| Aunip (ENSMUSG00000078521) | 68,44 | -2,61 | 0,94 | -2,77 | 0,01 | 0,04 |
| Ncapg (ENSMUSG00000015880) | 867,04 | -2,61 | 0,22 | -11,88 | 0,00 | 0,00 |
| Gm11223 (ENSMUSG00000046341) | 68,83 | -2,61 | 0,55 | -4,78 | 0,00 | 0,00 |
| Xlr (ENSMUSG00000054626) | 76,65 | -2,60 | 0,63 | -4,13 | 0,00 | 0,00 |
| Kif22 (ENSMUSG00000030677) | 1017,99 | -2,58 | 0,36 | -7,24 | 0,00 | 0,00 |
| Mtfr2 (ENSMUSG00000019992) | 158,29 | -2,58 | 0,46 | -5,63 | 0,00 | 0,00 |
| Cdca8 (ENSMUSG00000028873) | 1378,69 | -2,58 | 0,29 | -8,85 | 0,00 | 0,00 |
| Pcdhgc4 (ENSMUSG00000023036) | 159,07 | -2,58 | 0,72 | -3,58 | 0,00 | 0,01 |
| Gm11336 (ENSMUSG00000080999) | 31,30 | -2,57 | 0,54 | -4,74 | 0,00 | 0,00 |
| Lrr1 (ENSMUSG00000034883) | 103,00 | -2,55 | 0,45 | -5,60 | 0,00 | 0,00 |
| Cdca5 (ENSMUSG00000024791) | 721,99 | -2,54 | 0,40 | -6,43 | 0,00 | 0,00 |
| Bdh2 (ENSMUSG00000028167) | 45,99 | -2,54 | 0,54 | -4,75 | 0,00 | 0,00 |
| Cdca2 (ENSMUSG00000048922) | 695,64 | -2,54 | 0,26 | -9,83 | 0,00 | 0,00 |
| BC030867 (ENSMUSG00000034773) | 203,92 | -2,53 | 0,63 | -3,98 | 0,00 | 0,00 |
| Espl1 (ENSMUSG00000058290) | 673,60 | -2,51 | 0,42 | -5,92 | 0,00 | 0,00 |
| Slc17a6 (ENSMUSG00000030500) | 51,32 | -2,51 | 0,86 | -2,90 | 0,00 | 0,03 |
| Gm44359 (ENSMUSG00000105924) | 82,70 | -2,50 | 0,49 | -5,10 | 0,00 | 0,00 |
| Hist1h2ao (ENSMUSG00000094248) | 243,27 | -2,50 | 0,43 | -5,84 | 0,00 | 0,00 |
| Bub1b (ENSMUSG00000040084) | 1481,78 | -2,49 | 0,31 | -8,13 | 0,00 | 0,00 |
| Rad51ap1 (ENSMUSG00000030346) | 360,85 | -2,49 | 0,38 | -6,52 | 0,00 | 0,00 |
| Rad54b (ENSMUSG00000078773) | 181,97 | -2,47 | 0,50 | -4,99 | 0,00 | 0,00 |
| Ncapg2 (ENSMUSG00000042029) | 1136,77 | -2,47 | 0,35 | -7,01 | 0,00 | 0,00 |
| Knstrn (ENSMUSG00000027331) | 523,71 | -2,46 | 0,34 | -7,24 | 0,00 | 0,00 |
| Rad51 (ENSMUSG00000027323) | 1492,46 | -2,46 | 0,25 | -9,75 | 0,00 | 0,00 |
| Hist1h3i (ENSMUSG00000101972) | 77,45 | -2,46 | 0,36 | -6,84 | 0,00 | 0,00 |
| Spag5 (ENSMUSG00000002055) | 633,31 | -2,45 | 0,34 | -7,18 | 0,00 | 0,00 |
| Slc9b2 (ENSMUSG00000037994) | 353,86 | -2,45 | 0,39 | -6,29 | 0,00 | 0,00 |
| Gm20441 (ENSMUSG00000092360) | 10,85 | -2,44 | 0,80 | -3,04 | 0,00 | 0,02 |
| Kifc1 (ENSMUSG00000079553) | 127,08 | -2,43 | 0,80 | -3,05 | 0,00 | 0,02 |
| Serpinb1a (ENSMUSG00000044734) | 731,55 | -2,43 | 0,46 | -5,34 | 0,00 | 0,00 |
| Prc1 (ENSMUSG00000038943) | 948,37 | -2,41 | 0,58 | -4,13 | 0,00 | 0,00 |
| Eme1 (ENSMUSG00000039055) | 214,88 | -2,40 | 0,40 | -6,03 | 0,00 | 0,00 |
| Itih5 (ENSMUSG00000025780) | 1086,66 | -2,38 | 0,36 | -6,66 | 0,00 | 0,00 |
| Plpp1 (ENSMUSG00000021759) | 11,98 | -2,38 | 0,87 | -2,72 | 0,01 | 0,05 |
| Tk1 (ENSMUSG00000025574) | 1208,57 | -2,38 | 0,36 | -6,69 | 0,00 | 0,00 |
| Chit1 (ENSMUSG00000026450) | 84,48 | -2,38 | 0,68 | -3,48 | 0,00 | 0,01 |
| Cenpm (ENSMUSG00000068101) | 384,24 | -2,37 | 0,41 | -5,75 | 0,00 | 0,00 |
| Slc22a15 (ENSMUSG00000033147) | 148,63 | -2,37 | 0,63 | -3,75 | 0,00 | 0,00 |
| Gldc (ENSMUSG00000024827) | 70,30 | -2,36 | 0,77 | -3,06 | 0,00 | 0,02 |
| 2410004P03Rik (ENSMUSG00000071: | 117,49 | -2,36 | 0,49 | -4,85 | 0,00 | 0,00 |
| Brip1 (ENSMUSG00000034329) | 624,91 | -2,36 | 0,34 | -6,88 | 0,00 | 0,00 |
| Gen1 (ENSMUSG00000051235) | 160,48 | -2,36 | 0,78 | -3,04 | 0,00 | 0,02 |

| | | | | | | |
|---|---|---|---|---|---|---|
| Gm4890 (ENSMUSG00000097174) | 24,55 | -2,35 | 0,71 | -3,32 | 0,00 | 0,01 |
| Mcm10 (ENSMUSG00000026669) | 624,67 | -2,34 | 0,73 | -3,21 | 0,00 | 0,01 |
| Xkr5 (ENSMUSG00000039814) | 40,93 | -2,34 | 0,72 | -3,27 | 0,00 | 0,01 |
| Cenpk (ENSMUSG00000021714) | 350,98 | -2,34 | 0,42 | -5,59 | 0,00 | 0,00 |
| Rad54l (ENSMUSG00000028702) | 875,26 | -2,33 | 0,31 | -7,43 | 0,00 | 0,00 |
| Jazf1 (ENSMUSG00000063568) | 95,04 | -2,32 | 0,58 | -4,01 | 0,00 | 0,00 |
| D430020J02Rik (ENSMUSG00000112‹ | 130,85 | -2,31 | 0,45 | -5,11 | 0,00 | 0,00 |
| Stxbp1 (ENSMUSG00000026797) | 42,01 | -2,31 | 0,82 | -2,80 | 0,01 | 0,04 |
| Bambi-ps1 (ENSMUSG00000081219) | 32,46 | -2,31 | 0,80 | -2,89 | 0,00 | 0,03 |
| Dsel (ENSMUSG00000038702) | 13,79 | -2,29 | 0,71 | -3,24 | 0,00 | 0,01 |
| Rrm2 (ENSMUSG00000020649) | 3892,54 | -2,29 | 0,24 | -9,75 | 0,00 | 0,00 |
| Cenpp (ENSMUSG00000021391) | 182,29 | -2,29 | 0,69 | -3,32 | 0,00 | 0,01 |
| Cenpn (ENSMUSG00000031756) | 230,16 | -2,28 | 0,36 | -6,27 | 0,00 | 0,00 |
| Ltbp3 (ENSMUSG00000024940) | 98,73 | -2,28 | 0,74 | -3,09 | 0,00 | 0,02 |
| Spc25 (ENSMUSG00000005233) | 500,89 | -2,26 | 0,29 | -7,89 | 0,00 | 0,00 |
| Ticrr (ENSMUSG00000046591) | 584,44 | -2,25 | 0,37 | -6,06 | 0,00 | 0,00 |
| Kif18b (ENSMUSG00000051378) | 680,90 | -2,25 | 0,51 | -4,44 | 0,00 | 0,00 |
| Rgs8 (ENSMUSG00000042671) | 201,93 | -2,24 | 0,73 | -3,06 | 0,00 | 0,02 |
| Tacc3 (ENSMUSG00000037313) | 1965,04 | -2,24 | 0,20 | -10,96 | 0,00 | 0,00 |
| Klra3 (ENSMUSG00000067591) | 96,41 | -2,24 | 0,51 | -4,43 | 0,00 | 0,00 |
| Aebp1 (ENSMUSG00000020473) | 50,11 | -2,24 | 0,70 | -3,18 | 0,00 | 0,02 |
| Kif20b (ENSMUSG00000024795) | 371,48 | -2,24 | 0,25 | -8,83 | 0,00 | 0,00 |
| Asf1b (ENSMUSG00000005470) | 1682,60 | -2,23 | 0,21 | -10,49 | 0,00 | 0,00 |
| Spdl1 (ENSMUSG00000069910) | 218,43 | -2,23 | 0,38 | -5,82 | 0,00 | 0,00 |
| Sgo2a (ENSMUSG00000026039) | 188,32 | -2,22 | 0,51 | -4,40 | 0,00 | 0,00 |
| Il1r2 (ENSMUSG00000026073) | 302,07 | -2,21 | 0,57 | -3,85 | 0,00 | 0,00 |
| Hist1h2aj (ENSMUSG00000080076) | 54,23 | -2,20 | 0,45 | -4,89 | 0,00 | 0,00 |
| Hhat (ENSMUSG00000037375) | 98,65 | -2,20 | 0,74 | -2,98 | 0,00 | 0,03 |
| Ptpn5 (ENSMUSG00000030854) | 98,35 | -2,19 | 0,73 | -3,00 | 0,00 | 0,03 |
| Hist1h2ag (ENSMUSG00000069301) | 143,92 | -2,17 | 0,61 | -3,57 | 0,00 | 0,01 |
| Cdc6 (ENSMUSG00000017499) | 889,57 | -2,17 | 0,28 | -7,67 | 0,00 | 0,00 |
| Msh5 (ENSMUSG00000007035) | 37,72 | -2,17 | 0,80 | -2,72 | 0,01 | 0,05 |
| Ect2 (ENSMUSG00000027699) | 439,83 | -2,16 | 0,40 | -5,41 | 0,00 | 0,00 |
| Hist1h4d (ENSMUSG00000061482) | 887,81 | -2,16 | 0,35 | -6,24 | 0,00 | 0,00 |
| Rab4a (ENSMUSG00000019478) | 80,69 | -2,15 | 0,45 | -4,81 | 0,00 | 0,00 |
| Synpo (ENSMUSG00000043079) | 1177,24 | -2,15 | 0,31 | -6,84 | 0,00 | 0,00 |
| Ccnf (ENSMUSG00000072082) | 535,31 | -2,14 | 0,39 | -5,53 | 0,00 | 0,00 |
| Fcer1g (ENSMUSG00000058715) | 189,86 | -2,14 | 0,42 | -5,16 | 0,00 | 0,00 |
| Wdr62 (ENSMUSG00000037020) | 307,21 | -2,13 | 0,22 | -9,74 | 0,00 | 0,00 |
| Lat2 (ENSMUSG00000040751) | 572,03 | -2,10 | 0,28 | -7,56 | 0,00 | 0,00 |
| Top2a (ENSMUSG00000020914) | 4637,31 | -2,10 | 0,25 | -8,43 | 0,00 | 0,00 |
| Fam229b (ENSMUSG00000051736) | 26,06 | -2,09 | 0,75 | -2,79 | 0,01 | 0,04 |
| Cdkn2c (ENSMUSG00000028551) | 421,19 | -2,09 | 0,35 | -6,01 | 0,00 | 0,00 |
| Csrp2 (ENSMUSG00000020186) | 48,80 | -2,08 | 0,64 | -3,25 | 0,00 | 0,01 |
| Gins1 (ENSMUSG00000027454) | 250,29 | -2,08 | 0,44 | -4,72 | 0,00 | 0,00 |
| Traip (ENSMUSG00000032586) | 326,67 | -2,07 | 0,43 | -4,85 | 0,00 | 0,00 |
| Dhrs13os (ENSMUSG00000087050) | 13,93 | -2,07 | 0,52 | -3,95 | 0,00 | 0,00 |
| Cxcl16 (ENSMUSG00000018920) | 42,71 | -2,07 | 0,44 | -4,72 | 0,00 | 0,00 |
| Ska3 (ENSMUSG00000021965) | 217,27 | -2,06 | 0,36 | -5,70 | 0,00 | 0,00 |
| Fam129c (ENSMUSG00000043243) | 79,12 | -2,05 | 0,66 | -3,10 | 0,00 | 0,02 |
| Lockd (ENSMUSG00000098318) | 361,32 | -2,05 | 0,34 | -5,99 | 0,00 | 0,00 |
| Oip5 (ENSMUSG00000072980) | 195,82 | -2,05 | 0,55 | -3,71 | 0,00 | 0,00 |
| Cks1b (ENSMUSG00000028044) | 1534,65 | -2,04 | 0,25 | -8,16 | 0,00 | 0,00 |
| Pvrig (ENSMUSG00000109713) | 75,64 | -2,04 | 0,54 | -3,75 | 0,00 | 0,00 |
| Gm4668 (ENSMUSG00000111243) | 16,17 | -2,04 | 0,72 | -2,85 | 0,00 | 0,04 |
| Mad2l1 (ENSMUSG00000029910) | 1328,13 | -2,03 | 0,27 | -7,46 | 0,00 | 0,00 |
| Nrn1 (ENSMUSG00000039114) | 802,17 | -2,03 | 0,14 | -14,56 | 0,00 | 0,00 |

| | | | | | | |
|---|---|---|---|---|---|---|
| Ncaph (ENSMUSG00000034906) | 1427,51 | -2,02 | 0,30 | -6,78 | 0,00 | 0,00 |
| Cip2a (ENSMUSG00000033031) | 245,63 | -2,01 | 0,32 | -6,35 | 0,00 | 0,00 |
| Poc1a (ENSMUSG00000023345) | 377,13 | -2,00 | 0,21 | -9,57 | 0,00 | 0,00 |
| Ndc80 (ENSMUSG00000024056) | 807,18 | -1,99 | 0,23 | -8,63 | 0,00 | 0,00 |
| E2f2 (ENSMUSG00000018983) | 609,90 | -1,99 | 0,27 | -7,49 | 0,00 | 0,00 |
| Cenph (ENSMUSG00000045273) | 278,01 | -1,99 | 0,26 | -7,61 | 0,00 | 0,00 |
| Tcf19 (ENSMUSG00000050410) | 1259,98 | -1,99 | 0,27 | -7,44 | 0,00 | 0,00 |
| Arhgap19 (ENSMUSG00000025154) | 219,28 | -1,98 | 0,72 | -2,74 | 0,01 | 0,05 |
| E2f1 (ENSMUSG00000027490) | 940,18 | -1,98 | 0,22 | -9,19 | 0,00 | 0,00 |
| Kif23 (ENSMUSG00000032254) | 844,98 | -1,97 | 0,44 | -4,45 | 0,00 | 0,00 |
| Chaf1a (ENSMUSG00000002835) | 1725,40 | -1,96 | 0,34 | -5,78 | 0,00 | 0,00 |
| 1500009L16Rik (ENSMUSG0000008[illegible] | 167,00 | -1,96 | 0,36 | -5,39 | 0,00 | 0,00 |
| Arhgap11a (ENSMUSG00000041219) | 649,37 | -1,96 | 0,31 | -6,35 | 0,00 | 0,00 |
| Orc1 (ENSMUSG00000028587) | 331,72 | -1,95 | 0,47 | -4,10 | 0,00 | 0,00 |
| Eno3 (ENSMUSG00000060600) | 1733,29 | -1,95 | 0,34 | -5,78 | 0,00 | 0,00 |
| Tmem106a (ENSMUSG00000034947) | 74,78 | -1,94 | 0,52 | -3,75 | 0,00 | 0,00 |
| Gm42921 (ENSMUSG00000106441) | 65,04 | -1,93 | 0,53 | -3,63 | 0,00 | 0,00 |
| Tns1 (ENSMUSG00000055322) | 418,47 | -1,93 | 0,31 | -6,17 | 0,00 | 0,00 |
| Cenps (ENSMUSG00000073705) | 208,54 | -1,91 | 0,28 | -6,93 | 0,00 | 0,00 |
| Hspb1 (ENSMUSG00000004951) | 46,42 | -1,91 | 0,39 | -4,92 | 0,00 | 0,00 |
| Izumo1r (ENSMUSG00000031933) | 757,92 | -1,91 | 0,31 | -6,17 | 0,00 | 0,00 |
| Cryzl2 (ENSMUSG00000033488) | 51,31 | -1,91 | 0,60 | -3,17 | 0,00 | 0,02 |
| Gm26765 (ENSMUSG00000097664) | 21,44 | -1,91 | 0,70 | -2,73 | 0,01 | 0,05 |
| Tyms (ENSMUSG00000025747) | 1393,98 | -1,90 | 0,28 | -6,86 | 0,00 | 0,00 |
| Tmem171 (ENSMUSG00000052485) | 174,78 | -1,90 | 0,28 | -6,71 | 0,00 | 0,00 |
| Ptgr1 (ENSMUSG00000028378) | 84,65 | -1,90 | 0,41 | -4,64 | 0,00 | 0,00 |
| Prss2 (ENSMUSG00000057163) | 178,98 | -1,89 | 0,29 | -6,42 | 0,00 | 0,00 |
| Klri2 (ENSMUSG00000043932) | 338,84 | -1,88 | 0,32 | -5,85 | 0,00 | 0,00 |
| Ramp1 (ENSMUSG00000034353) | 109,76 | -1,87 | 0,36 | -5,26 | 0,00 | 0,00 |
| Cdc45 (ENSMUSG00000000028) | 1125,72 | -1,87 | 0,24 | -7,78 | 0,00 | 0,00 |
| Cntln (ENSMUSG00000038070) | 96,78 | -1,87 | 0,46 | -4,09 | 0,00 | 0,00 |
| Tox (ENSMUSG00000041272) | 1365,36 | -1,87 | 0,24 | -7,94 | 0,00 | 0,00 |
| Gm4739 (ENSMUSG00000112808) | 3907,19 | -1,86 | 0,22 | -8,30 | 0,00 | 0,00 |
| Dscc1 (ENSMUSG00000022422) | 223,47 | -1,86 | 0,31 | -5,99 | 0,00 | 0,00 |
| C1qtnf6 (ENSMUSG00000022440) | 212,74 | -1,86 | 0,32 | -5,74 | 0,00 | 0,00 |
| Dtl (ENSMUSG00000037474) | 1681,93 | -1,86 | 0,26 | -7,22 | 0,00 | 0,00 |
| Lig1 (ENSMUSG00000056394) | 2366,74 | -1,85 | 0,26 | -7,03 | 0,00 | 0,00 |
| Brca2 (ENSMUSG00000041147) | 286,00 | -1,85 | 0,30 | -6,18 | 0,00 | 0,00 |
| Rsad2 (ENSMUSG00000020641) | 203,17 | -1,85 | 0,40 | -4,64 | 0,00 | 0,00 |
| Ckap2 (ENSMUSG00000037725) | 405,59 | -1,85 | 0,28 | -6,64 | 0,00 | 0,00 |
| Fam83d (ENSMUSG00000027654) | 195,82 | -1,85 | 0,31 | -5,89 | 0,00 | 0,00 |
| Incenp (ENSMUSG00000024660) | 1843,74 | -1,83 | 0,21 | -8,51 | 0,00 | 0,00 |
| Gpx7 (ENSMUSG00000028597) | 110,58 | -1,83 | 0,54 | -3,38 | 0,00 | 0,01 |
| Fanca (ENSMUSG00000032815) | 280,00 | -1,82 | 0,31 | -5,81 | 0,00 | 0,00 |
| Lyn (ENSMUSG00000042228) | 191,03 | -1,81 | 0,26 | -6,99 | 0,00 | 0,00 |
| Zwilch (ENSMUSG00000032400) | 191,02 | -1,80 | 0,45 | -4,00 | 0,00 | 0,00 |
| Pter (ENSMUSG00000026730) | 72,19 | -1,78 | 0,45 | -3,97 | 0,00 | 0,00 |
| Mcm8 (ENSMUSG00000027353) | 270,92 | -1,78 | 0,33 | -5,47 | 0,00 | 0,00 |
| Hist1h4b (ENSMUSG00000069266) | 20,46 | -1,77 | 0,47 | -3,74 | 0,00 | 0,00 |
| Nmral1 (ENSMUSG00000063445) | 394,98 | -1,77 | 0,35 | -5,13 | 0,00 | 0,00 |
| Trip13 (ENSMUSG00000021569) | 539,50 | -1,77 | 0,41 | -4,32 | 0,00 | 0,00 |
| Dclk2 (ENSMUSG00000028078) | 119,88 | -1,76 | 0,49 | -3,61 | 0,00 | 0,00 |
| Hist1h2ai (ENSMUSG00000071516) | 50,42 | -1,76 | 0,63 | -2,80 | 0,01 | 0,04 |
| Gstt1 (ENSMUSG00000001663) | 105,91 | -1,76 | 0,41 | -4,27 | 0,00 | 0,00 |
| 1700097N02Rik (ENSMUSG0000009[illegible] | 297,06 | -1,75 | 0,32 | -5,55 | 0,00 | 0,00 |
| Hpse (ENSMUSG00000035273) | 231,48 | -1,75 | 0,29 | -6,04 | 0,00 | 0,00 |
| Brca1 (ENSMUSG00000017146) | 379,99 | -1,75 | 0,42 | -4,14 | 0,00 | 0,00 |

| | | | | | | |
|---|---|---|---|---|---|---|
| Mnd1 (ENSMUSG00000033752) | 35,38 | -1,75 | 0,39 | -4,54 | 0,00 | 0,00 |
| Ly6a2 (ENSMUSG00000102051) | 424,05 | -1,75 | 0,20 | -8,84 | 0,00 | 0,00 |
| Hmgb2 (ENSMUSG00000054717) | 7545,81 | -1,74 | 0,22 | -7,78 | 0,00 | 0,00 |
| Fignl1 (ENSMUSG00000035455) | 1026,42 | -1,74 | 0,15 | -11,64 | 0,00 | 0,00 |
| Nsl1 (ENSMUSG00000062510) | 187,91 | -1,72 | 0,40 | -4,35 | 0,00 | 0,00 |
| 2700099C18Rik (ENSMUSG00000098( | 62,74 | -1,72 | 0,48 | -3,62 | 0,00 | 0,00 |
| Xrcc3 (ENSMUSG00000021287) | 148,40 | -1,72 | 0,53 | -3,22 | 0,00 | 0,01 |
| Hist2h2ac (ENSMUSG00000068855) | 41,79 | -1,72 | 0,49 | -3,47 | 0,00 | 0,01 |
| Csf2ra (ENSMUSG00000059326) | 55,04 | -1,71 | 0,48 | -3,55 | 0,00 | 0,01 |
| Serpinb1b (ENSMUSG00000051029) | 74,49 | -1,71 | 0,52 | -3,28 | 0,00 | 0,01 |
| Dapl1 (ENSMUSG00000026989) | 701,04 | -1,71 | 0,30 | -5,62 | 0,00 | 0,00 |
| Dhfr (ENSMUSG00000021707) | 887,26 | -1,70 | 0,28 | -6,11 | 0,00 | 0,00 |
| Klre1 (ENSMUSG00000050241) | 870,71 | -1,67 | 0,21 | -8,00 | 0,00 | 0,00 |
| Hist2h3b (ENSMUSG00000074403) | 28,73 | -1,67 | 0,58 | -2,86 | 0,00 | 0,04 |
| Gm42674 (ENSMUSG00000105518) | 37,56 | -1,66 | 0,54 | -3,07 | 0,00 | 0,02 |
| Pask (ENSMUSG00000026274) | 413,99 | -1,66 | 0,35 | -4,77 | 0,00 | 0,00 |
| Ncapd2 (ENSMUSG00000038252) | 2818,95 | -1,66 | 0,19 | -8,82 | 0,00 | 0,00 |
| Lair1 (ENSMUSG00000055541) | 225,96 | -1,66 | 0,54 | -3,08 | 0,00 | 0,02 |
| Cenpw (ENSMUSG00000075266) | 369,61 | -1,66 | 0,27 | -6,20 | 0,00 | 0,00 |
| Racgap1 (ENSMUSG00000023015) | 3258,62 | -1,65 | 0,18 | -9,07 | 0,00 | 0,00 |
| Vim (ENSMUSG00000026728) | 5752,84 | -1,65 | 0,29 | -5,73 | 0,00 | 0,00 |
| Gins2 (ENSMUSG00000031821) | 845,98 | -1,65 | 0,32 | -5,20 | 0,00 | 0,00 |
| Fut7 (ENSMUSG00000036587) | 260,49 | -1,64 | 0,36 | -4,53 | 0,00 | 0,00 |
| Uhrf1 (ENSMUSG00000001228) | 4992,69 | -1,63 | 0,20 | -8,25 | 0,00 | 0,00 |
| Als2cl (ENSMUSG00000044037) | 361,61 | -1,62 | 0,40 | -4,09 | 0,00 | 0,00 |
| Gstt3 (ENSMUSG00000001665) | 250,74 | -1,62 | 0,25 | -6,36 | 0,00 | 0,00 |
| Serpinb9b (ENSMUSG00000021403) | 838,43 | -1,61 | 0,49 | -3,28 | 0,00 | 0,01 |
| Ypel2 (ENSMUSG00000018427) | 36,25 | -1,61 | 0,54 | -2,99 | 0,00 | 0,03 |
| Aurka (ENSMUSG00000027496) | 442,69 | -1,61 | 0,27 | -5,99 | 0,00 | 0,00 |
| Gm43305 (ENSMUSG00000105703) | 4970,77 | -1,60 | 0,17 | -9,44 | 0,00 | 0,00 |
| Eng (ENSMUSG00000026814) | 124,57 | -1,60 | 0,58 | -2,74 | 0,01 | 0,05 |
| Ajuba (ENSMUSG00000022178) | 272,82 | -1,60 | 0,35 | -4,61 | 0,00 | 0,00 |
| Gm10282 (ENSMUSG00000070713) | 24,62 | -1,60 | 0,52 | -3,09 | 0,00 | 0,02 |
| Gpm6b (ENSMUSG00000031342) | 308,37 | -1,59 | 0,41 | -3,90 | 0,00 | 0,00 |
| Idi1 (ENSMUSG00000058258) | 754,81 | -1,56 | 0,30 | -5,19 | 0,00 | 0,00 |
| Gpr34 (ENSMUSG00000040229) | 97,26 | -1,55 | 0,38 | -4,08 | 0,00 | 0,00 |
| Smc2 (ENSMUSG00000028312) | 1723,86 | -1,55 | 0,23 | -6,68 | 0,00 | 0,00 |
| Tnnt1 (ENSMUSG00000064179) | 63,53 | -1,54 | 0,41 | -3,78 | 0,00 | 0,00 |
| Ccdc34 (ENSMUSG00000027160) | 186,41 | -1,53 | 0,24 | -6,41 | 0,00 | 0,00 |
| Mns1 (ENSMUSG00000032221) | 330,80 | -1,53 | 0,27 | -5,70 | 0,00 | 0,00 |
| Lclat1 (ENSMUSG00000054469) | 611,78 | -1,53 | 0,36 | -4,28 | 0,00 | 0,00 |
| Rfc4 (ENSMUSG00000022881) | 1362,36 | -1,53 | 0,19 | -7,89 | 0,00 | 0,00 |
| S100a4 (ENSMUSG00000001020) | 2527,41 | -1,53 | 0,26 | -5,77 | 0,00 | 0,00 |
| Slc12a8 (ENSMUSG00000035506) | 59,04 | -1,53 | 0,43 | -3,55 | 0,00 | 0,01 |
| Pole (ENSMUSG00000007080) | 1255,18 | -1,52 | 0,38 | -4,01 | 0,00 | 0,00 |
| Med12l (ENSMUSG00000056476) | 48,04 | -1,52 | 0,50 | -3,03 | 0,00 | 0,02 |
| Gm11914 (ENSMUSG00000082674) | 47,30 | -1,51 | 0,39 | -3,89 | 0,00 | 0,00 |
| Ppic (ENSMUSG00000024538) | 295,77 | -1,51 | 0,23 | -6,49 | 0,00 | 0,00 |
| Gm48274 (ENSMUSG00000111083) | 43,16 | -1,50 | 0,44 | -3,40 | 0,00 | 0,01 |
| Slc35f5 (ENSMUSG00000026342) | 88,73 | -1,50 | 0,55 | -2,74 | 0,01 | 0,05 |
| Mcm5 (ENSMUSG00000005410) | 9562,13 | -1,50 | 0,23 | -6,48 | 0,00 | 0,00 |
| Lmnb1 (ENSMUSG00000024590) | 2724,73 | -1,50 | 0,27 | -5,60 | 0,00 | 0,00 |
| Arhgap21 (ENSMUSG00000036591) | 432,31 | -1,50 | 0,51 | -2,93 | 0,00 | 0,03 |
| Plk4 (ENSMUSG00000025758) | 704,65 | -1,50 | 0,28 | -5,40 | 0,00 | 0,00 |
| Gorasp1 (ENSMUSG00000032513) | 110,55 | -1,49 | 0,51 | -2,95 | 0,00 | 0,03 |
| Chtf18 (ENSMUSG00000019214) | 683,29 | -1,49 | 0,20 | -7,42 | 0,00 | 0,00 |
| Slc25a13 (ENSMUSG00000015112) | 439,65 | -1,49 | 0,40 | -3,76 | 0,00 | 0,00 |

| | | | | | | |
|---|---|---|---|---|---|---|
| Tbc1d7 (ENSMUSG00000021368) | 323,17 | -1,49 | 0,32 | -4,67 | 0,00 | 0,00 |
| Rfc3 (ENSMUSG00000033970) | 751,88 | -1,48 | 0,24 | -6,15 | 0,00 | 0,00 |
| Gm16576 (ENSMUSG00000087543) | 64,50 | -1,48 | 0,54 | -2,75 | 0,01 | 0,05 |
| Rmi2 (ENSMUSG00000037991) | 432,91 | -1,47 | 0,30 | -4,98 | 0,00 | 0,00 |
| Tspyl4 (ENSMUSG00000039485) | 208,54 | -1,47 | 0,41 | -3,58 | 0,00 | 0,00 |
| Fam221a (ENSMUSG00000047115) | 35,11 | -1,47 | 0,53 | -2,77 | 0,01 | 0,04 |
| Cenpa (ENSMUSG00000029177) | 3318,58 | -1,47 | 0,20 | -7,43 | 0,00 | 0,00 |
| Timp2 (ENSMUSG00000017466) | 207,76 | -1,47 | 0,44 | -3,31 | 0,00 | 0,01 |
| Atad5 (ENSMUSG00000017550) | 299,98 | -1,47 | 0,44 | -3,36 | 0,00 | 0,01 |
| Kif18a (ENSMUSG00000027115) | 225,51 | -1,46 | 0,21 | -6,80 | 0,00 | 0,00 |
| A430088P11Rik (ENSMUSG0000011[illegible] | 101,66 | -1,45 | 0,35 | -4,12 | 0,00 | 0,00 |
| Dck (ENSMUSG00000029366) | 1422,21 | -1,45 | 0,19 | -7,73 | 0,00 | 0,00 |
| Gmnn (ENSMUSG00000006715) | 837,04 | -1,45 | 0,25 | -5,83 | 0,00 | 0,00 |
| Mrnip (ENSMUSG00000020381) | 69,59 | -1,44 | 0,39 | -3,74 | 0,00 | 0,00 |
| Hells (ENSMUSG00000025001) | 941,79 | -1,44 | 0,29 | -4,97 | 0,00 | 0,00 |
| Blm (ENSMUSG00000030528) | 308,88 | -1,44 | 0,20 | -7,35 | 0,00 | 0,00 |
| Nebl (ENSMUSG00000053702) | 43,46 | -1,43 | 0,38 | -3,78 | 0,00 | 0,00 |
| Ccne2 (ENSMUSG00000028212) | 266,57 | -1,42 | 0,38 | -3,74 | 0,00 | 0,00 |
| Selenoh (ENSMUSG00000076437) | 1626,98 | -1,42 | 0,19 | -7,50 | 0,00 | 0,00 |
| Wls (ENSMUSG00000028173) | 1000,56 | -1,42 | 0,20 | -7,17 | 0,00 | 0,00 |
| 2810408I11Rik (ENSMUSG000000872 | 54,81 | -1,42 | 0,33 | -4,26 | 0,00 | 0,00 |
| Mcm7 (ENSMUSG00000029730) | 7441,89 | -1,42 | 0,24 | -5,80 | 0,00 | 0,00 |
| Cuedc1 (ENSMUSG00000018378) | 62,86 | -1,41 | 0,50 | -2,83 | 0,00 | 0,04 |
| Gm10182 (ENSMUSG00000094627) | 622,14 | -1,41 | 0,24 | -5,91 | 0,00 | 0,00 |
| H2-DMb1 (ENSMUSG00000079547) | 50,23 | -1,41 | 0,43 | -3,28 | 0,00 | 0,01 |
| Gm4316 (ENSMUSG00000110537) | 2276,80 | -1,40 | 0,16 | -8,94 | 0,00 | 0,00 |
| Hmgn2 (ENSMUSG00000003038) | 8162,51 | -1,40 | 0,21 | -6,63 | 0,00 | 0,00 |
| Ccdc28b (ENSMUSG00000028795) | 104,45 | -1,40 | 0,20 | -6,84 | 0,00 | 0,00 |
| Ncf2 (ENSMUSG00000026480) | 88,13 | -1,40 | 0,47 | -2,95 | 0,00 | 0,03 |
| Tmlhe (ENSMUSG00000079834) | 140,91 | -1,40 | 0,39 | -3,61 | 0,00 | 0,00 |
| Cmpk2 (ENSMUSG00000020638) | 540,98 | -1,40 | 0,30 | -4,65 | 0,00 | 0,00 |
| Rrm1 (ENSMUSG00000030978) | 7395,22 | -1,39 | 0,22 | -6,43 | 0,00 | 0,00 |
| Gm47761 (ENSMUSG00000112478) | 51,42 | -1,38 | 0,41 | -3,36 | 0,00 | 0,01 |
| Gm35279 (ENSMUSG00000114203) | 58,05 | -1,38 | 0,31 | -4,53 | 0,00 | 0,00 |
| Ccdc163 (ENSMUSG00000028689) | 107,78 | -1,38 | 0,38 | -3,61 | 0,00 | 0,00 |
| Iigp1 (ENSMUSG00000054072) | 533,04 | -1,38 | 0,20 | -6,71 | 0,00 | 0,00 |
| Ska2 (ENSMUSG00000020492) | 266,12 | -1,38 | 0,27 | -5,17 | 0,00 | 0,00 |
| BC055324 (ENSMUSG00000041406) | 240,84 | -1,37 | 0,33 | -4,11 | 0,00 | 0,00 |
| Lancl2 (ENSMUSG00000062190) | 226,63 | -1,37 | 0,23 | -6,01 | 0,00 | 0,00 |
| Bckdhb (ENSMUSG00000032263) | 135,42 | -1,37 | 0,27 | -5,05 | 0,00 | 0,00 |
| Rnf227 (ENSMUSG00000043419) | 99,26 | -1,36 | 0,38 | -3,54 | 0,00 | 0,01 |
| Ifi27l2a (ENSMUSG00000079017) | 5303,54 | -1,35 | 0,19 | -7,20 | 0,00 | 0,00 |
| Dut (ENSMUSG00000027203) | 2359,50 | -1,35 | 0,22 | -6,06 | 0,00 | 0,00 |
| Pbx4 (ENSMUSG00000031860) | 58,71 | -1,35 | 0,44 | -3,04 | 0,00 | 0,02 |
| Plekha8 (ENSMUSG00000005225) | 134,21 | -1,34 | 0,41 | -3,30 | 0,00 | 0,01 |
| Slc25a42 (ENSMUSG00000002346) | 127,81 | -1,34 | 0,37 | -3,62 | 0,00 | 0,00 |
| Gm26694 (ENSMUSG00000097518) | 56,75 | -1,34 | 0,37 | -3,63 | 0,00 | 0,00 |
| Zfp862-ps (ENSMUSG00000107476) | 407,88 | -1,34 | 0,29 | -4,63 | 0,00 | 0,00 |
| Mcm3 (ENSMUSG00000041859) | 9191,86 | -1,33 | 0,20 | -6,78 | 0,00 | 0,00 |
| Mmp16 (ENSMUSG00000028226) | 233,93 | -1,33 | 0,36 | -3,70 | 0,00 | 0,00 |
| Prim2 (ENSMUSG00000026134) | 882,82 | -1,33 | 0,23 | -5,69 | 0,00 | 0,00 |
| Gm37416 (ENSMUSG00000103865) | 333,23 | -1,32 | 0,23 | -5,70 | 0,00 | 0,00 |
| Mms22l (ENSMUSG00000045751) | 746,45 | -1,32 | 0,29 | -4,53 | 0,00 | 0,00 |
| Spin4 (ENSMUSG00000071722) | 135,88 | -1,32 | 0,29 | -4,61 | 0,00 | 0,00 |
| Gm14276 (ENSMUSG00000082163) | 61,90 | -1,32 | 0,39 | -3,37 | 0,00 | 0,01 |
| Cacna1s (ENSMUSG00000026407) | 137,02 | -1,31 | 0,42 | -3,13 | 0,00 | 0,02 |
| Rnf135 (ENSMUSG00000020707) | 201,48 | -1,31 | 0,37 | -3,54 | 0,00 | 0,01 |

| | | | | | | |
|---|---|---|---|---|---|---|
| Pole2 (ENSMUSG00000020974) | 326,72 | -1,31 | 0,25 | -5,17 | 0,00 | 0,00 |
| Tlcd2 (ENSMUSG00000038217) | 164,62 | -1,31 | 0,31 | -4,27 | 0,00 | 0,00 |
| Osgepl1 (ENSMUSG00000026096) | 210,25 | -1,30 | 0,31 | -4,15 | 0,00 | 0,00 |
| Dnph1 (ENSMUSG00000040658) | 185,88 | -1,30 | 0,32 | -4,03 | 0,00 | 0,00 |
| Pxmp2 (ENSMUSG00000029499) | 178,77 | -1,30 | 0,26 | -4,96 | 0,00 | 0,00 |
| Timeless (ENSMUSG00000039994) | 1241,06 | -1,30 | 0,11 | -11,49 | 0,00 | 0,00 |
| Emid1 (ENSMUSG00000034164) | 113,49 | -1,30 | 0,27 | -4,87 | 0,00 | 0,00 |
| Endod1 (ENSMUSG00000037419) | 2735,34 | -1,29 | 0,21 | -6,12 | 0,00 | 0,00 |
| Gm17767 (ENSMUSG00000099413) | 190,26 | -1,29 | 0,33 | -3,90 | 0,00 | 0,00 |
| Tspan2 (ENSMUSG00000027858) | 270,01 | -1,28 | 0,27 | -4,81 | 0,00 | 0,00 |
| Slc43a3 (ENSMUSG00000027074) | 800,24 | -1,28 | 0,29 | -4,36 | 0,00 | 0,00 |
| 2810455O05Rik (ENSMUSG0000008[illegible] | 33,10 | -1,28 | 0,38 | -3,34 | 0,00 | 0,01 |
| Rdm1 (ENSMUSG00000010362) | 721,74 | -1,26 | 0,23 | -5,59 | 0,00 | 0,00 |
| Lca5 (ENSMUSG00000032258) | 53,92 | -1,26 | 0,39 | -3,19 | 0,00 | 0,02 |
| Fah (ENSMUSG00000030630) | 120,61 | -1,26 | 0,43 | -2,90 | 0,00 | 0,03 |
| Hist1h1e (ENSMUSG00000051627) | 420,87 | -1,26 | 0,20 | -6,26 | 0,00 | 0,00 |
| Creb3l1 (ENSMUSG00000027230) | 85,61 | -1,26 | 0,39 | -3,23 | 0,00 | 0,01 |
| Plxdc2 (ENSMUSG00000026748) | 498,75 | -1,26 | 0,33 | -3,77 | 0,00 | 0,00 |
| Wdhd1 (ENSMUSG00000037572) | 906,41 | -1,25 | 0,36 | -3,50 | 0,00 | 0,01 |
| Hist1h4i (ENSMUSG00000060639) | 100,34 | -1,25 | 0,37 | -3,37 | 0,00 | 0,01 |
| Fam111a (ENSMUSG00000024691) | 1545,47 | -1,25 | 0,20 | -6,15 | 0,00 | 0,00 |
| Xdh (ENSMUSG00000024066) | 9180,95 | -1,25 | 0,15 | -8,53 | 0,00 | 0,00 |
| Cd63 (ENSMUSG00000025351) | 80,87 | -1,23 | 0,39 | -3,16 | 0,00 | 0,02 |
| Raph1 (ENSMUSG00000026014) | 254,25 | -1,23 | 0,32 | -3,80 | 0,00 | 0,00 |
| Gm10184 (ENSMUSG00000066878) | 242,02 | -1,23 | 0,28 | -4,42 | 0,00 | 0,00 |
| Ccp110 (ENSMUSG00000033904) | 305,24 | -1,23 | 0,44 | -2,79 | 0,01 | 0,04 |
| Pi4k2b (ENSMUSG00000029186) | 314,10 | -1,23 | 0,26 | -4,73 | 0,00 | 0,00 |
| Enkd1 (ENSMUSG00000013155) | 171,15 | -1,22 | 0,25 | -4,82 | 0,00 | 0,00 |
| Stard4 (ENSMUSG00000024378) | 667,37 | -1,22 | 0,24 | -5,05 | 0,00 | 0,00 |
| Cyp51 (ENSMUSG00000001467) | 1210,53 | -1,21 | 0,26 | -4,59 | 0,00 | 0,00 |
| Tjp2 (ENSMUSG00000024812) | 351,28 | -1,21 | 0,30 | -4,06 | 0,00 | 0,00 |
| Mcm4 (ENSMUSG00000022673) | 7958,76 | -1,21 | 0,14 | -8,92 | 0,00 | 0,00 |
| Haus4 (ENSMUSG00000022177) | 1532,12 | -1,20 | 0,18 | -6,53 | 0,00 | 0,00 |
| Ube2t (ENSMUSG00000026429) | 231,87 | -1,20 | 0,29 | -4,16 | 0,00 | 0,00 |
| Rfc5 (ENSMUSG00000029363) | 2749,82 | -1,20 | 0,25 | -4,89 | 0,00 | 0,00 |
| Sass6 (ENSMUSG00000027959) | 273,92 | -1,20 | 0,32 | -3,77 | 0,00 | 0,00 |
| Lgals1 (ENSMUSG00000068220) | 15248,09 | -1,20 | 0,17 | -7,17 | 0,00 | 0,00 |
| Gpr160 (ENSMUSG00000037661) | 584,22 | -1,20 | 0,17 | -7,23 | 0,00 | 0,00 |
| Nrm (ENSMUSG00000059791) | 1105,05 | -1,19 | 0,27 | -4,40 | 0,00 | 0,00 |
| Mzt2 (ENSMUSG00000022671) | 219,80 | -1,19 | 0,25 | -4,74 | 0,00 | 0,00 |
| Orc6 (ENSMUSG00000031697) | 814,94 | -1,19 | 0,19 | -6,42 | 0,00 | 0,00 |
| Mcm2 (ENSMUSG00000002870) | 8574,02 | -1,18 | 0,14 | -8,67 | 0,00 | 0,00 |
| Haspin (ENSMUSG00000050107) | 317,94 | -1,18 | 0,26 | -4,60 | 0,00 | 0,00 |
| Gm42047 (ENSMUSG00000110631) | 518,51 | -1,18 | 0,23 | -5,06 | 0,00 | 0,00 |
| Gm8203 (ENSMUSG00000101878) | 1509,98 | -1,18 | 0,20 | -5,77 | 0,00 | 0,00 |
| Spata24 (ENSMUSG00000024352) | 99,09 | -1,17 | 0,35 | -3,32 | 0,00 | 0,01 |
| Nt5dc2 (ENSMUSG00000071547) | 550,58 | -1,17 | 0,22 | -5,45 | 0,00 | 0,00 |
| Prdx4 (ENSMUSG00000025289) | 270,70 | -1,17 | 0,33 | -3,55 | 0,00 | 0,01 |
| Cmtm7 (ENSMUSG00000032436) | 5576,83 | -1,17 | 0,17 | -7,00 | 0,00 | 0,00 |
| Tipin (ENSMUSG00000032397) | 2084,77 | -1,17 | 0,17 | -7,05 | 0,00 | 0,00 |
| Pola1 (ENSMUSG00000006678) | 1378,16 | -1,16 | 0,30 | -3,92 | 0,00 | 0,00 |
| Cdc7 (ENSMUSG00000029283) | 494,39 | -1,15 | 0,17 | -6,96 | 0,00 | 0,00 |
| Naaa (ENSMUSG00000029413) | 153,07 | -1,15 | 0,34 | -3,39 | 0,00 | 0,01 |
| Gm45315 (ENSMUSG00000109925) | 250,86 | -1,15 | 0,18 | -6,54 | 0,00 | 0,00 |
| Gnb4 (ENSMUSG00000027669) | 194,52 | -1,15 | 0,33 | -3,49 | 0,00 | 0,01 |
| Abcb4 (ENSMUSG00000042476) | 100,33 | -1,15 | 0,40 | -2,89 | 0,00 | 0,03 |
| Cbx8 (ENSMUSG00000025578) | 184,01 | -1,14 | 0,35 | -3,25 | 0,00 | 0,01 |

| | | | | | | |
|---|---|---|---|---|---|---|
| H1f0 (ENSMUSG00000096210) | 685,29 | -1,14 | 0,23 | -4,97 | 0,00 | 0,00 |
| Foxd2os (ENSMUSG00000085399) | 115,28 | -1,14 | 0,29 | -3,94 | 0,00 | 0,00 |
| D630039A03Rik (ENSMUSG00000052 | 466,22 | -1,14 | 0,30 | -3,79 | 0,00 | 0,00 |
| Zfp367 (ENSMUSG00000044934) | 354,17 | -1,14 | 0,35 | -3,26 | 0,00 | 0,01 |
| Fanci (ENSMUSG00000039187) | 739,29 | -1,14 | 0,34 | -3,31 | 0,00 | 0,01 |
| Ikzf2 (ENSMUSG00000025997) | 211,11 | -1,14 | 0,33 | -3,48 | 0,00 | 0,01 |
| Chaf1b (ENSMUSG00000022945) | 1509,69 | -1,14 | 0,21 | -5,45 | 0,00 | 0,00 |
| Tmem154 (ENSMUSG00000056498) | 1150,32 | -1,13 | 0,24 | -4,77 | 0,00 | 0,00 |
| Chek1 (ENSMUSG00000032113) | 416,58 | -1,13 | 0,31 | -3,68 | 0,00 | 0,00 |
| Crabp2 (ENSMUSG00000004885) | 464,13 | -1,12 | 0,29 | -3,86 | 0,00 | 0,00 |
| Tnfrsf21 (ENSMUSG00000023915) | 235,32 | -1,12 | 0,34 | -3,33 | 0,00 | 0,01 |
| Wdr76 (ENSMUSG00000027242) | 165,19 | -1,12 | 0,40 | -2,82 | 0,00 | 0,04 |
| Gstm4 (ENSMUSG00000027890) | 178,73 | -1,12 | 0,36 | -3,14 | 0,00 | 0,02 |
| Myo1e (ENSMUSG00000032220) | 464,72 | -1,11 | 0,36 | -3,04 | 0,00 | 0,02 |
| Litaf (ENSMUSG00000022500) | 4052,35 | -1,11 | 0,15 | -7,25 | 0,00 | 0,00 |
| H2afz (ENSMUSG00000037894) | 10660,19 | -1,11 | 0,17 | -6,33 | 0,00 | 0,00 |
| Cd74 (ENSMUSG00000024610) | 739,45 | -1,10 | 0,37 | -2,95 | 0,00 | 0,03 |
| 2610318N02Rik (ENSMUSG0000004[illegible] | 363,80 | -1,10 | 0,24 | -4,56 | 0,00 | 0,00 |
| Cbx5 (ENSMUSG00000009575) | 824,84 | -1,09 | 0,12 | -8,77 | 0,00 | 0,00 |
| Mt1 (ENSMUSG00000031765) | 1875,00 | -1,08 | 0,23 | -4,68 | 0,00 | 0,00 |
| Gm43042 (ENSMUSG00000106634) | 195,82 | -1,08 | 0,37 | -2,95 | 0,00 | 0,03 |
| Prim1 (ENSMUSG00000025395) | 1251,67 | -1,08 | 0,20 | -5,48 | 0,00 | 0,00 |
| 2810429I04Rik (ENSMUSG000000977 | 82,73 | -1,08 | 0,36 | -3,01 | 0,00 | 0,03 |
| Nqo2 (ENSMUSG00000046949) | 420,17 | -1,07 | 0,24 | -4,47 | 0,00 | 0,00 |
| Lgalsl (ENSMUSG00000042363) | 231,66 | -1,07 | 0,36 | -2,98 | 0,00 | 0,03 |
| Gm45716 (ENSMUSG00000110344) | 787,64 | -1,07 | 0,26 | -4,17 | 0,00 | 0,00 |
| Gm43041 (ENSMUSG00000104563) | 379,84 | -1,06 | 0,19 | -5,72 | 0,00 | 0,00 |
| Vwa5a (ENSMUSG00000023186) | 899,86 | -1,06 | 0,22 | -4,77 | 0,00 | 0,00 |
| Nsd2 (ENSMUSG00000057406) | 1868,06 | -1,05 | 0,20 | -5,28 | 0,00 | 0,00 |
| Sord (ENSMUSG00000027227) | 238,32 | -1,05 | 0,34 | -3,07 | 0,00 | 0,02 |
| Slc25a10 (ENSMUSG00000025792) | 552,11 | -1,05 | 0,30 | -3,44 | 0,00 | 0,01 |
| Slc14a1 (ENSMUSG00000059336) | 640,33 | -1,05 | 0,21 | -4,89 | 0,00 | 0,00 |
| St14 (ENSMUSG00000031995) | 2114,55 | -1,04 | 0,17 | -5,98 | 0,00 | 0,00 |
| Prr5l (ENSMUSG00000032841) | 248,43 | -1,04 | 0,36 | -2,90 | 0,00 | 0,03 |
| Tmem107 (ENSMUSG00000020895) | 204,88 | -1,03 | 0,18 | -5,78 | 0,00 | 0,00 |
| Pidd1 (ENSMUSG00000025507) | 295,93 | -1,03 | 0,30 | -3,44 | 0,00 | 0,01 |
| 4930579G24Rik (ENSMUSG0000002[illegible] | 487,27 | -1,03 | 0,20 | -5,10 | 0,00 | 0,00 |
| Dnmt1 (ENSMUSG00000004099) | 6106,54 | -1,03 | 0,19 | -5,43 | 0,00 | 0,00 |
| Cdk2 (ENSMUSG00000025358) | 983,62 | -1,03 | 0,20 | -5,04 | 0,00 | 0,00 |
| Tfdp2 (ENSMUSG00000032411) | 470,50 | -1,03 | 0,32 | -3,22 | 0,00 | 0,01 |
| Decr1 (ENSMUSG00000028223) | 501,82 | -1,02 | 0,25 | -4,10 | 0,00 | 0,00 |
| Ifi30 (ENSMUSG00000031838) | 289,77 | -1,02 | 0,27 | -3,83 | 0,00 | 0,00 |
| Pold1 (ENSMUSG00000038644) | 4316,06 | -1,02 | 0,15 | -6,92 | 0,00 | 0,00 |
| Rnaseh2b (ENSMUSG00000021932) | 923,27 | -1,01 | 0,16 | -6,39 | 0,00 | 0,00 |
| Ndc1 (ENSMUSG00000028614) | 991,09 | -1,01 | 0,18 | -5,74 | 0,00 | 0,00 |
| Enpp1 (ENSMUSG00000037370) | 688,66 | -1,01 | 0,28 | -3,61 | 0,00 | 0,00 |
| Grn (ENSMUSG00000034708) | 1396,56 | -1,00 | 0,13 | -7,61 | 0,00 | 0,00 |
| Dynlt3 (ENSMUSG00000031176) | 489,01 | -1,00 | 0,22 | -4,47 | 0,00 | 0,00 |
| Cd200 (ENSMUSG00000022661) | 1060,76 | -1,00 | 0,28 | -3,55 | 0,00 | 0,01 |
| Atad2 (ENSMUSG00000022360) | 2194,44 | -0,99 | 0,27 | -3,63 | 0,00 | 0,00 |
| Cdca7 (ENSMUSG00000055612) | 2664,44 | -0,99 | 0,19 | -5,24 | 0,00 | 0,00 |
| Pmf1 (ENSMUSG00000028066) | 805,64 | -0,99 | 0,18 | -5,54 | 0,00 | 0,00 |
| Gm42031 (ENSMUSG00000110386) | 1042,71 | -0,99 | 0,19 | -5,27 | 0,00 | 0,00 |
| H2afv (ENSMUSG00000041126) | 2129,31 | -0,99 | 0,17 | -5,79 | 0,00 | 0,00 |
| Smc4 (ENSMUSG00000034349) | 6466,37 | -0,98 | 0,16 | -6,32 | 0,00 | 0,00 |
| Rab3d (ENSMUSG00000019066) | 494,50 | -0,98 | 0,23 | -4,33 | 0,00 | 0,00 |
| Hmgn5 (ENSMUSG00000031245) | 489,58 | -0,98 | 0,15 | -6,34 | 0,00 | 0,00 |

| | | | | | | |
|---|---|---|---|---|---|---|
| Gm49980 (ENSMUSG00000117465) | 423,74 | -0,97 | 0,21 | -4,65 | 0,00 | 0,00 |
| Serpina3g (ENSMUSG00000041481) | 6776,11 | -0,97 | 0,17 | -5,63 | 0,00 | 0,00 |
| Atg9a (ENSMUSG00000033124) | 527,16 | -0,97 | 0,31 | -3,14 | 0,00 | 0,02 |
| Nup37 (ENSMUSG00000035351) | 513,94 | -0,97 | 0,26 | -3,66 | 0,00 | 0,00 |
| Gins3 (ENSMUSG00000031669) | 328,52 | -0,96 | 0,35 | -2,78 | 0,01 | 0,04 |
| Tmem38b (ENSMUSG00000028420) | 152,06 | -0,96 | 0,30 | -3,20 | 0,00 | 0,01 |
| Ccne1 (ENSMUSG00000002068) | 943,59 | -0,95 | 0,21 | -4,61 | 0,00 | 0,00 |
| Il3ra (ENSMUSG00000068758) | 191,57 | -0,95 | 0,27 | -3,53 | 0,00 | 0,01 |
| Rpa2 (ENSMUSG00000028884) | 2886,08 | -0,95 | 0,14 | -6,98 | 0,00 | 0,00 |
| Gm48275 (ENSMUSG00000111202) | 90,78 | -0,95 | 0,30 | -3,14 | 0,00 | 0,02 |
| Rmnd1 (ENSMUSG00000019763) | 332,21 | -0,95 | 0,27 | -3,52 | 0,00 | 0,01 |
| Casp3 (ENSMUSG00000031628) | 6545,78 | -0,95 | 0,14 | -6,95 | 0,00 | 0,00 |
| Gm26527 (ENSMUSG00000097582) | 258,04 | -0,94 | 0,22 | -4,33 | 0,00 | 0,00 |
| Uxs1 (ENSMUSG00000057363) | 228,63 | -0,94 | 0,26 | -3,68 | 0,00 | 0,00 |
| Tspan32 (ENSMUSG00000000244) | 1179,76 | -0,94 | 0,19 | -4,98 | 0,00 | 0,00 |
| Psmc3ip (ENSMUSG00000019303) | 279,22 | -0,94 | 0,24 | -3,91 | 0,00 | 0,00 |
| C130026I21Rik (ENSMUSG000000524 | 49,96 | -0,94 | 0,33 | -2,85 | 0,00 | 0,04 |
| Mbnl3 (ENSMUSG00000036109) | 411,75 | -0,94 | 0,18 | -5,31 | 0,00 | 0,00 |
| Pctp (ENSMUSG00000020553) | 192,71 | -0,93 | 0,23 | -4,09 | 0,00 | 0,00 |
| Pkig (ENSMUSG00000035268) | 343,44 | -0,93 | 0,13 | -7,14 | 0,00 | 0,00 |
| Axl (ENSMUSG00000002602) | 318,28 | -0,93 | 0,27 | -3,51 | 0,00 | 0,01 |
| Mgst2 (ENSMUSG00000074604) | 613,42 | -0,93 | 0,17 | -5,42 | 0,00 | 0,00 |
| Apip (ENSMUSG00000010911) | 233,81 | -0,93 | 0,23 | -4,09 | 0,00 | 0,00 |
| Sema4c (ENSMUSG00000026121) | 269,27 | -0,93 | 0,29 | -3,24 | 0,00 | 0,01 |
| Lrrc40 (ENSMUSG00000063052) | 758,92 | -0,92 | 0,18 | -5,12 | 0,00 | 0,00 |
| Tube1 (ENSMUSG00000019845) | 204,89 | -0,92 | 0,28 | -3,32 | 0,00 | 0,01 |
| Wee1 (ENSMUSG00000031016) | 508,33 | -0,92 | 0,23 | -4,04 | 0,00 | 0,00 |
| Pdcd1 (ENSMUSG00000026285) | 6991,90 | -0,92 | 0,21 | -4,40 | 0,00 | 0,00 |
| Rfc1 (ENSMUSG00000029191) | 2047,88 | -0,92 | 0,17 | -5,43 | 0,00 | 0,00 |
| Mdm1 (ENSMUSG00000020212) | 306,99 | -0,92 | 0,27 | -3,45 | 0,00 | 0,01 |
| Pcyox1l (ENSMUSG00000024579) | 238,87 | -0,91 | 0,22 | -4,05 | 0,00 | 0,00 |
| Trdmt1 (ENSMUSG00000026723) | 184,45 | -0,90 | 0,27 | -3,29 | 0,00 | 0,01 |
| Cmc2 (ENSMUSG00000014633) | 459,51 | -0,90 | 0,16 | -5,70 | 0,00 | 0,00 |
| Cep57l1 (ENSMUSG00000019813) | 352,74 | -0,90 | 0,22 | -4,12 | 0,00 | 0,00 |
| Tfdp1 (ENSMUSG00000038482) | 4880,04 | -0,89 | 0,13 | -6,82 | 0,00 | 0,00 |
| Chst12 (ENSMUSG00000036599) | 1396,07 | -0,89 | 0,15 | -5,94 | 0,00 | 0,00 |
| H2afx (ENSMUSG00000049932) | 2448,99 | -0,89 | 0,14 | -6,26 | 0,00 | 0,00 |
| Eci2 (ENSMUSG00000021417) | 1544,52 | -0,89 | 0,17 | -5,20 | 0,00 | 0,00 |
| Dipk1b (ENSMUSG00000036186) | 220,59 | -0,89 | 0,26 | -3,44 | 0,00 | 0,01 |
| Pglyrp1 (ENSMUSG00000030413) | 1437,86 | -0,88 | 0,18 | -4,98 | 0,00 | 0,00 |
| Sh3bgrl (ENSMUSG00000031246) | 1821,05 | -0,88 | 0,15 | -5,99 | 0,00 | 0,00 |
| 9030025P20Rik (ENSMUSG00000116 | 130,72 | -0,88 | 0,32 | -2,76 | 0,01 | 0,05 |
| Oxct1 (ENSMUSG00000022186) | 2957,52 | -0,88 | 0,13 | -6,86 | 0,00 | 0,00 |
| Cenpu (ENSMUSG00000031629) | 255,13 | -0,88 | 0,25 | -3,55 | 0,00 | 0,01 |
| Txndc16 (ENSMUSG00000021830) | 557,28 | -0,88 | 0,22 | -3,97 | 0,00 | 0,00 |
| Eogt (ENSMUSG00000035245) | 365,43 | -0,88 | 0,23 | -3,84 | 0,00 | 0,00 |
| BC035044 (ENSMUSG00000090164) | 349,49 | -0,87 | 0,31 | -2,80 | 0,01 | 0,04 |
| Capg (ENSMUSG00000056737) | 4082,24 | -0,87 | 0,18 | -4,95 | 0,00 | 0,00 |
| BC052040 (ENSMUSG00000040282) | 310,17 | -0,87 | 0,31 | -2,86 | 0,00 | 0,04 |
| Gm46223 (ENSMUSG00000110720) | 140,36 | -0,87 | 0,25 | -3,54 | 0,00 | 0,01 |
| Mcm6 (ENSMUSG00000026355) | 18502,68 | -0,87 | 0,11 | -7,70 | 0,00 | 0,00 |
| Kpna2 (ENSMUSG00000018362) | 4163,40 | -0,87 | 0,19 | -4,58 | 0,00 | 0,00 |
| Tubb5 (ENSMUSG00000001525) | 11926,02 | -0,87 | 0,16 | -5,28 | 0,00 | 0,00 |
| Gm10075 (ENSMUSG00000100720) | 60,63 | -0,86 | 0,23 | -3,69 | 0,00 | 0,00 |
| B3glct (ENSMUSG00000051950) | 296,34 | -0,86 | 0,23 | -3,78 | 0,00 | 0,00 |
| Cenpj (ENSMUSG00000064128) | 301,27 | -0,86 | 0,28 | -3,12 | 0,00 | 0,02 |
| Ezh2 (ENSMUSG00000029687) | 2024,01 | -0,86 | 0,16 | -5,30 | 0,00 | 0,00 |

| | | | | | | |
|---|---|---|---|---|---|---|
| Khk (ENSMUSG00000029162) | 191,61 | -0,85 | 0,30 | -2,88 | 0,00 | 0,03 |
| Filip1l (ENSMUSG00000043336) | 842,37 | -0,85 | 0,20 | -4,21 | 0,00 | 0,00 |
| Kdelc2 (ENSMUSG00000034487) | 385,56 | -0,85 | 0,31 | -2,75 | 0,01 | 0,05 |
| Csrp1 (ENSMUSG00000026421) | 2977,15 | -0,85 | 0,14 | -6,00 | 0,00 | 0,00 |
| 0610009B22Rik (ENSMUSG0000007[illegible] | 123,81 | -0,85 | 0,31 | -2,78 | 0,01 | 0,04 |
| Ephx1 (ENSMUSG00000038776) | 2590,69 | -0,85 | 0,23 | -3,74 | 0,00 | 0,00 |
| Pcna (ENSMUSG00000027342) | 5916,70 | -0,85 | 0,18 | -4,63 | 0,00 | 0,00 |
| Abcb8 (ENSMUSG00000028973) | 894,27 | -0,84 | 0,18 | -4,66 | 0,00 | 0,00 |
| Arl14ep (ENSMUSG00000027122) | 600,41 | -0,84 | 0,21 | -4,06 | 0,00 | 0,00 |
| Gm17018 (ENSMUSG00000041035) | 722,24 | -0,84 | 0,18 | -4,76 | 0,00 | 0,00 |
| Lpgat1 (ENSMUSG00000026623) | 1270,23 | -0,84 | 0,19 | -4,47 | 0,00 | 0,00 |
| Tmco6 (ENSMUSG00000006850) | 292,30 | -0,84 | 0,28 | -3,05 | 0,00 | 0,02 |
| Fen1 (ENSMUSG00000024742) | 3906,41 | -0,84 | 0,12 | -7,11 | 0,00 | 0,00 |
| H2-Oa (ENSMUSG00000024334) | 462,26 | -0,82 | 0,17 | -4,79 | 0,00 | 0,00 |
| Emc8 (ENSMUSG00000031819) | 1055,17 | -0,82 | 0,12 | -7,13 | 0,00 | 0,00 |
| Mcub (ENSMUSG00000027994) | 352,20 | -0,82 | 0,29 | -2,84 | 0,00 | 0,04 |
| Rfc2 (ENSMUSG00000023104) | 3889,72 | -0,82 | 0,18 | -4,55 | 0,00 | 0,00 |
| Susd1 (ENSMUSG00000038578) | 222,74 | -0,81 | 0,20 | -4,03 | 0,00 | 0,00 |
| Tpst1 (ENSMUSG00000034118) | 424,92 | -0,81 | 0,29 | -2,81 | 0,00 | 0,04 |
| Zbed3 (ENSMUSG00000041995) | 342,22 | -0,81 | 0,21 | -3,87 | 0,00 | 0,00 |
| Arsb (ENSMUSG00000042082) | 4247,68 | -0,81 | 0,12 | -6,80 | 0,00 | 0,00 |
| Pck2 (ENSMUSG00000040618) | 593,81 | -0,80 | 0,20 | -4,12 | 0,00 | 0,00 |
| Cdc25b (ENSMUSG00000027330) | 2675,46 | -0,80 | 0,22 | -3,59 | 0,00 | 0,00 |
| Atpif1 (ENSMUSG00000054428) | 1535,33 | -0,80 | 0,16 | -4,98 | 0,00 | 0,00 |
| Marveld2 (ENSMUSG00000021636) | 368,31 | -0,80 | 0,25 | -3,21 | 0,00 | 0,01 |
| Rasgef1b (ENSMUSG00000089809) | 820,10 | -0,80 | 0,14 | -5,82 | 0,00 | 0,00 |
| Alg6 (ENSMUSG00000073792) | 234,86 | -0,80 | 0,28 | -2,85 | 0,00 | 0,04 |
| Acat2 (ENSMUSG00000023832) | 553,42 | -0,79 | 0,27 | -2,93 | 0,00 | 0,03 |
| Cisd1 (ENSMUSG00000037710) | 1530,55 | -0,79 | 0,18 | -4,35 | 0,00 | 0,00 |
| Serpina3f (ENSMUSG00000066363) | 1230,95 | -0,79 | 0,25 | -3,20 | 0,00 | 0,01 |
| AC158975.2 (ENSMUSG00000118012[illegible] | 3046,62 | -0,79 | 0,16 | -4,94 | 0,00 | 0,00 |
| Slco4a1 (ENSMUSG00000038963) | 1944,98 | -0,79 | 0,15 | -5,19 | 0,00 | 0,00 |
| Scpep1 (ENSMUSG00000000278) | 1003,96 | -0,79 | 0,15 | -5,19 | 0,00 | 0,00 |
| Tmem254c (ENSMUSG00000072680) | 641,72 | -0,79 | 0,22 | -3,54 | 0,00 | 0,01 |
| Plscr1 (ENSMUSG00000032369) | 2071,02 | -0,79 | 0,15 | -5,15 | 0,00 | 0,00 |
| Lgals3 (ENSMUSG00000050335) | 2834,34 | -0,78 | 0,24 | -3,24 | 0,00 | 0,01 |
| Cenpo (ENSMUSG00000020652) | 539,43 | -0,78 | 0,23 | -3,35 | 0,00 | 0,01 |
| 2010315B03Rik (ENSMUSG0000007[illegible] | 168,42 | -0,78 | 0,26 | -3,04 | 0,00 | 0,02 |
| Cbr4 (ENSMUSG00000031641) | 255,16 | -0,78 | 0,23 | -3,44 | 0,00 | 0,01 |
| Siva1 (ENSMUSG00000064326) | 1241,73 | -0,78 | 0,21 | -3,62 | 0,00 | 0,00 |
| Fgl2 (ENSMUSG00000039899) | 4340,13 | -0,78 | 0,13 | -5,91 | 0,00 | 0,00 |
| Plpp5 (ENSMUSG00000031570) | 254,36 | -0,78 | 0,28 | -2,75 | 0,01 | 0,05 |
| A430093F15Rik (ENSMUSG0000006[illegible] | 550,29 | -0,78 | 0,20 | -3,79 | 0,00 | 0,00 |
| Stx7 (ENSMUSG00000019998) | 581,83 | -0,77 | 0,14 | -5,56 | 0,00 | 0,00 |
| Tubg1 (ENSMUSG00000035198) | 1914,98 | -0,77 | 0,15 | -5,25 | 0,00 | 0,00 |
| Ccpg1os (ENSMUSG00000086158) | 42,01 | -0,77 | 0,27 | -2,87 | 0,00 | 0,04 |
| Cks2 (ENSMUSG00000062248) | 837,77 | -0,77 | 0,21 | -3,76 | 0,00 | 0,00 |
| Cd72 (ENSMUSG00000028459) | 1370,03 | -0,77 | 0,21 | -3,62 | 0,00 | 0,00 |
| Klhl22 (ENSMUSG00000022750) | 642,29 | -0,77 | 0,26 | -3,00 | 0,00 | 0,03 |
| Akip1 (ENSMUSG00000031023) | 625,27 | -0,77 | 0,18 | -4,38 | 0,00 | 0,00 |
| Slc25a24 (ENSMUSG00000040322) | 1035,02 | -0,77 | 0,14 | -5,34 | 0,00 | 0,00 |
| Ccdc28a (ENSMUSG00000059554) | 173,57 | -0,77 | 0,22 | -3,46 | 0,00 | 0,01 |
| Phf11a (ENSMUSG00000044703) | 556,27 | -0,77 | 0,19 | -4,02 | 0,00 | 0,00 |
| Tmem237 (ENSMUSG00000038079) | 337,82 | -0,77 | 0,16 | -4,68 | 0,00 | 0,00 |
| Cfap20 (ENSMUSG00000031796) | 794,30 | -0,77 | 0,16 | -4,72 | 0,00 | 0,00 |
| Gm15452 (ENSMUSG00000084353) | 189,99 | -0,77 | 0,27 | -2,82 | 0,00 | 0,04 |
| Hjurp (ENSMUSG00000044783) | 2410,37 | -0,77 | 0,15 | -5,13 | 0,00 | 0,00 |

| | | | | | | |
|---|---|---|---|---|---|---|
| Ckap5 (ENSMUSG00000040549) | 2186,89 | -0,77 | 0,18 | -4,28 | 0,00 | 0,00 |
| Ehd4 (ENSMUSG00000027293) | 1970,73 | -0,77 | 0,15 | -5,25 | 0,00 | 0,00 |
| Nup155 (ENSMUSG00000022142) | 1070,67 | -0,76 | 0,23 | -3,27 | 0,00 | 0,01 |
| Mmd (ENSMUSG00000003948) | 1778,34 | -0,76 | 0,21 | -3,70 | 0,00 | 0,00 |
| Lyrm9 (ENSMUSG00000072640) | 301,38 | -0,76 | 0,25 | -3,09 | 0,00 | 0,02 |
| Nde1 (ENSMUSG00000022678) | 1952,88 | -0,76 | 0,13 | -5,82 | 0,00 | 0,00 |
| Cldn10 (ENSMUSG00000022132) | 171,35 | -0,76 | 0,23 | -3,31 | 0,00 | 0,01 |
| Tmpo (ENSMUSG00000019961) | 7569,73 | -0,76 | 0,10 | -7,50 | 0,00 | 0,00 |
| Gm18194 (ENSMUSG00000109724) | 190,67 | -0,76 | 0,21 | -3,52 | 0,00 | 0,01 |
| Gm10143 (ENSMUSG00000064032) | 369,41 | -0,75 | 0,24 | -3,14 | 0,00 | 0,02 |
| C2cd5 (ENSMUSG00000030279) | 1182,17 | -0,75 | 0,24 | -3,20 | 0,00 | 0,01 |
| Tstd3 (ENSMUSG00000028251) | 671,83 | -0,75 | 0,13 | -5,93 | 0,00 | 0,00 |
| Galns (ENSMUSG00000015027) | 983,30 | -0,75 | 0,19 | -4,01 | 0,00 | 0,00 |
| Gstp3 (ENSMUSG00000058216) | 826,71 | -0,74 | 0,17 | -4,39 | 0,00 | 0,00 |
| Gm11273 (ENSMUSG00000079941) | 93,89 | -0,74 | 0,27 | -2,76 | 0,01 | 0,04 |
| Gm49384 (ENSMUSG00000113786) | 252,24 | -0,74 | 0,23 | -3,28 | 0,00 | 0,01 |
| Cep85 (ENSMUSG00000037443) | 1569,05 | -0,74 | 0,13 | -5,58 | 0,00 | 0,00 |
| Haao (ENSMUSG00000000673) | 381,35 | -0,74 | 0,24 | -3,10 | 0,00 | 0,02 |
| Ap1s1 (ENSMUSG00000004849) | 2056,13 | -0,74 | 0,13 | -5,72 | 0,00 | 0,00 |
| Ncapd3 (ENSMUSG00000035024) | 1484,79 | -0,74 | 0,22 | -3,38 | 0,00 | 0,01 |
| Hmga1 (ENSMUSG00000046711) | 1822,43 | -0,73 | 0,14 | -5,17 | 0,00 | 0,00 |
| Fabp5 (ENSMUSG00000027533) | 961,26 | -0,73 | 0,26 | -2,85 | 0,00 | 0,04 |
| Fam3c (ENSMUSG00000029672) | 2172,62 | -0,73 | 0,13 | -5,48 | 0,00 | 0,00 |
| Gm21987 (ENSMUSG00000095464) | 369,39 | -0,73 | 0,23 | -3,23 | 0,00 | 0,01 |
| Dkkl1 (ENSMUSG00000030792) | 344,73 | -0,73 | 0,17 | -4,37 | 0,00 | 0,00 |
| Gm42427 (ENSMUSG00000105263) | 2188,66 | -0,72 | 0,11 | -6,90 | 0,00 | 0,00 |
| Mindy3 (ENSMUSG00000026767) | 662,14 | -0,72 | 0,21 | -3,46 | 0,00 | 0,01 |
| Gemin6 (ENSMUSG00000055760) | 477,15 | -0,72 | 0,22 | -3,36 | 0,00 | 0,01 |
| Suv39h1 (ENSMUSG00000039231) | 1546,20 | -0,72 | 0,16 | -4,54 | 0,00 | 0,00 |
| Snx13 (ENSMUSG00000020590) | 464,11 | -0,72 | 0,23 | -3,13 | 0,00 | 0,02 |
| Tceal9 (ENSMUSG00000042712) | 891,85 | -0,72 | 0,12 | -5,97 | 0,00 | 0,00 |
| 2310031A07Rik (ENSMUSG0000009 | 318,59 | -0,72 | 0,21 | -3,37 | 0,00 | 0,01 |
| Sephs1 (ENSMUSG00000026662) | 1687,41 | -0,72 | 0,12 | -5,78 | 0,00 | 0,00 |
| Paqr4 (ENSMUSG00000023909) | 1242,38 | -0,72 | 0,23 | -3,13 | 0,00 | 0,02 |
| Haus5 (ENSMUSG00000078762) | 832,00 | -0,71 | 0,24 | -3,03 | 0,00 | 0,02 |
| Cinp (ENSMUSG00000021276) | 1257,89 | -0,71 | 0,14 | -5,09 | 0,00 | 0,00 |
| Cdk5rap1 (ENSMUSG00000027487) | 526,93 | -0,71 | 0,21 | -3,40 | 0,00 | 0,01 |
| Ccdc181 (ENSMUSG00000026578) | 267,42 | -0,71 | 0,24 | -3,02 | 0,00 | 0,02 |
| Coq7 (ENSMUSG00000030652) | 407,02 | -0,71 | 0,24 | -3,01 | 0,00 | 0,02 |
| Dynlt1b (ENSMUSG00000096255) | 786,25 | -0,71 | 0,18 | -3,95 | 0,00 | 0,00 |
| Dbi (ENSMUSG00000026385) | 1838,11 | -0,71 | 0,20 | -3,52 | 0,00 | 0,01 |
| Psen2 (ENSMUSG00000010609) | 1532,48 | -0,71 | 0,18 | -3,88 | 0,00 | 0,00 |
| Pfas (ENSMUSG00000020899) | 1128,22 | -0,71 | 0,25 | -2,86 | 0,00 | 0,04 |
| Cyfip1 (ENSMUSG00000030447) | 900,00 | -0,71 | 0,15 | -4,62 | 0,00 | 0,00 |
| Reep4 (ENSMUSG00000033589) | 1030,02 | -0,70 | 0,12 | -5,63 | 0,00 | 0,00 |
| Nudt1 (ENSMUSG00000036639) | 582,49 | -0,70 | 0,15 | -4,70 | 0,00 | 0,00 |
| Ripk3 (ENSMUSG00000022221) | 1501,79 | -0,70 | 0,22 | -3,24 | 0,00 | 0,01 |
| Twsg1 (ENSMUSG00000024098) | 3752,00 | -0,70 | 0,12 | -5,77 | 0,00 | 0,00 |
| Stoml1 (ENSMUSG00000032333) | 400,32 | -0,70 | 0,22 | -3,17 | 0,00 | 0,02 |
| Itgb3bp (ENSMUSG00000028549) | 296,09 | -0,70 | 0,19 | -3,59 | 0,00 | 0,00 |
| Psat1 (ENSMUSG00000024640) | 2910,81 | -0,70 | 0,21 | -3,33 | 0,00 | 0,01 |
| Haus1 (ENSMUSG00000041840) | 522,08 | -0,70 | 0,17 | -4,14 | 0,00 | 0,00 |
| Dynlt1-ps1 (ENSMUSG00000082691) | 359,75 | -0,70 | 0,22 | -3,15 | 0,00 | 0,02 |
| Nkap (ENSMUSG00000016409) | 602,80 | -0,70 | 0,19 | -3,68 | 0,00 | 0,00 |
| Tssc4 (ENSMUSG00000045752) | 602,77 | -0,69 | 0,23 | -3,07 | 0,00 | 0,02 |
| Ccnh (ENSMUSG00000021548) | 1746,42 | -0,69 | 0,15 | -4,73 | 0,00 | 0,00 |
| Nsmce1 (ENSMUSG00000030750) | 1338,12 | -0,69 | 0,21 | -3,33 | 0,00 | 0,01 |

| | | | | | | |
|---|---|---|---|---|---|---|
| Smyd1 (ENSMUSG00000055027) | 220,95 | -0,69 | 0,21 | -3,27 | 0,00 | 0,01 |
| Haghl (ENSMUSG00000061046) | 301,38 | -0,69 | 0,20 | -3,42 | 0,00 | 0,01 |
| Kmt5c (ENSMUSG00000059851) | 481,36 | -0,69 | 0,18 | -3,77 | 0,00 | 0,00 |
| Pola2 (ENSMUSG00000024833) | 2403,90 | -0,69 | 0,13 | -5,24 | 0,00 | 0,00 |
| Tacc2 (ENSMUSG00000030852) | 719,78 | -0,68 | 0,21 | -3,32 | 0,00 | 0,01 |
| Gm49804 (ENSMUSG00000117338) | 138,45 | -0,68 | 0,22 | -3,17 | 0,00 | 0,02 |
| Ighm (ENSMUSG00000076617) | 9293,91 | -0,68 | 0,20 | -3,43 | 0,00 | 0,01 |
| Ipp (ENSMUSG00000028696) | 506,60 | -0,68 | 0,19 | -3,49 | 0,00 | 0,01 |
| Pafah1b3 (ENSMUSG00000005447) | 528,39 | -0,68 | 0,15 | -4,42 | 0,00 | 0,00 |
| Serinc3 (ENSMUSG00000017707) | 18496,99 | -0,68 | 0,11 | -6,04 | 0,00 | 0,00 |
| Polh (ENSMUSG00000023953) | 502,22 | -0,68 | 0,23 | -3,01 | 0,00 | 0,03 |
| Usp1 (ENSMUSG00000028560) | 3390,82 | -0,67 | 0,17 | -4,04 | 0,00 | 0,00 |
| Slamf6 (ENSMUSG00000015314) | 2998,60 | -0,67 | 0,24 | -2,74 | 0,01 | 0,05 |
| Mtch1 (ENSMUSG00000024012) | 2798,96 | -0,67 | 0,15 | -4,58 | 0,00 | 0,00 |
| Tmcc1 (ENSMUSG00000030126) | 516,78 | -0,67 | 0,18 | -3,74 | 0,00 | 0,00 |
| Msl3 (ENSMUSG00000031358) | 972,05 | -0,66 | 0,12 | -5,57 | 0,00 | 0,00 |
| Fdps (ENSMUSG00000059743) | 964,77 | -0,66 | 0,23 | -2,88 | 0,00 | 0,03 |
| Pdlim7 (ENSMUSG00000021493) | 336,03 | -0,66 | 0,20 | -3,35 | 0,00 | 0,01 |
| Tyw5 (ENSMUSG00000048495) | 237,60 | -0,66 | 0,24 | -2,73 | 0,01 | 0,05 |
| Tuba1b (ENSMUSG00000023004) | 40383,49 | -0,65 | 0,15 | -4,33 | 0,00 | 0,00 |
| Bspry (ENSMUSG00000028392) | 919,06 | -0,65 | 0,22 | -2,98 | 0,00 | 0,03 |
| S100a6 (ENSMUSG00000001025) | 7999,47 | -0,65 | 0,15 | -4,40 | 0,00 | 0,00 |
| Cdt1 (ENSMUSG00000006585) | 1488,43 | -0,65 | 0,15 | -4,34 | 0,00 | 0,00 |
| St7 (ENSMUSG00000029534) | 513,68 | -0,65 | 0,17 | -3,94 | 0,00 | 0,00 |
| Prelid2 (ENSMUSG00000056671) | 903,97 | -0,65 | 0,17 | -3,93 | 0,00 | 0,00 |
| Dynlt1f (ENSMUSG00000095677) | 928,59 | -0,65 | 0,19 | -3,50 | 0,00 | 0,01 |
| Anxa2 (ENSMUSG00000032231) | 27646,27 | -0,65 | 0,14 | -4,72 | 0,00 | 0,00 |
| Ccsap (ENSMUSG00000031971) | 479,35 | -0,65 | 0,21 | -3,06 | 0,00 | 0,02 |
| Hmox2 (ENSMUSG00000004070) | 2824,95 | -0,65 | 0,15 | -4,20 | 0,00 | 0,00 |
| Rpa1 (ENSMUSG00000000751) | 5259,65 | -0,64 | 0,09 | -7,08 | 0,00 | 0,00 |
| Ubash3b (ENSMUSG00000032020) | 5949,07 | -0,64 | 0,07 | -8,76 | 0,00 | 0,00 |
| Hmgb3 (ENSMUSG00000015217) | 1313,70 | -0,64 | 0,17 | -3,73 | 0,00 | 0,00 |
| Mlkl (ENSMUSG00000012519) | 535,71 | -0,64 | 0,15 | -4,35 | 0,00 | 0,00 |
| Pradc1 (ENSMUSG00000030008) | 588,63 | -0,64 | 0,23 | -2,80 | 0,01 | 0,04 |
| Cdca7l (ENSMUSG00000021175) | 2620,82 | -0,64 | 0,13 | -4,85 | 0,00 | 0,00 |
| Psph (ENSMUSG00000029446) | 1406,85 | -0,64 | 0,16 | -4,09 | 0,00 | 0,00 |
| Rbmx (ENSMUSG00000031134) | 1101,75 | -0,64 | 0,14 | -4,74 | 0,00 | 0,00 |
| Ppih (ENSMUSG00000060288) | 747,64 | -0,64 | 0,16 | -3,94 | 0,00 | 0,00 |
| Plp2 (ENSMUSG00000031146) | 3320,02 | -0,64 | 0,15 | -4,33 | 0,00 | 0,00 |
| Eef1akmt1 (ENSMUSG00000021951) | 1018,50 | -0,64 | 0,19 | -3,35 | 0,00 | 0,01 |
| Hmgb1 (ENSMUSG00000066551) | 12866,31 | -0,64 | 0,12 | -5,36 | 0,00 | 0,00 |
| Cdk6 (ENSMUSG00000040274) | 1383,60 | -0,63 | 0,15 | -4,23 | 0,00 | 0,00 |
| Taf5 (ENSMUSG00000025049) | 388,27 | -0,63 | 0,21 | -2,96 | 0,00 | 0,03 |
| Rnf26 (ENSMUSG00000053128) | 2353,76 | -0,63 | 0,18 | -3,48 | 0,00 | 0,01 |
| Pycard (ENSMUSG00000030793) | 2165,92 | -0,63 | 0,21 | -2,99 | 0,00 | 0,03 |
| Slc37a4 (ENSMUSG00000032114) | 646,78 | -0,63 | 0,21 | -2,98 | 0,00 | 0,03 |
| Bola1 (ENSMUSG00000015943) | 237,12 | -0,63 | 0,21 | -3,08 | 0,00 | 0,02 |
| Brip1os (ENSMUSG00000085208) | 1013,11 | -0,63 | 0,18 | -3,53 | 0,00 | 0,01 |
| Nup107 (ENSMUSG00000052798) | 1915,41 | -0,63 | 0,11 | -5,57 | 0,00 | 0,00 |
| Ttf2 (ENSMUSG00000033222) | 900,00 | -0,63 | 0,23 | -2,74 | 0,01 | 0,05 |
| Crip1 (ENSMUSG00000006360) | 8353,27 | -0,63 | 0,14 | -4,37 | 0,00 | 0,00 |
| Ifi27 (ENSMUSG00000064215) | 2764,98 | -0,63 | 0,10 | -6,23 | 0,00 | 0,00 |
| Dtymk (ENSMUSG00000026281) | 1371,83 | -0,63 | 0,18 | -3,41 | 0,00 | 0,01 |
| Comt (ENSMUSG00000000326) | 1412,19 | -0,63 | 0,13 | -4,92 | 0,00 | 0,00 |
| Mkrn2 (ENSMUSG00000000439) | 1224,02 | -0,63 | 0,15 | -4,17 | 0,00 | 0,00 |
| Pdk1 (ENSMUSG00000006494) | 2578,08 | -0,62 | 0,12 | -5,16 | 0,00 | 0,00 |
| Fdx1 (ENSMUSG00000032051) | 429,63 | -0,62 | 0,20 | -3,08 | 0,00 | 0,02 |

| | | | | | | |
|---|---|---|---|---|---|---|
| Slbp (ENSMUSG00000004642) | 4022,57 | -0,62 | 0,13 | -4,78 | 0,00 | 0,00 |
| Cep57 (ENSMUSG00000031922) | 962,50 | -0,62 | 0,12 | -5,15 | 0,00 | 0,00 |
| Smpdl3a (ENSMUSG00000019872) | 1918,49 | -0,62 | 0,23 | -2,75 | 0,01 | 0,05 |
| Ankrd9 (ENSMUSG00000037904) | 291,51 | -0,62 | 0,19 | -3,28 | 0,00 | 0,01 |
| Pcbd2 (ENSMUSG00000021496) | 312,05 | -0,62 | 0,20 | -3,08 | 0,00 | 0,02 |
| Serpinb6a (ENSMUSG00000060147) | 1320,79 | -0,62 | 0,18 | -3,38 | 0,00 | 0,01 |
| Rpa3 (ENSMUSG00000012483) | 666,64 | -0,62 | 0,23 | -2,73 | 0,01 | 0,05 |
| Stambpl1 (ENSMUSG00000024776) | 605,90 | -0,62 | 0,20 | -3,06 | 0,00 | 0,02 |
| Adcy7 (ENSMUSG00000031659) | 3700,81 | -0,61 | 0,16 | -3,96 | 0,00 | 0,00 |
| Bcl2l12 (ENSMUSG00000003190) | 271,57 | -0,61 | 0,19 | -3,27 | 0,00 | 0,01 |
| Mpp1 (ENSMUSG00000031402) | 1611,44 | -0,61 | 0,15 | -4,09 | 0,00 | 0,00 |
| Fam92a (ENSMUSG00000028218) | 822,78 | -0,61 | 0,13 | -4,81 | 0,00 | 0,00 |
| Sec22a (ENSMUSG00000034473) | 403,27 | -0,61 | 0,23 | -2,71 | 0,01 | 0,05 |
| Gmfg (ENSMUSG00000060791) | 4321,82 | -0,61 | 0,15 | -4,08 | 0,00 | 0,00 |
| Nup85 (ENSMUSG00000020739) | 2190,56 | -0,61 | 0,14 | -4,26 | 0,00 | 0,00 |
| Vamp8 (ENSMUSG00000050732) | 2273,17 | -0,61 | 0,13 | -4,52 | 0,00 | 0,00 |
| Gm26880 (ENSMUSG00000096979) | 441,51 | -0,61 | 0,16 | -3,69 | 0,00 | 0,00 |
| Dpysl2 (ENSMUSG00000022048) | 838,15 | -0,61 | 0,14 | -4,39 | 0,00 | 0,00 |
| Zdhhc2 (ENSMUSG00000039470) | 776,59 | -0,61 | 0,20 | -3,02 | 0,00 | 0,02 |
| Hcfc2 (ENSMUSG00000020246) | 561,29 | -0,61 | 0,17 | -3,49 | 0,00 | 0,01 |
| Rnft1 (ENSMUSG00000020521) | 776,67 | -0,60 | 0,19 | -3,13 | 0,00 | 0,02 |
| Itga6 (ENSMUSG00000027111) | 1341,44 | -0,60 | 0,18 | -3,31 | 0,00 | 0,01 |
| Xaf1 (ENSMUSG00000040483) | 859,47 | -0,60 | 0,20 | -3,08 | 0,00 | 0,02 |
| Emp3 (ENSMUSG00000040212) | 3081,86 | -0,60 | 0,16 | -3,84 | 0,00 | 0,00 |
| Rad18 (ENSMUSG00000030254) | 1315,31 | -0,60 | 0,12 | -5,00 | 0,00 | 0,00 |
| Pacsin1 (ENSMUSG00000040276) | 1771,91 | -0,60 | 0,17 | -3,48 | 0,00 | 0,01 |
| Mgme1 (ENSMUSG00000027424) | 356,54 | -0,60 | 0,21 | -2,89 | 0,00 | 0,03 |
| Mfsd10 (ENSMUSG00000001082) | 1361,30 | -0,60 | 0,17 | -3,59 | 0,00 | 0,00 |
| Dusp4 (ENSMUSG00000031530) | 3106,57 | -0,59 | 0,20 | -2,98 | 0,00 | 0,03 |
| Ubr7 (ENSMUSG00000041712) | 1553,84 | -0,59 | 0,09 | -6,27 | 0,00 | 0,00 |
| Dera (ENSMUSG00000030225) | 2406,16 | -0,59 | 0,15 | -4,02 | 0,00 | 0,00 |
| Rbl1 (ENSMUSG00000027641) | 1076,18 | -0,59 | 0,10 | -5,81 | 0,00 | 0,00 |
| Topbp1 (ENSMUSG00000032555) | 2550,08 | -0,59 | 0,17 | -3,42 | 0,00 | 0,01 |
| Sumo3 (ENSMUSG00000020265) | 2970,72 | -0,59 | 0,17 | -3,49 | 0,00 | 0,01 |
| Gm9888 (ENSMUSG00000052724) | 110,03 | -0,59 | 0,21 | -2,85 | 0,00 | 0,04 |
| Dnajc9 (ENSMUSG00000021811) | 4884,19 | -0,59 | 0,15 | -3,84 | 0,00 | 0,00 |
